# The role of active mRNA-ribosome dynamics and closing constriction in daughter chromosome separation in *Escherichia coli*

**DOI:** 10.1101/2025.04.05.647368

**Authors:** Chathuddasie I. Amarasinghe, Mu-Hung Chang, Jaana Männik, Scott T. Retterer, Maxim O. Lavrentovich, Jaan Männik

## Abstract

The mechanisms by which two sister chromosomes separate and partition into daughter cells in bacteria remain poorly understood. A recent theoretical model has proposed that out-of-equilibrium central dogma reactions involving mRNA and ribosomes play a significant role in this process. Here we test this idea in the *Escherichia coli* model system using high-throughput fluorescence microscopy in microfluidic devices. We compare our experimental observations with predictions from a reaction-diffusion model that includes central dogma-related reactions and excluded volume interactions between ribosomal subunits, polysomes, and chromosomal DNA. Our results show that the non-equilibrium reactions of ribosomes cause them to aggregate at the midcell, and this process facilitates the separation of the two daughter chromosomes. However, the observed effects are weaker in live cells than our one-dimensional reaction-diffusion model predicts. Rather than relying solely on active mRNA–ribosome dynamics, our data suggest that the closing division septum via steric interactions and potentially entropic forces between two DNA strands coupled to cell elongation act as additional mechanisms to ensure faithful partitioning of the nucleoids to two daughter cells.

**Significance:** The mitotic spindle separates chromosomes in eukaryotic cells, but bacteria lack this structure. It remains unclear how bacterial chromosomes partition before cell division. It has been hypothesized that non-equilibrium dynamics of polysomes, that is, mRNA-ribosome complexes, actively drive the separation of bacterial chromosomes. Using quantitative microscopy combined with computational modeling, we show that polysome dynamics facilitates the separation of daughter chromosomes in *Escherichia coli*, but this process does not constitute the sole mechanism. Our findings suggest that the closing division septum via steric interactions and potentially entropic forces between the two DNA strands act as additional mechanisms.

## Introduction

Chromosome replication and segregation are essential processes in the life cycle of a cell (1-3). Segregation can be divided into two stages: the individualization of daughter chromosomes and their further separation, which enables their partitioning into daughter cells during cytokinesis (4). Completion of both stages is required for the successful propagation of cells from one generation to the next.

In bacteria, the individualization stage, commonly referred to as DNA segregation, occurs for the most part concurrently with DNA replication (5,6). In *Escherichia coli*, the newly synthesized DNA loci near *oriC* begin to separate from each other with some delay after initiation of DNA replication (7). After this initial cohesion phase, the sister loci segregate from each other sequentially in the order in which they appear in the genome (8-10). Despite extensive previous research, the underlying molecular mechanism driving the segregation phase in bacteria remains poorly understood (2,3,11). Several active mechanisms for chromosome segregation have been investigated, including ParAB (ParA-ParB-*parS*) (2,11) and the homologous Spo0J-Soj system (12), but these are missing in many bacterial species, including *E. coli* (13,14). In addition to the Par-systems, structural maintenance of chromosomes (SMC) proteins, encoded by MukBEF in *E. coli*, have also been implicated in the active segregation (15,16). SMC involvement in segregation is postulated to occur via a successive condensation of DNA around *oriC* via loop extrusion and repulsion of condensed DNA masses away from each other due to excluded volume interactions and chain entropy (15,16). While this is a plausible scenario, MukBEF is not essential for DNA segregation in *E. coli*, except possibly at the fastest growth rates when multifork replication is present (17).

Rather than relying on dedicated protein machinery for segregation, which would expend energy through ATP hydrolysis, it is plausible that the free energy needed for the segregation of chromosomal DNA is derived mostly from the replication process itself. Replication creates two new proximal DNA strands, which, according to polymer physics, may experience a repulsive entropic force in confined cylindrical conditions (18,19). This scenario, which we refer to as the entropic segregation mechanism, has been supported by numerous computer simulations (20-27) but it has yet to be experimentally validated. However, the driving force from this so-called entropic mechanism vanishes once the spatial overlap between the two nucleoids becomes zero (27). Therefore, if only the entropic mechanism were present, the two individualized and segregated nucleoids would remain in contact rather than fully separating from each other. The entropic mechanism can, thus, explain the segregation but not the separation phase of the two nucleoids.

The question then arises: what drives chromosomes apart during the separation stage? In eukaryotes, the separation is carried out by the mitotic spindle, but no spindles are known to exist in bacteria. DNA translocase FtsK, which is a part of the bacterial cell division apparatus (divisome) (28-30), could be involved in this process (29). However, cells with inactivated FtsK (FtsK K997A) (31) appear to separate their nucleoids and partition them into two daughter cell compartments in the majority of cells (32).

Evidence for a different mechanism for separation was revealed by Wu *et al*. (33). These authors observed that in filamentous cells (more than 10 µm) with two nonreplicating nucleoids, the nucleoids separated from each other over large distances so that their centers stayed close to the cell’s ¼ positions (33). Wu *et al*. explained such positioning by osmotic pressure from the accumulation of macromolecular crowders, but their explanation relied on the assumption that these crowders are produced in a specific way outside the two nucleoids (33). An alternative explanation, which did not rely on this assumption, was proposed by Miangolarra *et al*. (34). These authors developed a model that included reactions between chromosomal DNA, mRNA, ribosomal subunits, and polyribosomes (polysomes), along with their diffusion within the cell. This reaction-diffusion model, which we refer to as the active mRNA-ribosome dynamics model, included reactions for mRNA synthesis, ribosome assembly and disassembly on mRNA, and mRNA degradation. The diffusion of these species was affected by the steric (excluded volume) interactions between these molecules. The model predicted that due to these interactions, the 30S and 50S ribosomal subunits can diffuse relatively unhindered into the nucleoid region, consistent with single-molecule tracking experiments (35,36), while 70S ribosomes, which are further assembled into polyribosomes (polysomes), cannot. The key insight from the model was a finding that mRNA synthesis within the nucleoid and the assembly of 30S and 50S ribosomal subunits into polysomes around these mRNAs create instability within the nucleoid. This instability splits the nucleoid into two distinct nucleoid lobes at some critical cell length (34). The model also predicted that the two daughter nucleoid centers would position at the quarter points of the cell once the transition occurred and stay in these locations even when cells filamented, consistent with the experimental observations by Wu *et al*. (33).

Here, we investigate the mechanisms that individualize chromosomes in the late stages of segregation and drive their separation in normal growth conditions. In particular, we test the active mRNA-ribosome dynamics model in non-filamenting cells growing in slow-growth conditions when chromosomal DNA has a simple topology. To this end, we use high-throughput microscopy in microfluidic devices to determine the distributions of ribosomes, nucleoids, and replication machinery during cell and replication cycles. We quantitatively compare the acquired experimental data to our own reaction-diffusion model, which is conceptually similar to the one proposed by Miangolarra *et al*. (34). We first find that the termination of DNA replication does not have a significant effect on the nucleoid segregation dynamics. Instead, DNA expands and daughter nucleoids separate in proportion to cell length increase during this stage of the cell cycle. Our data, when compared to the model, supports the idea that active ribosome dynamics facilitates the separation of two nucleoids in normal growth conditions, as recently reported (37). However, in cells where active ribosome dynamics are halted, we observe that the separation of daughter nucleoids still increases proportionally with cell length. In addition to this effect, we identify closing constriction, as a new mechanism that has a significant effect in separating the two daughter nucleoids and partitioning them into two daughter cells. Based on this evidence, we propose a multi-process model to explain the nucleoid segregation and separation dynamics in *E. coli*.

## Results

### The midcell DNA density minimum evolves independently of replication termination

DNA segregation cannot occur without DNA being first replicated. We first hypothesized that the replication solely drives the progression of segregation. In previous works on the segregation of nucleoids using tracking of fluorescently labeled loci in the chromosome (5,7,10,38,39), following bulk segregation (26,37), or labeling the replication terminus by MatP fusion (26,40), the segregation process had not been directly tracked during the replication cycle at the single-cell level. To do so now, we labeled the beta-clamp protein, DnaN, with a fluorescent fusion (Ypet-DnaN) (41). To follow the dynamics of the whole chromosome, we used HupA-mCherry, which has been tested before and is known to bind uniformly throughout the chromosome (42). We imaged cells in mother machine microfluidic devices (43,44), obtaining about 30-50 time points per cell cycle from several hundred cells (Movie S1-2, SI Fig. S1). We averaged these data over the cell population and produced kymographs (Fig. 1A-B). We chose to carry out measurements in slow-growth conditions where, at most, only a single replication cycle is ongoing (45) and where we could accurately determine timings for the initiation and termination of the replication (SI Fig. S2).

**Fig. 1.**
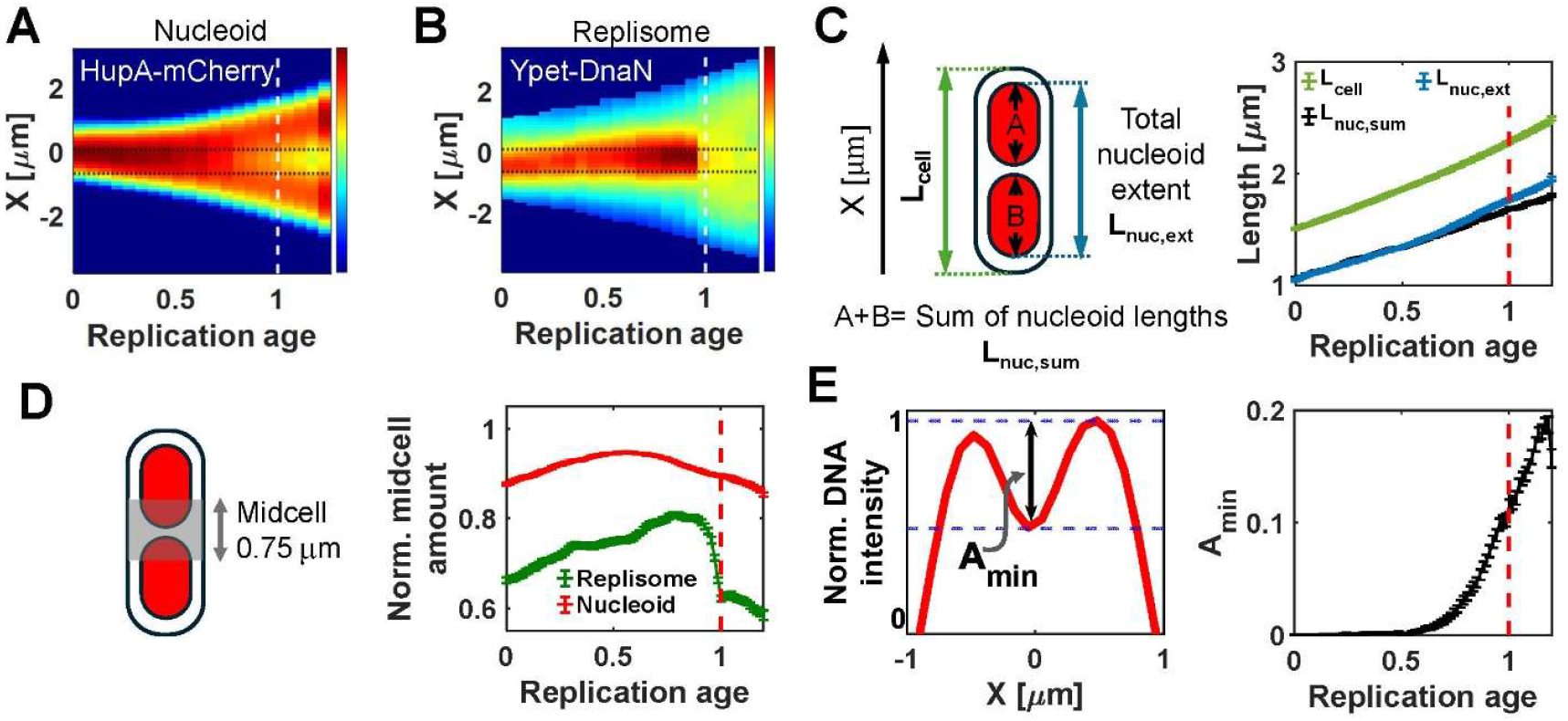
Segregation of nucleoids as a function of DNA replication time (age). (A) The population average DNA density distribution along the long axis of the cell as a function of replication cycle age. Time zero corresponds to replication initiation and one to termination (also indicated by a vertical dashed line). HupA-mCherry labeling is used to infer DNA density distribution. Red corresponds to high- and blue to low-intensity values. The dashed horizontal lines indicate a midcell region. (B) The same for the average replisome density distribution. Ypet-DnaN labeling is used to infer this distribution. (C) Left: definition of parameters characterizing the dimensions of nucleoids: *L*_*nuc,sum*_ and *L*_*nuc,ext*_. Right: change of cell length *L*_*cell*_, *L*_*nuc,sum*_ and *L*_*nuc,ext*_ as a function of replication cycle time. The dashed vertical line marks replication termination. (D) Left: schematics showing the region where HupA-mCherry and Ypet-DnaN midcell amount is measured (indicated also by the horizontal dashed lines in panels A and B). The midcell amount is based on the integrated intensity of fluorescent labels in this band. Right: the normalized midcell amount for nucleoid (red) and replisome (green) as a function of replication age. The curves for individual cells are normalized by the maximum value during the whole measurement period and then averaged over the cell population. (E) Left: schematics showing the DNA density distribution for a single cell at a single time point and how the depth of the local minimum, *A*_*min*_, is defined. Right: *A*_*min*_ as a replication cycle time. Error bars in panels C-E are s.e.m. N=438

To characterize the segregation process, we determined the total extent of all nucleoids in the cell, *L*_*nuc,ext*_, and the sum of nucleoid lengths, *L*_*nuc,sum*_ (Fig. 1C; for details, see Materials and Methods). These measurements showed that the outer edges of the nucleoids, as determined by *L*_*nuc,ext*_, stayed at about a constant distance (0.5/2 µm ≈ 0.25 µm) from the cell ends throughout the replication cycle (Fig. 1C). Notably, the slope of *L*_*nuc,ext*_ and *L*_*nuc,sum*_ vs. time curves did not show any discernible change at the termination of replication (Fig. 1C). We would have expected the slope of these curves to decrease at the termination because the increase in the amount of chromosomal DNA stopped. To further verify this result, we aligned the curves from individual cells relative to replication termination time, *Trt*, but still found only a minor changes in the slope of the curves at the replication termination (Fig. S3). We also found that the outer edges of the nucleoids remain at approximately constant distance from the cell poles (*i. e. L*_*cell*_ − *L*_*nuc,ext*_ ≈ *const*) not only throughout the replication cycle but also during the division cycle in both slow (SI Fig. S4) and moderately fast growth conditions (SI Fig. S5). Note that the measured exclu. sion distance of nucleoids from poles (0.25 µm) is slightly less than the length of the hemispherical cap region (0.3-0.35 µm) (46). The data thus show that instead of nucleoid elongation and separation halting at the replication termination, the length of nucleoids (*L*_*nuc,ext*_ and *L*_*nuc,ext*_) continues to increase and the increase is in proportion to cell length. This dependence has also been noticed in some earlier studies (26,33,47), although the replication period was not directly determined in these works.

In addition to *L*_*nuc,sum*_ and *L*_*nuc,ext*_, we also determined the DNA density at midcell during the replication cycle as a different measure of DNA segregation and separation (Fig. 1D). In the early stages of replication, there was a maximum DNA density at midcell (Fig. 1A). Upon completion of about 60% of the replication cycle, the DNA amount at the midcell started to decrease (Fig.1D). This decrease corresponded to the formation of a local DNA density minimum (Fig. 1E). As with our other nucleoid measurements, the midcell minimum continued to decline further after replication termination without any distinct changes in the slope of the curve. Altogether, these results, combined with the nucleoid length measurements, indicate that while chromosome replication is a prerequisite for segregation, nucleoid segregation is not strongly driven by the replication process itself at late stages of the replication cycle and after its completion. Instead, the separation of two nucleoids follows overall cell growth.

### Correlated and anti-correlated periods of DNA and ribosome density distributions during the cell cycle

In balanced growth, cell size increases proportionally to ribosome numbers. We hypothesized that the DNA segregation at late stages of the replication cycle could be driven by active ribosome dynamics that involve mRNA synthesis and degradation and are associated with the assembly and disassembly of polyribosomes (34). To test this hypothesis, we constructed a strain (for details, see SI Text, *Strain construction*) where the nucleoids were labeled with HupA-mYpet fluorescent marker (26,42) and ribosomes with RpsB-mCherry; the latter being part of the 30S ribosomal subunit (48,49). Both labels had been tested in previous studies (26,42,48,49). Since this strain lacked a marker for replication, we followed nucleoid segregation and ribosome accumulations throughout the division cycle (Fig. 2A-C, SI Fig. 6, Movie S3-4), which we customarily refer to as the cell cycle.

**Figure 2.**
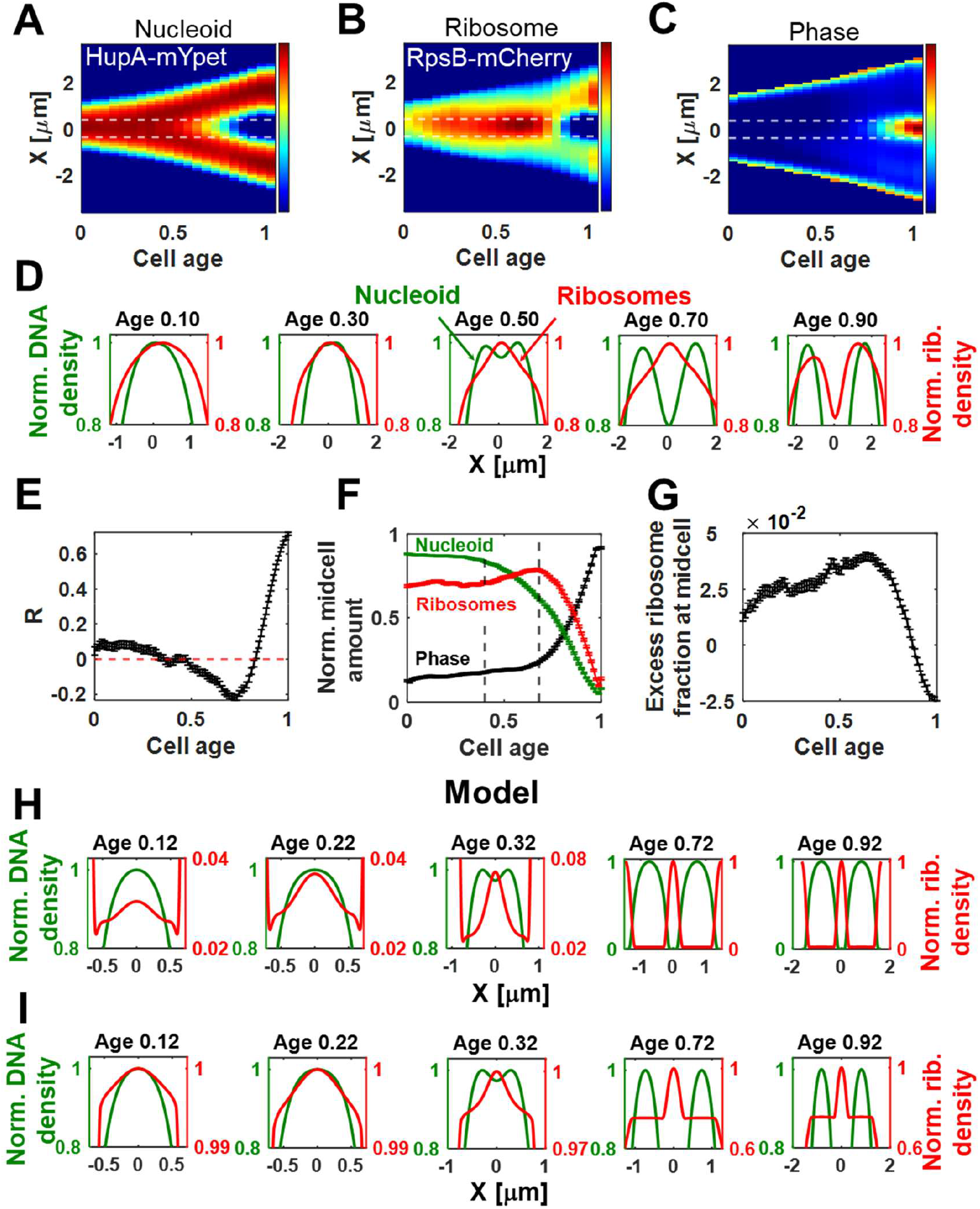
Cell-cycle dependent changes in DNA and ribosome density distributions and their comparison to model predictions. (A) Population averaged kymograph of DNA density distribution (HupA-mYpet) along the long axis of the cell as a function of cell age. Zero corresponds to cell birth and one to division. Red corresponds to high- and blue to low-intensity values. The dashed horizontal lines indicate a midcell region. (B) The same for ribosome density (RpsB-mCherry) distribution. (C) Phase contrast kymograph for the same cell population. (D) The normalized population-averaged density of nucleoid (green) and ribosomes (red) along the long axis of the cell at different cell ages. The curves are normalized by their peak values.Pearson correlation coefficient, *R*, between nucleoid and ribosome densities as a function of cell age. The coefficient is calculated for each cell at different time points and then averaged over the cell population. The integrated intensity within the 0.75 µm wide band around the cell middle is shown for the nucleoid (green), ribosome (red) and phase contrast signals (black). The curves from individual cells are normalized by their peak values and then averaged over the cell population. The left vertical dashed line corresponds to cell age of 0.40 and the right one to 0.68. (G) The relative excess fraction of ribosomes in the midcell band as a function of cell cycle time. The curve is derived from a fitting of the ribosome signal (see Methods). N=486 for panels A-G. The error bars are s.e.m. for panels E-G. (H) The normalized DNA and ribosome densities along the long axis of the cell at different cell ages, as predicted by the coupled DNA and ribosome dynamics model. The curves are normalized by their peak values. Note that the DNA (left) and ribosome (right) signals have different scales, and the cell ages are comparable but not identical as those in panel D.(I) The same curves that are recalculated, assuming a concentric shell of ribosomes around the nucleoid (for details, see the Concentric Shell Model in Methods). The shell’s inner radius is 0.5×*R*_*cell*_.

In the first half of the cell cycle, the population-averaged data show a unimodal distribution of DNA with a maximum close to the cell center (Fig. 2D, Age 0.10, 0.30). The cell cycle age (age for short) here is expressed as the fraction of a completed cell cycle. Notably, the ribosome density distribution is similar to the DNA density distribution from the same cells, showing a broad density maximum at the midcell. Here, the measured ribosome density is the sum of free 30S ribosome subunits and densities of 30S subunits incorporated into free ribosome particles and polysomes. The co-localization of the two distribution maxima appears inconsistent with equilibrium thermodynamic models (50,51), where ribosomes phase-separate from DNA-rich regions. Note that there are no local maxima in the ribosome density at cell poles in distributions in Fig. 2D. The excess ribosome density at cell poles is not expected to result in maxima in the extreme ends of 1D density distributions, which are projections of the 3D density onto the long axis of the cell. The maxima are missing because of the rounded shape of the poles and diffraction-limited imaging. Also, there could be an excess ribosome density present in the vicinity of the inner membrane of the cell, but this is not visible in microscopy images (SI Fig. S6) and in the density distributions along the short axes of the cell (SI Fig. S7), likely due to projection effects.

As the cell cycle progresses, the nucleoid expands, and the population-average DNA density distribution becomes bimodal (also referred to as bilobed nucleoid (7)) with a local minimum near its center (Fig.2D, Age=0.50). This minimum is accompanied by a sharper ribosome density maximum at the same location. In this case, the observed densities of ribosomes and chromosomal DNA at the midcell are anti-correlated at age = 0.5, consistent with the phase separation of nucleoid and ribosomes, as predicted by some equilibrium thermodynamic models (50,51). The excess ribosome density increases, and the DNA density at midcell decreases further as the cell cycle progresses (Fig. 2D, Age=0.70). However, at the last stages of the cell cycle, the ribosome density decreases instead of further increasing, while the DNA density decreases too (Fig. 2D, Age=0.90). Again, this positively correlated behavior cannot be explained by equilibrium considerations of volume exclusion between ribosomes and chromosomes.

To show these changes occurring in the cell cycle, we plot the Pearson correlation coefficient, *R*, between the DNA and ribosome distributions along the long axis of the cell as a function of cell age (Fig. 2E). At the beginning of the cell cycle, the two distributions show small but positive *R* values, as expected based on the coincidence of their maxima. At about cell age 0.5, the correlation coefficient starts to decrease and becomes negative. This decrease in *R* corresponds to a concurrent decrease in DNA and an increase in ribosome amounts at midcell (Fig. 2F). The correlation coefficient reaches its minimum at cell age ∼0.75, then starts to increase and becomes positive by the end of the cell cycle. At cell cycle age ∼0.75, the midcell ribosome numbers reach their peak and start to decrease thereafter, while the amount of DNA at midcell continues to decrease until the end of the cell cycle (Fig. 2F). To understand the cause of the decrease in ribosome numbers at midcell at the end of the cell cycle, we compared these data with the midcell phase contrast signal from the same cells (Fig. 2F, black line). There is an upturn in the phase contrast signal at the time when the ribosome numbers start to decrease. The increase in the phase contrast signal signifies the onset of constriction (51). We find that at single-cell level the midcell ribosome density starts to decrease after the onset of constriction (SI Fig. S8A) and shortly (10 min) before cells divide (SI Fig. S8B). The simplest explanation for the decrease in midcell ribosome numbers after the onset of constriction is the decrease in cell volume in the constricted region.

Based on traces in Fig. 2D, the excess fraction of ribosomes in the midcell is small throughout the cell cycle. To quantify this fraction, we decompose the ribosomal density distribution into a midcell fraction and a uniform background by fitting the data from individual cells (see SI Text, *Quantification of midcell amount and the excess midcell fraction*). We find that the population-average excess fraction of ribosomes is only 4% of the cell total at its peak at cell age ∼0.75 (Fig. 2G). At the same time, the excess fraction before the DNA density minimum at midcell begins to form is 2.5%. The equilibrium thermodynamic models (50,51) can qualitatively explain the increase in the excess fraction from 2.5% to 4%. However, they fail to explain the 2.5% fraction present in the early cell cycle when there is no DNA density minimum at the center of the nucleoid.

To verify the validity of the above findings, we constructed a strain in which the fluorescent mCherry fusion was added to the C-terminus of the 50S ribosomal protein RplI (RplI-mCherry) instead of labeling the 30S subunit (SI Fig. S9A-C). This label has also been previously characterized and tested (52). The cells with RplI-mCherry label showed the same behavior as the one with RpsB-mCherry (SI Fig. S9D-G), including co-localization of ribosome and DNA density distributions at the midcell at early cell cycle (SI Fig. S9D, Age=0.15) and comparably small excess fraction of ribosomes at midcell at its peak value (3.5%) (SI Fig. S9G).

We also investigated whether the same behaviors would occur in faster growth rates in M9 glucose + CAS medium (SI Fig. S10). Under these conditions, the replication cycle initiates predominantly in the previous cell cycle and completes at the early cell cycle (45). Accordingly, a minimum in the nucleoid density distribution is already observed at the midcell at cell birth (SI Fig. S10C). Consequently, unlike in slow-growth conditions, there is no positive correlation between DNA and ribosome density distribution during the early stages of the cell cycle (SI Fig. S10D). Aside from this difference, the other behaviors (SI Fig. S10E-F), including a small excess ribosome fraction at the cell center at its peak value (5.5%), are consistent with those observed under slower growth conditions.

### The DNA and ribosome correlation patterns in the early and midcell cycle are qualitatively explained by the coupled DNA and ribosome dynamics model

To understand if the observed ribosome and nucleoid localization patterns can be explained by the active dynamics of mRNA and ribosomes, we developed our own reaction-diffusion model (see SI Text, *The Coupled DNA and Ribosome Dynamics Model*). We then compared the calculated cell cycle-dependent DNA and ribosome densities from the model to the experimental data. Our 1D model is similar to the one developed by Miangolarra *et al*. (34), considering mRNA synthesis and degradation, as well as the assembly of polysomes (polyribosomes) from ribosomal subunits. Polysome and ribosomal subunits interact with each other and with DNA via excluded volume interactions. However, unlike the previously published model, ours uses a power-law term instead of a virial expansion to describe DNA (51,53) and polysome (51) self-interactions, as the latter has been tested in experiments (53). As a simplification, our model does not explicitly simulate bare mRNA and polysomes of varying sizes; instead, it considers only an average-sized polysome. Unlike the previous model (34), ours includes cell growth, assuming DNA and ribosome amounts increase with cell length (or volume).

To qualitatively compare the model to experimental data, we plotted the nucleoid and ribosome density distribution along the cell length at different cell cycle times (Fig. 2H, SI Fig. S11A-B). The modeled ribosome density represents the total density of ribosomal subunits (either 30S or 50S) and includes contributions from both polysome-associated and free ribosomal subunits (SI Fig. S11C–D). As such, this density is directly comparable to our experimental measurements. While the cell cycle times in Fig. 2H were chosen to approximate those in Fig. 2D, they are not identical because the model does not exactly reproduce the cell cycle timing when a single nucleoid starts to split into two lobes. Notably, however, the ribosome distribution from the model shows a small local maximum at the cell middle, coinciding with the maximum of the nucleoid distribution at the early stages of the cell cycle (Fig. 2H, Age 0.12, 0.22). This maximum is qualitatively consistent with experimental data (Fig. 2D, Age 0.10, 0.30). The midcell maximum in ribosome distribution arises in the model because 30S and 50S ribosomal subunits can diffuse into the nucleoid region, where they assemble into polysomes. Since the rate of polysome formation is proportional to DNA density in the model, the maxima of ribosome and DNA densities coincide.

The model predicts that excess ribosome density at the midcell cleaves the initial single nucleoid into two nucleoid masses at some critical cell length due to excluded volume interactions (Fig. 2H, Age=0.32). After this transition, ribosome and DNA density distributions become anti-correlated (Fig. 2H, Age=0.72; SI Fig. S11E), which is consistent with experimental observations. Separation of the two DNA masses in the model occurs rapidly, within 5% (0.05) of the cell cycle. During this transition, the excess fraction of ribosomes at midcell reaches 48% and approximately maintains this value for the rest of the cycle (SI Fig. S11F). In contrast, the experiment shows that the fraction reaches only 4.5% (Fig. 2G). The model thus correctly predicts a local maximum in ribosome density at the cell middle in the early stages. However, it shows a significantly higher accumulation of ribosomes in the cell middle in the later stages of the cell cycle compared to the experimental data. We hypothesized that the substantially higher ribosome fractions observed in the model are due to its 1D nature. In a 3D cell, ribosomes could form an outer shell around the nucleoid, as has been proposed by earlier single-molecule tracking experiments of ribosomes (34). Since this shell could not be derived from our 1D model, we added its effect *ad hoc* to our model (SI Fig. S12A-C). Additionally, we assumed a uniform ribosome volume density in the shell that matches the density predicted by the 1D model at the cell poles (for details, see SI Text, *The Concentric Shell Model*). With this addition, the experimental (Fig. 2D) and modeling densities (Fig. 2I) showed similar values, with the model yielding about 3% excess of ribosomes at midcell at its peak (SI Fig. S12E). However, to reach this agreement, the inner radius of the added shell needed to be 50% of the cell radius, while it has been estimated based on single-molecule tracking experiments that the inner radius is about 80% of the cell’s radius (35). These comparisons suggest that the coupled DNA and ribosome dynamics model could better explain the data if it were implemented in 3D.

### Constriction pushes aside daughter nucleoids

In the late stages of the cell cycle, the model and experimental data do not agree with each other. The data reveal decreasing ribosome and nucleoid densities at the midcell (Fig. 2F), whereas the model maintains almost a constant fraction (SI Fig. S11 A-B, F). The discrepancy appears to be due to the closing constriction. To further understand the role of constriction closure in separating daughter nucleoids, we carried out experiments in which the beta-lactam antibiotic cephalexin inhibited constriction formation. It is well known that despite the inhibition of constriction, cephalexin-treated cells continue to grow in volume and replicate DNA at the same rates as untreated cells (54). We then compared the changes in ribosome and DNA densities between the cephalexin-treated and untreated cells. Since cephalexin prevents cell division, cell age cannot be used to track the progression of DNA and ribosome densities, as done in the previous measurements in Fig. 2. Instead, we determined the timing for the first appearance of the local DNA density minimum at the cell center, *T*_*min*_, following the procedure defined earlier (45) (for details see SI Text, *Determining A*_*min*_ *and T*_*min*_ *from DNA and ribosomal distributions*). The appearance of this minimum correlated with the replication termination timing, *T*_*rt*_, and was detected on average shortly after *T*_*rt*_ in the strain that carries DNA and replisome markers (SI Fig. S13A-B). We then aligned the time-dependent data relative *T*_*min*_ (Fig. 3A, B) processing the data from untreated cells the same way (SI Fig. S14A-B). At *T*_*min*_, both the treated and untreated cells show an increase in midcell ribosome numbers (Fig. 3C) and a simultaneous decrease in the midcell DNA amount (Fig. 3D), consistent with the excluded volume interactions between ribosomes and DNA. However, at later times (*time* − *T*_*min*_ > 50 min), the midcell ribosome numbers stay constant in treated cells, while they decrease in untreated ones (Fig. 3C). This comparison further confirms that the decrease in ribosome numbers in untreated cells is caused by the closing constriction.

**Figure 3.**
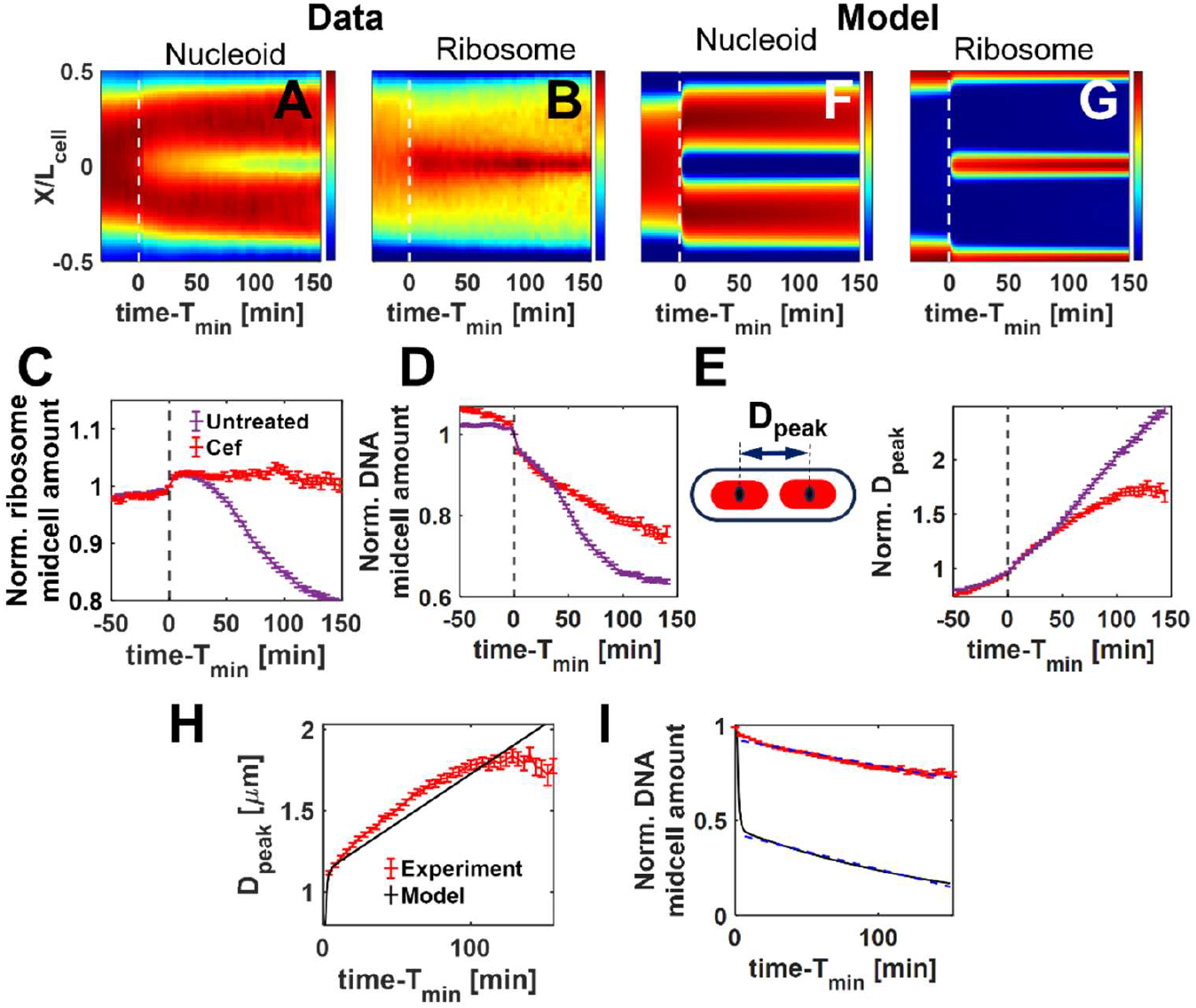
Segregation of nucleoids with and without closing constriction. (A) A population-averaged kymograph of DNA density along the long axis of the cell as a function of absolute (unscaled) time from the formation of local DNA density minimum, *T*_*min*_, for cephalexin-treated non-constricting cells. The vertical axis shows normalized cell length, which remains unchanged throughout the experiment. Note that cells filament in this measurement. (B) The same for ribosome density distribution in this cell population. (C) The average number of ribosomes at midcell as a function of time from *T*_*min*_ for cephalexin-treated (red) and untreated cells (violet). The curves are normalized so that the amounts are one at time *T*_*min*_. (D) The normalized DNA amount at midcell is presented the same way as in the previous panel. (E) The average distance between nucleoid centers (peak distance) as a function of time from *T*_*min*_ for cephalexin-treated (red) and untreated cells (violet). The distances are normalized at *T*_*min*_. The schematic on the top shows how the peak distance is defined. (F) Kymograph of nucleoid density from the coupled DNA and ribosome dynamics model presented the same way as the experimental data in panel A. (G) Kymograph of ribosome density from the model. (H) The peak distance as a function of time relative to *T*_*min*_ from cephalexin-treated cells (red) and from the model (black). (I) Normalized midcell DNA amount for cephalexin-treated cells (red) and model (black). The linear fits to both curves are shown by dashed lines. The model is fitted for time − *T*_*min*_ > 5 min. (J) The fit coefficients for panel I are listed in Table S3. All error bars are s.e.m. N=235.

The data also indicates that, in addition to reducing the number of ribosomes at midcell, the constriction affects the separation of the two daughter nucleoids. Comparison of DNA density at the midcell reveals a clear decrease in DNA in untreated cells relative to treated ones for *time* − *T*_*min*_ > 50 min (Fig. 3D). The effect of constriction is also noticeable in the measured distance between two nucleoid lobes, increasing more significantly in untreated cells than in treated ones (Fig. 3E). Altogether, these data show that the closing constriction separates two chromosomes in the late stages of the cell cycle. The effect is not caused by FtsK, since FtsK is recruited to the divisome in both cephalexin-treated and untreated cells (55). The constriction-driven separation appears, thus, to be a steric effect: the closing constriction physically pushes aside the daughter nucleoids.

In untreated cells in slow-growth conditions, the onset of constriction occurs effectively at the time of the DNA density minimum, *T*_*min*_ (45). Therefore, the effects arising from the active ribosome dynamics and constriction on the separation of two daughter nucleoids are superimposed for *time* > *T*_*min*_. However, the data from cephalexin-treated cells can be directly compared with our model (for *time* > *T*_*min*_) because it excludes effects arising from the constriction process. The comparison shows that the DNA and ribosome density distributions predicted by the model (Fig. 3 F-G) are at a qualitative level similar to the experimental data from cephalexin-treated cells (Fig. 3A-B). In quantitative comparisons, the model reproduced some trends in the experimental data but missed others. The model quantitatively correctly predicts the distance between the peaks of two nucleoid lobes (Fig. 3H) but fails to predict the time-dependent decrease in the amount of DNA at midcell (Fig. 3I). In the experiments, the DNA amount changes only 10% over 60 min, while in the model it changes 50% within about 5 min. However, after this initial 5 min period, the model and experiment show comparable rates of DNA density decrease at the midcell (Fig. 3I). The reason for the ten-fold slower segregation dynamics observed in the experiments could be related to the choice of model parameters. Reducing the diffusion coefficient of DNA 10-fold considerably slowed down the rate of the change in DNA amount at the midcell, although the cell cycle time for the transition for DNA density from high to low values shifted to later times (SI Fig. S15). Further adjustments of model parameters may overcome this shift. Altogether, the model shows qualitative agreement with the experimental data. Its quantitative accuracy could be potentially improved by extending the model to three dimensions and refining its unknown parameters through additional experiments and optimization.

### The effect of disruption of polysome formation on nucleoid separation

To perturbatively test the effects of active ribosome dynamics on the separation of two nucleoids, we treated *E. coli* cells with a lethal concentration of rifampicin (300 µg/ml), which inhibits the initiation of RNA synthesis (56) but does not stop ongoing DNA replication (6). As in the previous measurements, we also added cephalexin to the culture medium at the same time as rifampicin to prevent cells from constricting. Since rifampicin inhibits RNA synthesis, no new polysomes can form after the drug reaches the cells. The existing transcripts can still be completed, but this occurs on the timescale of tens of seconds for a single gene and about a minute for longer operons. These timescales are much shorter than our measurement frame rate (4 min). On a slightly longer timescale (3-5 min (57)), the existing polysomes disassemble as their associated mRNAs degrade, releasing ribosomal subunits.

To understand the effects arising from these processes, we compared data from cells treated simultaneously with rifampicin and cephalexin (SI Fig. S16A–C) to cells treated only with cephalexin (SI Fig. S16D–F). In this comparison, we align the time-dependent data to the antibiotic treatment start time, *T*_*ab*_, instead of *T*_*min*_ because most Rif-treated cells lack a detectable midcell DNA density minimum. Additionally, we excluded from the analysis cells with visible constriction at the start of the treatment. As expected, the localized accumulation of ribosomes at the midcell is not present in cells following rifampicin treatment (SI Fig. S16B, C) due to polysome dissociation and subsequent diffusion of free ribosomal subunits onto the nucleoid region.

Following rifampicin addition, the midcell DNA density minimum almost immediately ceases to deepen, although it persists rather than disappearing (Fig. 4A). If steric interactions between DNA and polysomes alone created this minimum, we would expect its complete disappearance on a timescale comparable to polysome dissociation. In contrast, cephalexin-only treated cells continue to show a gradual decrease in midcell DNA density (Fig. 4A). This difference between treatment conditions indicates that active mRNA-ribosome dynamics contribute significantly to establishing the midcell DNA density minimum but are not solely responsible.

**Figure 4.**
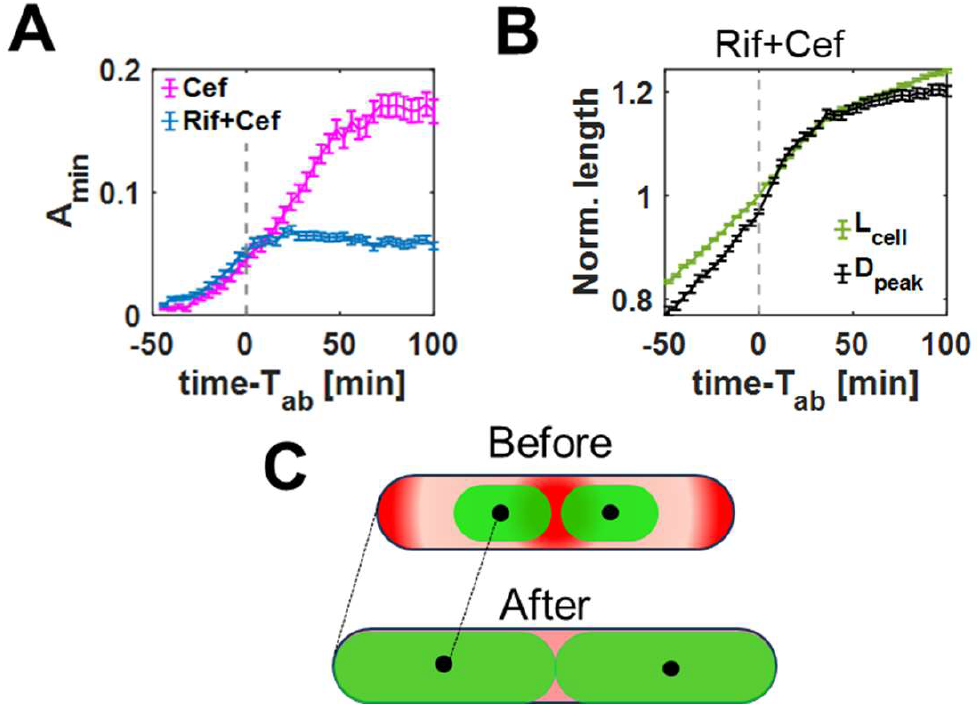
The effects from the disruption of active mRNA-polysome dynamics with rifampicin (Rif) on nucleoid separation. (A) The depth of the local DNA density minimum at midcell as a function of time relative to the start of antibiotic treatment with Rif + Cef (blue) and Cef only (pink). The times are relative to the start of antibiotic treatment, *T*_*ab*_, with Rif and Cef (cephalexin), and with only Cef, respectively. All analyzed cells in this figure are non-constricting. (B) The average cell length (green) and the nucleoid peak distance (black) as a function of time from the start of the treatment for Rif + Cef treated cells. The curves from individual cells are normalized to one at *T*_*ab*_. Error bars in panels G and H are s.e.m. (C) Schematic summarizing the expansion and movement of nucleoids and ribosomes during Rif treatment. The top corresponds to a cell just before and bottom after Rif (+Cef) treatment. *N* = 786 for Rif+Cef and *N* = 310 for only Cef treated cells.

We next examined nucleoid segregation by measuring the distance between nucleoid centers (*D*_*peak*_) in rifampicin-treated cells (Fig. 4B, SI Fig. S17A). This distance continues to increase for approximately 50 minutes, long after active ribosome dynamics ceases (within minutes of rifampicin addition) and polysomes dissociate (3–5 min). Under normal growth conditions, nucleoids separate proportionally to cell elongation (Fig. 1C). To determine if a similar relationship exists following rifampicin treatment, we compared nucleoid center distances to cell length increases (SI Fig. S17), normalizing both metrics at the time of antibiotic addition (Fig. 4B). Indeed, even without active ribosome dynamics, nucleoid centers continue separating proportionally to cell elongation, which continues for about 50 min after treatment starts. This finding indicates that cell growth alone is sufficient to drive nucleoid segregation (Fig. 4C). However, rifampicin treatment also results in nucleoid expansion, causing increased midcell DNA density when two nucleoids are present (Fig. 4A). These findings show that chromosome segregation progresses despite active ribosome dynamics and polysome dissociation, while the separation of daughter chromosomes at the midcell is affected.

To compare these data to our model, we divide the rifampicin-treated cells into three age groups based on their nucleoid segregation stages. This grouping was necessary because rifampicin treatment starts at different cell cycle ages for the asynchronous cell population in Fig. 4. The first group, representing 12% of cells, includes those cells where *T*_*min*_ was detected before treatment began (*T*_*min*_ < *T*_*ab*_, *N* = 92) (SI Fig. S18). The second group, comprising 17% of cells, includes cells in which nucleoid separation was detected after treatment started (*T*_*min*_ > *T*_*ab*_, *N* = 138). The largest group, accounting for 71% of cells (*N* = 556), shows no detectable nucleoid separation either before or after treatment. Our following analysis focuses on the first two groups, as the third group shows minimal changes in ribosome and nucleoid distributions after the treatment. To model these two groups of cells, we set the polysome formation rate to zero at the start of the treatment. We then evolve the model so that cell length changes the same way as in the experiments (see SI Text, *Rifampicin Treatment in the Model*, for details).

The group of cells that were late in their replication cycle at the onset of antibiotic treatment (*T*_*min*_ < *T*_*ab*_), show a local DNA density minimum (Fig. 5A) and a midcell ribosome accumulation at the start of rifampicin treatment (Fig. 5B). The midcell ribosome accumulation vanishes within about 16 min from the start of the treatment (Fig. 5B-C). The same occurs in the model, although at a faster rate (Fig. 5E-F). At the same time, the model and the data qualitatively differ in how DNA distributions evolved. The experimental data show that the midcell DNA density minimum persists, albeit becoming shallower over a 50-minute period, before plateauing (Fig. 5A, C, SI Fig. S19A). In contrast, in the model, the two nucleoid masses merge so that the local DNA density minimum at the midcell completely vanishes (Fig. 5D-F, SI Fig. S19B-C). The coupled DNA ribosome dynamics model is thus unable to explain the persistence of the experimentally observed nucleoid minimum after treatment.

**Figure 5.**
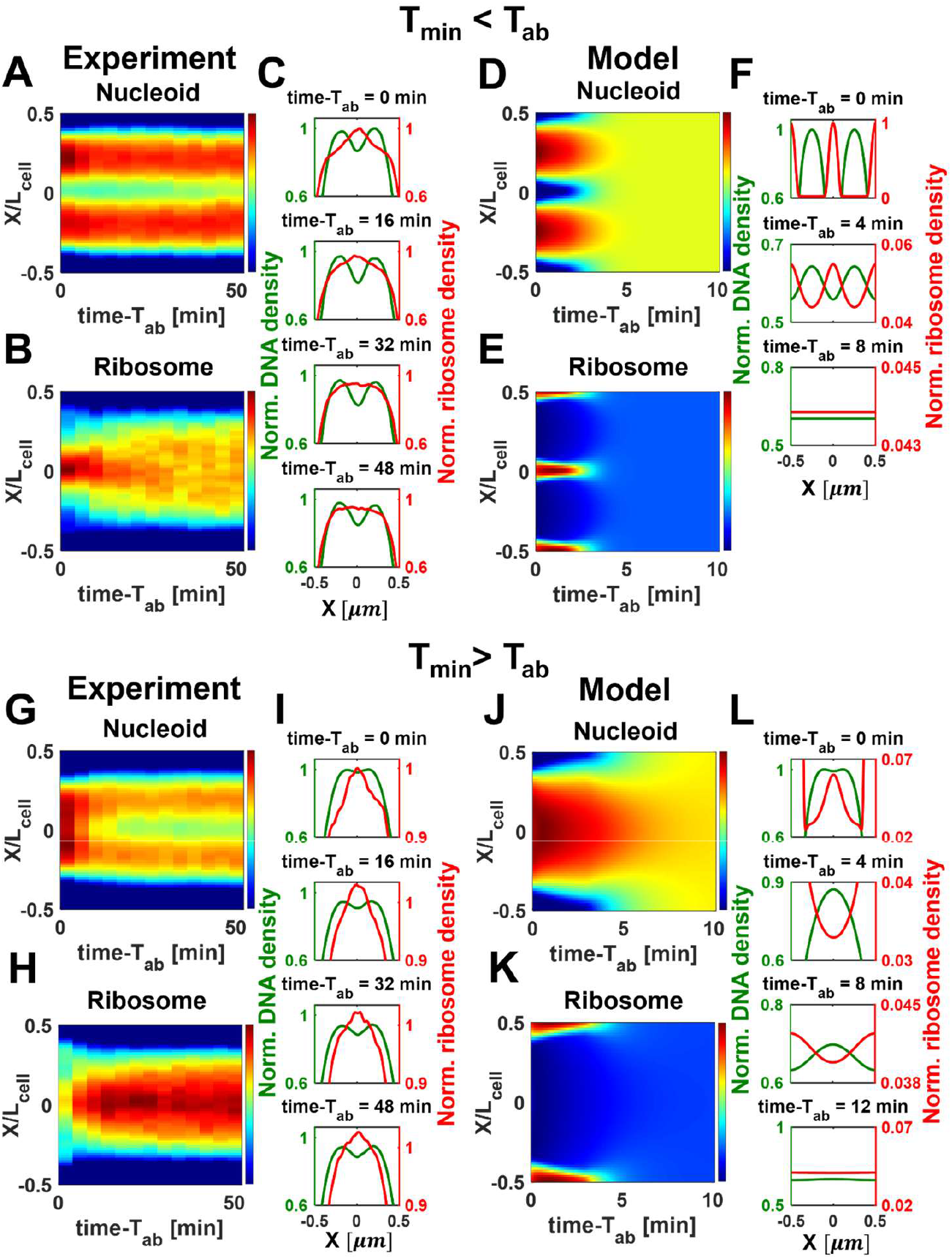
Comparing DNA and ribosome dynamics in Rif+Cef treated cells to the coupled DNA and ribosome dynamics model. Cells from two age groups are compared. (A-F) Data and model for cells that have formed a local DNA density minimum before the antibiotic treatment, *T*_*min*_ < *T*_*ab*_ (*N* = 92). (A, B) The kymographs of DNA and ribosome density, respectively. The times are relative to the start of antibiotic treatment. (C) The average DNA (green) and ribosome (red) distributions along the long axis of the cell at different time points of experiment. (D, E) The DNA and ribosome density kymographs, respectively, from the model corresponding to group of cells with *T*_*min*_ < *T*_*ab*_. The cell length in the beginning of the treatment is 2.87 µm. Note shorter times in the model compared to experimental data. (F) The average DNA (green) and ribosome (red) distributions along the long axis of the cell at different time points from the model. (G-L) Data and model for cells that formed a local DNA density minimum after the start of the antibiotic treatment, *T*_*min*_ > *T*_*ab*_ (*N* = 138). (G,H) The kymographs of DNA and ribosome density, respectively. (I) The average DNA (green) and ribosome (red) distributions along the long axis of the cell at different time points of the experiment. (J, K) The DNA and ribosome density kymographs, respectively, from the model corresponding to group of cells with *T*_*min*_ > *T*_*ab*_. The cell length in the beginning of the treatment is 2.24 µm. Note shorter times in the model compared to experimental data. (F) The average DNA (green) and ribosome (red) distributions along the long axis of the cell at different time points from the model.

We next examine the second group of cells (*T*_*min*_ > *T*_*ab*_) where the DNA density minimum develops after the start of rifampicin treatment (Fig. 5G-I, SI Fig. S20A). Although the observed DNA density minimum is small in this group, its presence also contradicts the active mRNA-ribosome dynamics model. In the model, the midcell DNA density minimum does not form without active ribosome dynamics (Fig. 5J-L, SI Fig. S20B-C). The lack of a minimum is a qualitative feature of the model, independent of parameter selection. This results from the overdamped nature of diffusive processes. Once the driving force is removed, that is, the formation of polysomes is halted, then the separation of the two nucleoid lobes stops immediately (SI Fig. S20B-C). The qualitative discrepancy between the model and the data for cells with *T*_*min*_ > 0 shows that the active dynamics of mRNA and ribosomes is not responsible for DNA separation in this group of cells.

## Discussion

### Active mRNA-ribosome dynamics

This study used experiments and modeling to examine the role of mRNA-ribosome dynamics in the segregation and separation of chromosomal DNA in *E. coli* cells. A characteristic feature of this mechanism, as also observed in our measurements (Fig. 2), is the accumulation of polyribosomes (polysomes) at the nucleoid (and cell) center before it starts to split into two lobes. According to our model, the accumulation of polyribosomes results from strong excluded volume interactions between polysomes and chromosomal DNA (51), leading to the exclusion of DNA from the midcell region. These qualitative features of the model are consistent with our data in steady-state growth conditions. Perturbative measurements with rifampicin treatment furthermore show that the nucleoid split does not develop in the majority of cells (77%) when polysome formation is inhibited due to the lack of mRNA substrates. However, these measurements also show that in cells where nucleoid lobes have already formed at the start of the treatment, the nucleoid centers continue to move apart as the cells continue to elongate (Fig. 4H). The latter finding indicates that the active mRNA-ribosome dynamics is not the sole process driving the late segregation of nucleoids. Similar conclusions were recently reported by Papagiannakis *et al*., although based on different evidence (37).

### Constriction effect

Our measurements also indicate that, in addition to active mRNA-ribosome dynamics, a closing constriction plays an important role in separating the two daughter nucleoids and partitioning them into daughter cell compartments (Fig. 3). This likely occurs via steric interactions between the constriction and DNA. These steric interactions could potentially be mediated by crowders, ensuring that the inner membrane and chromosomal DNA do not come into direct contact. Interestingly, earlier research has shown that the onset of constriction is blocked until DNA segregation has progressed to the point where there is a local DNA density minimum at midcell/mid-nucleoid (45). If the constriction were to start closing earlier, then it could likely result in the trapping of incompletely replicated DNA under the constriction. Altogether, DNA segregation to the point where there is a local DNA density minimum at the midcell appears to trigger the onset of constriction, which completely separates one daughter chromosome from another. The segregation process can thus be self-amplifying through the formation of the constriction.

### Evidence for additional mechanisms

In addition to the active mRNA-ribosome dynamics and closing constriction it is known from previous research that DNA translocase FtsK (58,59) provides a DNA partitioning mechanism (29,32). FtsK usually acts on about a 400 kb region near the replication termination region around the *dif*-site (29), but it can directionally pump more than 3 Mb of chromosomal DNA across a closing constriction (32). This activity mostly occurs at the end of the constriction phase when the pore separating the two daughters has a diameter of less than 250 nm (32). The pore does not close fully until all the DNA is pumped across it. Altogether, FtsK effectively guarantees a fail-safe completion of DNA partitioning. However, in most cell divisions, FtsK is dispensable, as mutant cells are viable even when FtsK translocase activity has been inhibited (32,60).

Our data suggest that, beyond active mRNA-ribosome dynamics, closing constriction, and FtsK-driven translocation, additional mechanism(s) drive nucleoid segregation and separation. The main evidence for this hypothetical mechanism comes from our finding that the two nucleoid centers move apart in proportion to cell elongation. This movement also occurs during cephalexin and combined rifampicin and cephalexin treatment. What could be this mechanism(s)? First, it can be ruled out that this process is due to the equilibrium macromolecular crowding via excluded volume effects, which remains present after rifampicin treatment. Indeed, passive crowding without reactions could lead to aggregation of DNA at high crowding levels or simply hinder DNA segregation and separation due to increased viscosity. Consistent with this view, a fusion of nucleoids can be observed in *E. coli* cells treated with chloramphenicol (61) – an antibiotic which stops protein synthesis but not RNA synthesis. In this case, the high level of macromolecular crowding appears to result from an increased level of RNA species (rRNA, tRNA, mRNA) in the cytosol (62). Alternatively, it has been proposed that cell elongation could drive nucleoid segregation via transertion linkages (63). However, this hypothesis can also be ruled out by our rifampicin measurements because the linkages would disappear within a few minutes from the start of the treatment, whereas the daughter nucleoids continue to move apart for a much longer period (50 min).

At the same time, the cell elongation-dependent movement of nucleoid lobes apart is potentially consistent with the entropic segregation mechanism (17,18,25) under certain conditions. In particular, the entropic segregation mechanism could explain that the distance between nucleoid lobes (*D*_*peak*_) continues to increase in proportion to cell length during rifampicin treatment (Fig. 4B, SI Fig. S17). During rifampicin treatment, macromolecular crowding effects weaken because of the dissociation of polysomes to ribosomal subunits (64). As a result, the chromosomal DNA expands and fills almost the whole cell volume (50,51). According to the entropic segregation mechanism, one chromosome will occupy one half of the cell, and the other chromosome the other half. The centers of each nucleoid are then at the ¼ position of the cell, irrespective of cell length (33), and the distance between the lobes is proportional to cell length as observed in our experiments. In our experiments, the nucleoids do not strictly fill the whole cell volume but remain excluded from the poles (46). However, this exclusion does not significantly alter the above arguments because the exclusion from poles is small, and the daughter DNA strands overlap at cell center (Fig. 4A). As emphasized before, the overlap condition is strictly needed for the entropic mechanism to play any role in driving the daughter nucleoids apart, and it cannot explain nucleoid separation when there is no overlap.

In summary, our measurements and modeling, combined with findings from previous research, bring out the involvement of several partially overlapping processes in the segregation and separation of daughter chromosomes in *E. coli*. Based on these findings, we propose the following sequence of events unfolding during the cell cycle (SI Fig. S21): The entropic segregation mechanism may be involved during the segregation stage when two chromosomes overlap (18,19). When about half of the DNA is replicated in slow growth conditions, active mRNA-ribosome dynamics via excluded volume interactions become a sufficiently strong driving force to lead to a midcell DNA density minimum. At this point, the entropic mechanism of segregation still remains important, and, in fact, the associated force increases as more DNA is replicated (26,27). However, the entropic force vanishes once two DNA molecules no longer overlap. In parallel, the formation of the midcell DNA density minimum via the active mRNA-ribosome dynamics triggers the onset of constriction in slow-growth conditions (45). The constriction further drives the two daughter chromosomes apart and contributes to their separation. The final stages of DNA partitioning are further aided by FtsK if DNA becomes trapped by the closing constriction.

Further work is needed to validate all the proposed processes mentioned above. In particular, the existing evidence for the entropic mechanism remains indirect. Significant effort is also required to model more faithfully the active mRNA-ribosome dynamics implementing the model in a 3D setting and including the chain connectivity for DNA.

## Materials and Methods

Details of bacterial growth conditions, microchip design, fluorescence microscopy setup, image analysis, and modeling are given in the SI Text. The list of all strains and plasmids used in this study, including their sources, is presented in the SI, Table S1. All experimental and modeling data are used in the main text, and the SI figures are available from Zenodo (65). The Jupyter Notebooks used in our modeling are available in reference (66).

## Acknowledgments

The authors thank Rachel McCord and Steve Abel for their valuable comments, and Sriram Tiruvadi-Krishnan for help in strain construction. A part of this research was conducted at the Center for Nanophase Materials Sciences, which is a US Department of Energy Office of Science User Facility at Oak Ridge National Laboratory. This work was supported by the National Institutes of Health award GM127413 (JM) and NSF MCB2313719 (JM).

## Notes

### Competing Interest Statement

The authors have declared no competing interest.

### Summary of Updates

1.We added timings associated with the appearance of DNA and ribosomal density minima from single-cell data and comparison of these timings to other cell cycle events (the onset of constriction, cell cycle time, and termination of DNA replication) to SI (Fig. S8 and S13). 2.We added representative cell images (SI Figs. S1, S6). 3.We added population averaged kymographs showing ribosomal and DNA density variations along the short axes of the cell as a function of cell cycle time (SI Fig. S7). 4.We now show separately the contributions of ribosomal subunits and polysomes in the total ribosomal signal (SI Fig. S11 C-D). 5.We describe now in detail the main assumptions and simplifications of the model in the "Model Overview" section of the SI Text. 6.We added a comparative discussion of our model with those by Miangolarra et al. (PNAS 2021) and Papagiannakis et al. (eLife 2025) in a new section of the SI Text titled "Comparing our model to other recent reaction-diffusion models". 7.We performed a stability analysis of our model, included as two new figures (SI Figs. S21-S22) and a section titled "The sensitivity of the model" in the SI Text. 8.We simplified the discussion of effects arising from the closure of the constriction in the main text. 9.We rearranged the Discussion section and rewrote several paragraphs there for clarity. 10.To shorten the main text, we moved Fig. 6 and panels A-E of Fig. 4 to the SI. We left out panels J & K of Fig. 3 and related discussion as it appeared tangential to the main message of our work.

