## Supporting Information for "The role of active mRNA-ribosome dynamics and closing constriction in daughter chromosome separation in *Escherichia coli*"

**This PDF file includes:**

- Supporting Information Text
- Figures S1 to S22
- Tables S1 to S6
- Captions for movies S1 to S4
- SI references

**Other supporting materials for this manuscript include the following:**

- Movies S1 to S4

### SI Text

#### Strain construction

All *E. coli* strains used in the reported experiments are derivatives of K12 BW27783 obtained from the Yale Coli Genetic Stock Center (CGSC#: 12119). Information on all strains is listed in Table S1. Oligonucleotide information is given in Table S2.

To follow DNA replication and chromosome dynamics simultaneously, we constructed strain STK32 (*hupA::hupA-mCherry-frt*, *dnaN::frt-Ypet-dnaN*) using P1 transduction. The *hupA-mCherry-frt-aph-frt* allele from strain PB384 (a kind gift from P. Bissichia and D. Sherratt) was first introduced into the BW27783 background, followed by the *frt-aph-frt-Ypet-dnaN* allele from strain RRL190 (a kind gift from R. Reyes-Lamothe) (1). After each P1 transduction, the kanamycin resistance gene was removed by expressing Flp recombinase from plasmid pCP20 (2).

To follow ribosomes and chromosome dynamics, the strains CA4 (*rpsB::rpsB-mCherry-frt*, *hupA::hupA-mYpet-frt*) and CA7 (*rplI::rplI-mCherry-frt*, *hupA::hupA-mYpet-frt*) were constructed by λ-Red engineering (3) and by P1 transduction. For chromosomal replacement of *rpsB* (encoding 30S ribosomal protein S2) with its fluorescence derivative, we used primers carrying tails with identical sequences to the *rpsB* chromosomal locus (see Table S2 for primer sequences) and a plasmid pROD62 (a kind gift from R. Reyes-Lamothe) carrying a copy of mCherry followed by a kanamycin resistance cassette flanked by frt-sites (*mCherry-frt-aph-frt*) as PCR template. The linker sequence (CAGGAAAGGCGACAGGAG) used between *rpsB* and *mCherry* sequence was the same as for the *rpsB* construct published earlier (4). The resulting PCR product was transformed by electroporation into a strain carrying the λ-Red-expressing plasmid pKD46 in BW25113 background. Colonies were selected by kanamycin resistance, verified by fluorescence microscopy, and by PCR using primers annealing to regions flanking the *rpsB* gene. After verification, the resulting construct was transferred by P1 transduction to BW27783 cells, and the kanamycin resistance gene was eliminated by plasmid pCP20. Next, the *hupA-mYpet* gene fusion from strain JM52 was transferred into the strain by P1 transduction, and the kanamycin cassette was removed using plasmid pCP20, resulting in a final strain CA4.

To construct strain CA7, we followed the approach of Chai *et al.* (5), in which no linker sequence was added between *rplI* (encoding L9 of 50S subunit) and *mCherry* sequences. The termination codon of *rplI* in the *E. coli* BW25113 chromosome was replaced with a DNA sequence containing *mCherry* and a kanamycin resistance cassette using λ-Red engineering, as described above. After verifying the strain by PCR and microscopy, the *rplI-mCherry* construct was P1 transduced into BW27783. Additionally, hupA-mYpet was introduced via P1 transduction, as previously described for CA4, resulting in strain CA7. *hupA-mYpet* originated from strain JM52, which was constructed by λ-Red engineering. To construct JM52, the *linker-mYpet-frt-aph-frt* cassette was amplified from a template plasmid pJM21 (Mannik lab) using long primers with identical sequences to the *hupA* chromosomal locus and electroporated into MG1655/pKD46 strain. A flexible peptide with sequence SAGSAAGSGEF was used as a linker between the mYpet and HupA protein.

For *E. coli* strain engineering, the cells were grown in lysogeny broth (LB) with appropriate selective antibiotics.

**Cell preparation and culturing in microfluidic devices**

All bacterial strains were streaked onto agar plates containing M9 minimal salts supplemented with 2 mM magnesium sulfate (MilliporeSigma, MO), trace metals mixture (Teknova Inc., CA, #T1001) and 0.3% glycerol (Fisher Scientific, NH) as the carbon source for slow growth measurements. For moderately fast growth, glycerol was replaced by 0.5% glucose (MilliporeSigma), and 0.2% casamino acids (Fisher Scientific). A day before the experiment, a single colony was inoculated into 3 ml of M9 minimal salts media (Teknova Inc., CA) supplemented with the appropriate carbon source and additives described above. For microscopy experiments, cells were grown in a liquid medium to an $OD_{600}$ of ~0.07. The culture was then concentrated ~100x by centrifugation in the presence of 75 µg/ml of Bovine Serum Albumin (BSA; MilliporeSigma) to minimize cell clumping. The resulting concentrated cell suspension was used to inoculate a polydimethylsiloxane (PDMS)-based microfluidic mother machine device. The latter was first wetted with media, after which 2 µl of resuspended concentrated culture was pipetted into its main flow channel. The cells were left to populate the dead-end channels for 0.5-1 hours before connecting the tubing for constant media flow. Media flow was maintained throughout the experiment using a NE-1000 Syringe Pump (New Era Pump Systems, NY) at a rate of 6 µl/min. The cells were grown overnight to achieve steady-state growth before imaging. All cells were cultured, and experiments were conducted at 28°C.

**Antibiotic treatments**

For treatments with rifampicin or/and cephalexin two 10-ml syringes were prepared with fresh growth medium. To one syringe antibiotic(s) were added. The two syringes were mounted on two separate NE-1000 Syringe Pumps. Both syringes were connected via tubing to a T-connector. The tubing from the T-connector connected further to a single inlet hole in the microfluidic mother machine chip. Before applying antibiotics the cells were imaged in regular M9 media. After 8 hours of imaging (3-4 doublings), the first pump with the regular media was turned off, and the second syringe pump containing media with antibiotics was turned on without interrupting imaging. The two pumps were controlled by a custom LabVIEW program. The concentration for rifampicin was 300 µg/ml (MilliporeSigma) and for cephalexin 20 µg/ml (MilliporeSigma). The same concentrations were used also when both antibiotics were present in the growth media at the same time. Cells were imaged for ~10-12 hours in the media with antibiotics.

There was about a 10-minute delay from turning on the syringe pump with the antibiotic medium to the antibiotic reaching cells in the microfluidic chip. This delay varied slightly from one measurement to another. To account for this delay in measurements with rifampicin treatment we used an abrupt shrinkage of the nucleoid dimensions that has been interpreted as the arising from breakage of transertion linkages (45, 47) as the start time, $T_{ab}$, for these measurements.

**Fabricating and assembling mother machine devices**

The assembly of mother machine microfluidic devices was based on soft-lithography of PDMS following a previously described protocol (6). To define channels in PDMS a 4″ silicon (Si) wafer mold was used. The fabrication protocol for the mold can be found in the section below. In the first step of assembly, PDMS elastomer (Sylgard 184 kit, Dow Corning, MI) was mixed in a 9:1 base/linker weight ratio and then degassed. Si mold was placed into a plastic holder, and the elastomer was then poured over the mold, forming about 5 mm thick layer. The elastomer was then further degassed. The PDMS and the mold were baked at 90ºC for 20 min in a convection oven and then left in the oven for at least two more hours as the oven cooled down from 90ºC. Solidified PDMS was then peeled from the Si mold and individual patterns were cut out from it, and access holes to the main channels in each pattern were punched using a biopsy needle. These pieces were subsequently bonded to coverslips. For bonding #1.5 coverslips were cleaned in isopropanol (both from Fisher Scientific) by sonication and then treated in $O_2$ plasma at 200 mTorr for 70 s. The PDMS elastomer piece and glass coverslips are additionally treated in $O_2$ plasma for 3-5 s before bonding. After bonding, the chips were kept at room temperature for at least 12 hours before starting live cell measurements.

**Fabrication of silicon molds for mother machine devices**

The fluidic circuitry in each microfluidic chip consists of a main channel serving media flow and 2400 side channels connected to the main channel. The side channels are populated by bacteria that are imaged in experiments. Silicon (Si) molds were used to imprint these channels in PDMS. The molds were fabricated using a combination of electron beam lithography (EBL), photolithography and reactive ion etching (RIE) using a new protocol described below.

In the first step, PMMA electron beam resist (495K MW in 4% Anisole) was spin-coated onto a clean 4” silicon wafer at 6000 rpm for 45 s. The wafer was then soft-baked at 180°C for 2 min. JEOL 8100 FS Electron Beam Lithography System set to a beam current of 2 nA was used to pattern the mother machine side channels. After exposure, the patterned wafer was developed in MIBK: IPA (1:3) for 1 min, then rinsed with IPA and dried with nitrogen gas. A brief descumming step in Oxford Plasmalab System 100 Reactive Ion Etcher for 6 s was used to remove the residual resist from the exposed areas.

A thin chromium (Cr) layer with a thickness of 0.150 kÅ is deposited on the wafer using Thermionics VE-240 e-beam evaporator. Next, Cr and PMMA resist were lifted off by sonicating the wafer for 2 min in acetone. This was followed by rinsing the wafer with IPA, water, and drying it with nitrogen gas. The patterned wafer was then baked at 180°C for 1 hour to further stabilize the Cr layer.

For photolithography, the SU-8 2015 photoresist was spin-coated onto the Cr-patterned wafer at 2000 rpm for 45 s. The wafer was pre-baked at 65°C for 1 min and then at 95°C for 6 min to prepare the resist for exposure. The wafer was exposed to UV light at an energy dose of 150 mJ/cm² using SUSS MicroTech UV Contact Aligner. The wafer was baked at 65°C for 1 min and 95°C for 6 min post-exposure. The exposed patterns were developed in SU-8 developer for 2.5 min, rinsed with the developer and IPA, and dried with nitrogen gas. A final baking step at 180°C for 15 min was used to reflow SU-8. The resulting main channel was about 18 μm high and 200 μm wide.

To create the side channel reliefs, the wafer was etched in Oxford Plasmalab System 100 Reactive Ion Etcher using $C_4F_8$, $SF_6$ and Ar gases. The resulting side channels were ~1.2 μm high.

In the final step, the wafer was silanized using 97% Trichloro(1H,1H,2H,2H-perfluorooctyl)silane (MilliporeSigma) in a desiccator overnight.

**Microscopy**

Bacterial imaging was conducted using a Nikon Ti-E inverted epifluorescence microscope (Nikon Instruments, Japan) equipped with a 100X (NA = 1.45) oil immersion phase contrast objective. Images were acquired with an iXon DU897 EMCCD camera (Andor Technology, Ireland) and recorded via NIS-Elements software (Nikon Instruments, Japan). Fluorophores were excited with a 200 W Hg lamp that was attenuated by an ND4 and ND8 neutral density filters. Chroma 41004 and 41001 filter cubes (Chroma Technology Corp., VT) were used in capturing fluorescence from mYpet and mCherry fluorophores, respectively. A motorized stage (Prior Scientific Inc., MA) and a Nikon Perfect Focus® system were utilized throughout time-lapse imaging. Images were obtained at a 4 min frame rate. The typical exposure times were 300 ms at an EM gain of 200 for mYpet; 300 ms at an EM gain of 200 for mCherry, and 400 ms without EM gain for the phase images.

**Image and data analysis**

MATLAB, along with the Image Analysis Toolbox, Optimization Toolbox and DipImage Toolbox (http://www.diplib.org/) and Python 3.11.4 with torch and torchvision packages (https://pytorch.org/) were used for image analysis. First, the phase contrast images were correlated on a frame-by-frame basis to determine subpixel shifts between different frames. From shift-corrected images, images of individual channels were cropped. Bacterial cells from the latter images were segmented using a U-Net convolutional neural network (CNN), referred to as the Omnipose (7). This model was previously re-trained on images from microscopy setup used in this work (8). Using a custom MATLAB script (8), the cell masks from segmentation were used to assemble cell lineages that stored the coordinates of cell centerlines and cell lengths ($L_{cell}$).

**DNA and ribosome density distributions**

The DNA, ribosome, and replisome density distributions, $I(x,t)$, along the cell length (x-axis) at time $t$ were calculated based on fluorescence intensities of DNA (HupA-mCherry, HupA-mYpet), ribosome (RpsB-mCherry, RplI-mCherry) and replisome labels (Ypet-DnaN), respectively. To calculate these distributions, fluorescence intensities from images along an 11-pixel (1.2 $\mu m$) wide line perpendicular to the long axis of the cell (y-axis) were sampled and integrated for each length bin along the cell length (x-axis). The perpendicular lines were centered at the cell's long symmetry-axis. The latter was defined by a segmentation mask. The intensity was then normalized per pixel value, and the image background level was subtracted from this value. We refer to this way processed signal as the uncompensated density along the long axis of the cell, $I_u(x,t)$.

The cell level concentrations of DNA (HupA-mCherry, HupA-mYpet) and ribosome labels (RpsB-mCherry, RplI-mCherry), calculated as $c(t) = \sum_i I_u(x_i,t)\,/L_{cell}$, varied by less than 5% during the cell cycle in normal growth conditions. However, during rifampicin treatment, these concentrations increase due to the maturation of the fluorophores and cessation of cell growth (9). To compensate for this effect, we divided $I_u(x,t)$ by $c(t)$. For consistency, we applied the same compensation also for measurements without rifampicin treatment although in the latter cases, this resulted in the renormalization of $I_u(x,t)$ by effectively a constant value and had no significant impact on results because these were presented in arbitrary units (a.u.). The normalized densities $I(x,t) = I_u(x,t)/c(t)$, are referred to as density distributions. Note that the arbitrary units used for density distributions are consistent from one measurement to another.

The density distributions were compiled over time to construct population-averaged kymographs. In these kymographs, the horizontal axis represents the time, and the vertical axis is the x-axis of the cell (the long axis of the cell). The zero of the vertical axis corresponds to the center of the cell.

**Replication and cell cycle-dependent kymographs**

The population-averaged kymographs are created by averaging $I(x,t)$ from individual cells at fixed cell cycle ages over cell population. The replication age, $a_r$, is calculated for each cell as $a_r = t - T_{ri}/(T_{rt} - T_{rt})$ where $T_{ri}$ is the time for DNA replication initiation and $T_{rt}$ for termination. The cell age in the division cycle, which is referred to as the cell age, $a$, is calculated as $a = t/T_d$ where $T_d$ is the duration of the cell cycle. All times, including $t$, are measured from cell birth. Since cells at a given cell or replication age are of different lengths, the cell coordinates, $x$, are scaled based on an average length elongation curve. For the division cycle kymographs, the average length elongations curve is $L_{cell,av}(a) =< L_{cell,b} > 2^a$ where $< L_{cell,b} >$ is the population average cell length at birth as determined from the same measurement. For the replication cycle kymographs, the average length elongation curve is $L_{cell,av}(a_r) =< L_{cell,Tri} > 2^{a_r}$ where $< L_{cell,Tri} >$ is the measured population average cell length at the initiation of DNA replication. The new coordinate values for each cell, $i$, at the age $a$ are $x' = xL_{cell,av}(a)/L_{cell,i}(a)$ where $L_{cell,i}(a)$ is the length of this cell at age $a$. The $I(x',a)$ values are then resampled at a fixed spatial grid and the resulting curves are averaged over different cells. The origin of the coordinate axis is chosen at the center of the cell. The most negative value of the coordinate corresponds to the old pole of the cell.

**Kymographs aligned relative to specific times**

We also created kymographs for the entire cell population or groups of cells aligned to a specific event at the measurement time $T_x$ such as the start of antibiotic treatment ($T_{ab}$), formation of the nucleoid density minimum ($T_{min}$) and so on. The time zero of the kymographs thus corresponds to this event. To account for variation in the cell lengths, cell lengths were normalized to one for each cell at every time point. In these kymographs, the y-axis corresponds to the normalized length of the cell, with the center set to zero and the axis values varying from -0.5 to 0.5. -0.5 value corresponds to the old pole of the cell. Due to the normalization of cell lengths, there is no cell elongation apparent in kymographs, even when cell filament as they do in measurements with cephalexin treatment (Fig. 3).

**Quantification of midcell amount and the excess midcell fraction**

The normalized midcell amount of ribosomes and DNA as a function of time, $t$, was calculated as the total signal intensity of corresponding fluorescence labels in a 0.75 μm wide band centered at the midcell using:

$$N_{mid}(t) = \sum_{x=x_0-br}^{x_0+b} I(x,t)$$

Here $x_0$ and $br$ denote the cell center coordinate and half-width of the band, respectively. $br$ was taken 3 pixels so that the total width of the band was 7 pixels (0.75 μm).

The relative excess fraction is calculated as:

$$\frac{\Delta N(t)}{N(t)} = \frac{N_{mid}(t) - N_{1/4}(t)}{N_{tot}(t)} = \frac{\left(\sum_{x=x_0-b}^{x_0+br} I(x,t) - \sum_{x=x_{0.25}-br}^{x_{0.25}+b} I(x,t)\right)}{\sum I(x,t)}$$

where $x_{0.25}$ is the coordinate of the cell's quarter position and the denominator is the total integrated intensity from the cell.

**Quantification of nucleoid dimensions**

To determine the dimensions of the nucleoids HupA-mCherry (or HupA-mYpet) signal from a single cell was fitted with multiple generalized Gaussians $(gG)$ functions:

$$gG_{fit}(x) = A_0 + \sum_i A_i \exp\left(-\frac{1}{2}\left(\frac{x - x_{0,i}}{\sigma_i}\right)^{2\beta_i}\right) (1)$$

Here $A_0$ is the signal background, $x$ is the coordinate along the long axis of the cell, is $x_{0,i}$the centroid for individual nucleoid $i$, $A_i$ is the signal amplitude, and parameters $\sigma_i$ and $\beta_i$ determine the width and the shape of the nucleoid density distribution. The fit was constrained so that $0.9 < \beta_i < 10. \beta_i = 1$ corresponds to a regular Gaussian function and $\beta_i > 1$ to functions that are flatter than the Gaussians. We used an additional generalized Gaussian on each side of the fitted cell to account for the neighboring cells as they grew in tight files in microfluidic channels. To better account for the "leakage" of the fluorescence from the two adjacent cells, we extended the intensity profiles from the cell of interest by 0.75 mm (7 pixels) to these neighbor cells. A generalized Gaussian function term in the sum above was used to account for fluorescence from these two neighboring cells (SI Fig. S22). The fit function thus had at least three terms with the generalized Gaussian functions. Additional generalized Gaussian terms were used for multi-lobed nucleoids. The number of terms (above three) in the sum was determined using the Akaike information criterion (10). For cells in slow growth conditions in the M9 glycerol medium, there was only one extra term possible, while for cells in the M9 glucose+CAS medium, three additional terms were possible.

Based on the fitting, we determined the total nucleoid extent, $L_{extent}$, the sum of nucleoid lengths, $L_{sum}$, and the peak distance, $D_{peak}$.

$L_{extent}$ is defined as the distance between the outermost inflection points ($\frac{d^2I}{dx^2} = 0$) of the fitted signal $I(x)$ that excluded contributions of the neighboring cells. $\frac{d^2I}{dx^2}$ was calculated numerically from the fitted data using the same grid as for the experimental data. The zero crossing of $\frac{d^2I}{dx^2}$ was found with a sub-pixel accuracy by a linear interpolation of two data points closest to the zero-crossing in the fitted signal.

$L_{nuc,sum}$ is calculated as the sum of fitted nucleoid lengths, each defined as the full width at the half maximum:

$$L_{nuc,sum} = \sum_i 2\sigma_i \cdot (\ln\ 4)^{\frac{1}{2\beta_i}}$$

The sum includes only terms from the cell of interest and not from the neighboring cells.

$D_{peak}$ was calculated as a distance between the centers of two generalized Gaussians that were closest to the cell center ($D_{peak} = x_{0,imiddle} - x_{0,imiddle+1}$). In most cases, except cephalexin treatment measurements presented in Fig. 3, there were only two nucleoids, and, in this case, $D_{peak} = x_{0,1} - x_{0,2}$.

**Quantification of ribosome distributions**

The ribosome distributions along the cell's long axis (RpsB-mCherry or RplI-mYpet) were used to quantify the excess ribosome fraction at the midcell and the width of the ribosome distribution that localizes to the midcell. The excess ribosome fraction at the midcell (Fig. 3J) corresponds to the excess fraction of ribosomes that are present at midcell as compared to other locations along the cell body except poles. This fraction was determined by two independent methods. In one approach, the integrated signal intensity in 0.75 μm (7 pixel) wide band centered at cell middle and the quarter positions were calculated as explained in '*Quantification of midcell amount and the excess midcell fraction*' in above.

In the other approach, the excess fraction of ribosomes at midcell was determined from the least-square fitting of the ribosome density signal from the cell. The fitting procedure was similar to the one for DNA density distributions, where the signal was extended to neighboring cells and a different number of generalized Gaussians, given by eq. (1), were used to model the signal (SI Fig. S22). The signal originating from each of the two neighboring cells was accounted for by a generalized Gaussian term in the fit function ($i = 3,4$). In addition to these and background terms, the ribosome signal from the cell of interest was represented by either one or two generalized Gaussian functions. The choice between one or two generalized Gaussian terms was again made based on the Akaike information criterion. One of these two generalized Gaussians represented the near uniform distribution of ribosomes along the long axis of cells ($i = 1$) while the other one to the ribosomes localized at midcell ($i = 2$). The parameter $\sigma_2$ of the latter generalized Gaussian was constrained during the fit. The amplitude for this term, $A_2$, could be negative to account for the lower density of ribosomes during the constriction. Based on this fitting, the integrated signal intensity $I_{\Sigma,\mathrm{I}}$ for each of the two generalized Gaussian terms that corresponded to the strain of interest were calculated from:

$$I_{\Sigma,\mathrm{i}} = \sigma_i A_i \Gamma\left(\frac{0.5}{\beta_i}\right)/\beta_i$$

where $\Gamma$ is the gamma function. Based on these two intensities, the excess midcell fraction from the fit was calculated as

$$\left(\frac{\Delta N(t)}{N(t)}\right)_{fit} = \frac{I_{\Sigma,2}}{I_{\Sigma,1} + I_{\Sigma,2}}$$

The width of the ribosome midcell accumulation was calculated as $W = 2\sigma_1 \cdot (\ln\ 4)^{\frac{1}{2\beta_1}}$.

**Pearson correlation coefficient**

In calculations of the Pearson correlation coefficient, $R,$ between DNA and ribosome densities along the cell's long axis, 0.64 μm (6 pixels) long region from each cell end was excluded from the calculations. Pearson correlation coefficient was calculated for each time frame for every cell, and then the resulting values were extrapolated to equally spaced cell age bins. The data in each age bin from a population of cells were then averaged. s.e.m. was calculated for each age bin and are shown as error bars in the plots.

**Determining the replication period**

To determine replication period replication initiation ($T_{ri}$) and termination ($T_{rt}$) time points were manually marked on Ypet-DnaN (beta clamp) kymographs of individual cells, as shown in SI Fig. S2. $T_{ri}$ and $T_{rt}$ markings tracked the beginning and end of the midcell high-density region of the Ypet-DnaN signal.

**Correcting birth and division timings**

To determine the correct division time/frame ($T_d$), $N_{mid}(t)$ from phase contrast images were used. In the majority of cells, $N_{mid}(t)$ curve showed a cusp at the time of division. In this cusp, the signal that increased in time plateaued. The corrected $T_d$ was then also used to correct the birth times/frames of the daughter cells.

**Determining $A_{min}$ and $T_{min}$ from DNA and ribosomal distributions**

To determine the depth of the local minimum of DNA (ribosomal) midcell density distribution, $A_{min}$, ($A_{min,ribo}$) and the time when a minimum in the nucleoid (ribosomal) density forms, $T_{min}$,($T_{min,ribo}$) we used a previously described approach based on thresholding (42). In this approach, the DNA (ribosomal) density profile for each time frame is first normalized so that the maximum value is unity. All the local minima in this curve were found except those at cell poles. The most prominent minimum that was no more than 0.4 $\mu m$ from the cell center was recorded if the value of this minimum was less than 0.95 in normalized units. Different threshold values could be set using this automated approach. The chosen value of 0.95 represents a trade-off between recording spurious minima due to noise in the signal and missing a true minimum in DNA (ribosomal) density (11).

The recorded $A_{min}$ values were used to find $T_{min}$. If $A_{min}$ was zero in two or more consecutive frames starting from the division frame, then that last recorded frame when it was present determined the $T_{min}$ for this cell.

**The coupled DNA and ribosome dynamics model**

**Model Overview.** Our coupled DNA and ribosome dynamics model is derived using non-equilibrium considerations and incorporates the major reactions involved in the Central Dogma processes. We consider the volume fractions of the DNA, ribosomal subunits, and the assembled polysomes represented by $\phi_{\text{DNA}}$, $\phi_{\text{sub}}$, and $\phi_{\text{poly}}$, respectively. The volume fractions $\phi_i \equiv \phi_i(x,t)$ depend on the position $x$ along the long axis of the cell and the time $t$. Our model includes the following reactions: (1) the formation of a polysome (polyribosome), (2) the disassembly of a polysome, (3) the assembly of ribosomes into polysomes, and (4) the removal of ribosomes from the polysome as they complete translation. Polysomes are formed from ribosomal subunits in the vicinity of DNA. The disassembly of polysomes results from the degradation of the mRNA on which the ribosomes assemble. All species interact with each other via excluded volume interactions. Our model also accommodates cell growth, and we fix constant volume fractions of each species $i$, $\phi_i$, during cell growth (balanced growth).

The model treats mRNA in an effective manner, assuming the molecule to be part of polysomes. We also do not consider soluble proteins and nucleoid-associated proteins. We assume that their effects on DNA can be incorporated in an effective DNA self-energy (12) and in a renormalization (overall decrease) of the polysome and DNA diffusion coefficients. We also treat polysomes with a single concentration field, $\phi_{\text{poly}}$, which captures the entire polysome distribution. We assume that, on average, each polysome has $N_{\text{ribo}} = 10$ (13,14) ribosomes attached and diffuses with some average diffusion coefficient $D_{\text{poly}}$. Such a simplification was also recently applied in a similar study (15).

The interactions between the various species are based on excluded volume considerations and Flory theory. As discussed in our previous work (13), polysomes are too large to diffuse through DNA supercoils so their excluded volume with DNA is treated differently from the DNA and free ribosome interaction. The various equilibrium properties of these interactions are captured by an effective, coarse-grained free energy $\mathcal{F}$, which we discuss in the following section. We will then describe the reactions in further detail. We then estimate model parameters, summarize all equations, and explain the implementation of the model.

**The equilibrium free energy.** Assuming a local equilibrium and slow variations of the volume fractions of the species (DNA, polysomes, ribosomal subunits), we expect our system to be described by a free energy $\mathcal{F} \equiv \mathcal{F}[\phi_{\text{DNA}}, \phi_{\text{sub}}, \phi_{\text{poly}}]$, which can be written as a functional of the various volume fractions. The system evolves toward a minimum of $\mathcal{F}$ via the flow of each species $i$ determined by the gradient of the corresponding chemical potential $\mu_i$. The latter is given by the functional derivative $\mu_i = v_i \delta\mathcal{F}/\delta\phi_i$, with $v_i$ the molecular volume of the species. The corresponding currents $\mathbf{J}_i$ for species $i$ are given by

$$\mathbf{J}_i = -\beta D_i \phi_i \nabla \mu_i = -\beta D_i \phi_i v_i \nabla \frac{\delta\mathcal{F}}{\delta\phi_i}, \tag{1}$$

where $\beta = (k_{\mathrm{B}}T)^{-1}$ is the inverse temperature parameter and $D_i$ is the diffusion coefficient. We will assume that the time evolution of the volume fraction $\phi_i$ is governed by a simple relaxation of the free energy $\mathcal{F}$ according to the current in Eq. (1), neglecting any hydrodynamic or elastic contributions to the dynamics. Without reactions, the various species are conserved within the cell, so the volume fractions obey the continuity equation:

$$\partial_t \phi_i + \nabla \cdot \mathbf{J}_i = 0. \tag{2}$$

In the following subsections, we introduce all of the terms in $\mathcal{F}$, which are based on the free energy model described in (13).

**I. DNA interactions.** DNA is a complex polymeric molecule decorated with various proteins, and it has a hierarchical structure, including supercoils. Our model neglects this complexity and considers DNA as a concentration (volume fraction) field. We envision the concentration field to arise from a collection of DNA "monomers" where monomer size is about the size of a typical DNA supercoil. As such DNA in our model lacks chain connectivity. The same approach has been used previously by Miangolarra *et al.* (16) and Papagiannakis *et al.* (15).

The complex self-interactions within the DNA yield an effective power-law behavior in the free energy as a function of the nucleoid volume, $V_{\mathrm{nuc}}$ (13,17): $g\left(\frac{V_{\mathrm{DNA},0}}{V_{\mathrm{nuc}}}\right)^{\alpha}$, where $V_{\mathrm{DNA},0} \approx 27\ \mu\mathrm{m}^3$ is the relaxed DNA volume, and $g \approx 147$ is related to the degree of cross-linking with the numerical values determined from *in vitro* measurements of liberated *E. coli* nucleoids. The exponent $\alpha$ is related to the Flory exponent as $\nu = (1+\alpha)/3\alpha$ (13). We now assume that such a power law holds on smaller length scales within the nucleoid. Then, given that the volume fraction of DNA, $\phi_{\mathrm{DNA}}$, vanishes outside of the nucleoid, and satisfies $\phi_{\mathrm{DNA}} \approx V_{\mathrm{DNA}}/V_{\mathrm{nuc}}$ inside the nucleoid (where $V_{\mathrm{DNA}}$ is the molecular volume of DNA), the self-energy can be written as an integral over the cell interior:

$$\beta\mathcal{F}_{\mathrm{DNA}} = g \int \mathrm{d}x \left(\frac{V_{\mathrm{DNA},0}\phi_{\mathrm{DNA}}}{V_{\mathrm{DNA}}}\right)^{\alpha} \frac{\phi_{\mathrm{DNA}}}{V_{\mathrm{DNA}}} = g \int \mathrm{d}x\, \frac{V_{\mathrm{DNA},0}^{\alpha}}{V_{\mathrm{DNA}}^{\alpha+1}} \phi_{\mathrm{DNA}}^{\alpha+1}. \tag{3}$$

Next, we consider the interaction term between DNA and polysomes, which we assume is captured by the leading order term in the virial expansion of the excluded volume interaction:

$$\beta\mathcal{F}_{\mathrm{DNA-poly}} = \int \mathrm{d}x\, \frac{\phi_{\mathrm{poly}}}{v_{\mathrm{poly}}} \frac{B_{\mathrm{DNA\text{-}poly}}}{V_{\mathrm{DNA}}} \phi_{\mathrm{DNA}}, \tag{4}$$

where $B_{\text{DNA-poly}}$ is the excluded volume between a single polysome and the DNA (estimated below), and $v_{\text{poly}}$ is the molecular volume of a polysome. Similarly, the interaction term between DNA and free ribosomal subunits is:

$$\beta\mathcal{F}_{\text{DNA-sub}} = \int \mathrm{d}x \frac{\phi_{\text{sub}}}{v_{\text{sub}}} \frac{B_{\text{DNA-sub}}}{V_{\text{DNA}}} \phi_{\text{DNA}}, \tag{5}$$

with $B_{\text{DNA-sub}}$ the DNA-subunit excluded volume. The continuity equation for the DNA volume fraction thus reads:

$$\begin{aligned} \partial_t \phi_{\text{DNA}} &= \beta D_{\text{DNA}} v_{\text{DNA}} \nabla \cdot \left[ \phi_{\text{DNA}} \nabla \left( \frac{\delta \mathcal{F}_{\text{DNA}}}{\delta \phi_{\text{DNA}}} \right) \right] \\ &= D_{\text{DNA}} v_{\text{DNA}} \nabla \cdot \left[ \frac{g\alpha(\alpha+1) V_{\text{DNA},0}^{\alpha}}{V_{\text{DNA}}^{\alpha+1}} \phi_{\text{DNA}}^{\alpha} \nabla \phi_{\text{DNA}} + \frac{B_{\text{DNA-poly}}}{v_{\text{poly}} V_{\text{DNA}}} \phi_{\text{DNA}} \nabla \phi_{\text{poly}} + \frac{B_{\text{DNA-sub}}}{v_{\text{sub}} V_{\text{DNA}}} \phi_{\text{DNA}} \nabla \phi_{\text{sub}} \right], \end{aligned} \tag{6}$$

where $v_{\text{DNA}}$ is the molecular volume of the diffusive DNA "monomer," which we take to be a DNA supercoil. The volume $v_{\text{DNA}}$ can be calculated by modeling the DNA supercoil as a cylinder. Then,

$$v_{\text{DNA}} \approx \frac{\pi L_{\text{DNA}} d_s^2 \sin \delta}{8 N_s} \approx 6.65 \times 10^{-5} \mu\text{m}^3, \tag{7}$$

where $L_{\text{DNA}} = 1.6$ mm is the length of the chromosomal DNA, $\delta = 52°$ is the plectoneme opening angle, $N_s = 6700$ is the number of DNA supercoiling segments and $d_s = 30$ nm is the diameter of the DNA supercoiling segment (13). The total volume occupied by the DNA is estimated as $V_{\text{DNA}} = 0.1\,\mu\text{m}^3$. In addition, we assume that DNA supercoil segments diffuse 10 times slower than the polysomes so that $D_{\text{DNA}} = 0.1 D_{\text{poly}}$. Finally, the virial coefficients can be calculated by estimating the corresponding excluded volumes:

$$B_{\text{DNA-poly}} = \frac{\pi}{8} L_{\text{DNA}} \left(d_s + 2R_g^{\text{poly}}\right)^2 \sin\delta + \frac{\pi}{6} N_s \left(d_s + 2R_g^{\text{poly}}\right)^3 \approx 8.47\ \mu\text{m}^3 \tag{8}$$

$$B_{\text{DNA-sub}} = \frac{\pi}{4} L_{\text{DNA}} (d_{\text{DNA}} + a_{\text{sub}})^2 \approx 0.6\ \mu\text{m}^3, \tag{9}$$

where $d_{\text{DNA}} = 2$ nm is the diameter of the double helix DNA, $a_{\text{sub}} = 20$ nm is the ribosomal subunit diameter, and $R_g^{\text{poly}} \approx 35$ nm is the polysome radius of gyration (13). We thus estimate the molecular volumes of the ribosome and polysome as $v_{\text{sub}} = \frac{\pi}{6} a_{\text{sub}}^3 \approx 4.2 \times 10^{-6}\ \mu\text{m}^3$ and $v_{\text{poly}} = \frac{4\pi}{3} \left(R_g^{\text{poly}}\right)^3 \approx 1.8 \times 10^{-4}\ \mu\text{m}^3$, respectively. Note that in this treatment, we consider the polysome as a polymeric chain with a certain radius of gyration, which depends on the number of ribosomes $N_{\text{ribo}}$ comprising the polysome, which here we hold fixed at $N_{\text{ribo}} = 10$. The radius of gyration corresponds to a polymer under good solvent conditions, as $R_g^{\text{poly}} > a_{\text{sub}} \sqrt{N_{\text{ribo}}/6} \approx 26$ nm.

**II. Polysome interactions.** Unlike DNA, polysomes can move freely throughout the cell, so we consider the entropy of mixing in our free energy formulation. Assuming a small volume fraction $\phi_{\text{poly}} \ll 1$, we have

$$\beta\mathcal{F}_{\text{poly}}^{\text{ent}} = \int \mathrm{d}x \, \frac{\phi_{\text{poly}}}{v_{\text{poly}}} \left(\ln \phi_{\text{poly}} - 1\right), \tag{10}$$

with a corresponding diffusive current

$$\mathbf{J}_{\text{poly}}^{\text{ent}} = -D_{\text{poly}} v_{\text{poly}} \phi_{\text{poly}} \nabla \left[\frac{1}{v_{\text{poly}}}\left(\ln \phi_{\text{poly}} - 1\right) + \frac{1}{v_{\text{poly}}}\right] = -D_{\text{poly}} \nabla \phi_{\text{poly}}. \tag{11}$$

The polysome is a chain of ribosomes assembled on an mRNA. We will follow previous results (18) on the excluded volume interactions between two polymers, which yields a free energy of the form:

$$\beta\mathcal{F}_{\text{poly}-\text{poly}} = \int \mathrm{dx} \frac{\phi_{\text{poly}}}{\gamma v_{\text{poly}}} \left(\frac{\phi_{\text{poly}}}{0.69\phi_{\text{poly}}^{\text{ov}}}\right)^{\gamma}, \tag{12}$$

where $\phi_{\text{poly}}^{\text{ov}} = 1.86$ is the overlap volume fraction of polysomes, and $\gamma = 1.309$ is an exponent derived from a renormalization group calculation (18). The corresponding current is thus

$$\mathbf{J}_{\text{poly}-\text{pol}} = -D_{\text{poly}} v_{\text{poly}} \phi_{\text{poly}} \nabla \left[\frac{\gamma + 1}{\gamma v_{\text{poly}}} \left(\frac{\phi_{\text{poly}}}{0.69\phi_{\text{poly}}^{\text{ov}}}\right)^{\gamma}\right] = -D_{\text{poly}} \frac{(\gamma + 1)\phi_{\text{poly}}^{\gamma}}{\left(0.69\phi_{\text{poly}}^{\text{ov}}\right)^{\gamma}} \nabla \phi_{\text{poly}}. \tag{13}$$

The polysome-subunit interactions are again treated using the virial expansion:

$$\beta\mathcal{F}_{\text{poly}-\text{sub}} = \int \mathrm{dx} \frac{\phi_{\text{poly}}\phi_{\text{sub}}}{v_{\text{poly}} v_{\text{sub}}} B_{\text{poly-sub}}. \tag{14}$$

The virial coefficient of polysome-subunit interaction is estimated as

$$B_{\text{poly-sub}} = \frac{\pi}{6} N_{\text{ribo}} (2a_{\text{sub}})^3 \approx 3.35 \times 10^{-4}\ \mu\text{m}^3, \tag{15}$$

where we assume the free ribosomal subunits interact with the ribosomes attached to the mRNA individually. The subunits diffuse more quickly than the polysomes, and we estimate that $D_{\text{sub}} = 3.5 D_{\text{poly}}$, based on the ratio of the linear sizes of the two species.

Combining all these interactions (along with the DNA-polysome interaction in Eq. (4)), the continuity equation for the polysome volume fraction reads:

$$\partial_t \phi_{\text{poly}} = D_{\text{poly}} \nabla \cdot \left[\nabla \phi_{\text{poly}} + \frac{(\gamma + 1)\phi_{\text{poly}}^{\gamma}}{\left(0.69\phi_{\text{poly}}^{\text{ov}}\right)^{\gamma}} \nabla \phi_{\text{poly}} + \frac{B_{\text{DNA-poly}}}{V_{\text{DNA}}} \phi_{\text{poly}} \nabla \phi_{\text{DNA}} + \frac{B_{\text{poly-sub}}}{v_{\text{sub}}} \phi_{\text{poly}} \nabla \phi_{\text{sub}}\right]. \tag{16}$$

The effective diffusion coefficient of polysomes is set to $D_{\text{poly}} = 0.01\ \mu\text{m}^2/s$, which is within an order of magnitude of the measured value from single-molecule tracking (4). We expect smaller values of the diffusion coefficients due to additional crowding from other macromolecules not considered here, especially the proteins.

**III. Subunit interactions.** For simplicity, we consider subunits forming 70S ribosome particles in the cytosol instead of individual 30S and 50S units. This approach simplifies the treatment of reactions that involve ribosome subunits. In the reactions we consider (discussed in detail later), the binding and unbinding of one subunit from mRNA is quickly followed by the binding and unbinding of the other. So, the subunits effectively act as one 70S ribosome particle.

Replacing ribosome subunits with a single 70S particle somewhat alters their excluded volume interactions with the other molecules, but we expect the effect to be small because only a small fraction of ribosome subunits is free (20 %), and they are distributed throughout the whole cell volume. So, the subunits act as a dilute gas. We include their effects in the first-order virial coefficients, where the difference between the 30S and 50S subunits and a single 70S particle is small.

We consider the virial expansion of a hard-sphere gas for the subunit-subunit self-interaction. The latter is well-approximated by the Carnahan-Starling equation of state, which we expand around small $\phi_{\text{sub}}$. Keeping just the first two leading-order terms, the corresponding current is

$$\mathbf{J}_{\text{sub}} = -D_{\text{sub}}(1 + 8\phi_{\text{sub}})\nabla\phi_{\text{sub}}. \tag{17}$$

Taking into account the ribosome-DNA and ribosome-polysome interactions yields the equation of motion:

$$\partial_t\phi_{\text{sub}} = D_{\text{sub}}\nabla\cdot\left[(1 + 8\phi_{\text{sub}})\nabla\phi_{\text{sub}} + \frac{B_{\text{DNA-sub}}}{V_{\text{DNA}}}\phi_{\text{sub}}\nabla\phi_{\text{DNA}} + \frac{B_{\text{poly-sub}}}{v_{\text{poly}}}\phi_{\text{sub}}\nabla\phi_{\text{poly}}\right]. \tag{18}$$

**IV. Computational domain and rescaling.** Our computational domain for the continuity equations is the interior of the *E. coli* cell, taken to be a cylinder of length $L$. The coordinate $x$ will denote the long dimension of the cell. We will assume a one-dimensional model in which the volume fractions only vary along $x$, maintaining a constant value in the other directions (i.e., the radial direction $r$ in cylindrical coordinates). However, the real cell is not homogeneous in the radial direction as the nucleoid typically does not occupy the whole cross-section of the cell. For simplicity, we choose a dimensionless length $x \in (-1,1)$, with $x = 0$ representing the center of the cell. This dimensionless coordinate can be converted to physical units by multiplying by $L/2$. We also define a dimensionless time $\tau$ by $\tau = 4D_{\text{poly}}t/L_0^2$, where $L_0 = 2\ \mu\text{m}$ is a reference cell length. This change enables us to remove the diffusion coefficient $D_{\text{poly}}$ from the equation of motion for $\phi_{\text{poly}}$. Our continuity equations (Eq. (6), (16), and (18)) then become

$$\partial_\tau\phi_{\text{DNA}} = \frac{L_0^2 D_{\text{DNA}} v_{\text{DNA}}}{L^2 D_{\text{poly}}}\,\partial_x\left[\frac{g\alpha(\alpha+1)V_{\text{DNA},0}^{\alpha}\phi_{\text{DNA}}^{\alpha}}{V_{\text{DNA}}^{\alpha+1}}\partial_x\phi_{\text{DNA}} + \frac{\phi_{\text{DNA}}}{V_{\text{DNA}}}\left(\frac{B_{\text{DNA-poly}}}{v_{\text{poly}}}\partial_x\phi_{\text{poly}} + \frac{B_{\text{DNA-sub}}}{v_{\text{sub}}}\partial_x\phi_{\text{sub}}\right)\right] \tag{19}$$

$$\partial_\tau \phi_{\text{poly}} = \frac{L_0^2}{L^2}\partial_x\left[\left(1+\frac{(\gamma+1)\phi_{\text{poly}}^{\gamma}}{\left(0.69\phi_{\text{poly}}^{\text{ov}}\right)^{\gamma}}\right)\partial_x\phi_{\text{poly}} + \frac{B_{\text{DNA-poly}}\phi_{\text{poly}}}{V_{\text{DNA}}}\partial_x\phi_{\text{DNA}} + \frac{B_{\text{poly-sub}}\phi_{\text{poly}}}{v_{\text{sub}}}\partial_x\phi_{\text{sub}}\right] \tag{20}$$

$$\partial_\tau \phi_{\text{sub}} = \frac{L_0^2 D_{\text{sub}}}{L^2 D_{\text{poly}}}\partial_x\left[(1+8\phi_{\text{sub}})\,\partial_x\phi_{\text{sub}} + \frac{B_{\text{DNA-sub}}\phi_{\text{sub}}}{V_{\text{DNA}}}\partial_x\phi_{\text{DNA}} + \frac{B_{\text{poly-sub}}\phi_{\text{sub}}}{v_{\text{poly}}}\partial_x\phi_{\text{poly}}\right]. \tag{21}$$

When the excluded volume interactions between DNA and polysomes are sufficiently strong, two minima in free energy are present, one corresponding to a DNA-rich “nucleoid” phase and the other to a DNA-poor cytosolic region (13).

**V. Interfacial free energies.** So far, we have only focused on the terms of free energy, describing the bulk of the phases. In a spatial model, however, there will be interfaces between the nucleoid and cytosol which would incur a free energy cost. In the phase separation regime, we also consider the free energy cost of interfaces between the nucleoid and cytosol phases. In an $N$-component mixture, the leading order interaction terms between species $i$ and species $j$ are proportional to $\nabla\phi_i \cdot \nabla\phi_j$. The general form of the interfacial energy to this order is thus

$$\beta\mathcal{F}_{\text{interface}} = \int \mathrm{d}x \sum_{i,j} \frac{\kappa_{ij}}{2v_i}\nabla\phi_i \cdot \nabla\,\phi_j, \tag{22}$$

where $\kappa_{ij}$ are interaction parameters (discussed in more detail in, e.g., (19)), and we sum over all species $i$ and $j$. The corresponding currents are

$$\mathrm{J}_{\text{CH},i} = -D_i v_i \phi_i \nabla \frac{\delta\beta\mathcal{F}_{\text{CH},i}}{\delta\phi_i} = D_i\phi_i \textstyle\sum_j \kappa_{ij}\nabla\nabla^2\phi_j\,. \tag{23}$$

The parameters $\kappa_{ij}$ (with units of area) depend on the details of the interfaces between species $i$ and $j$, which we expect to be narrow (molecular length scale). In our one-dimensional model, these interfacial terms contribute the following term to the equation of motion:

$$-\frac{L_0^2}{4D_{\text{poly}}}\nabla\cdot \mathrm{J}_{\text{CH},i} = -\frac{4L_0^2 D_i}{L^4 D_{\text{poly}}}\partial_x\left(\phi_i \sum_{\text{j}} \kappa_{ij}\,\partial_x^3\phi_j\right) \tag{24}$$

Note that sharp interfaces are only expected for the DNA and the polysomes, which are large polymeric species with distributions strongly influenced by crowding. On the other hand, the relatively small ribosomal subunits more freely diffuse between the cytosol and nucleoid regions, and we expect that spatial variations in $\phi_{\text{sub}}$ are relatively small. Thus, we will drop the interfacial energies for the ribosomal subunit.

We need to estimate the four $\kappa_{ij}$ that characterize the polysome and DNA volume fraction spatial variations. These parameters have units of area and set the interface width (see, e.g., (20)), with larger values increasing the cost of spatial variations and favoring larger interface widths. In practice, the stability of the coupled partial differential equations is sensitive to the values of $\kappa_{ij}$. The empirical values we set for this work are $\kappa_{\rm DD} \equiv \kappa_{\rm DNA-DNA} = 0.5\ \mu{\rm m}^2, \kappa_{\rm rr} \equiv \kappa_{\rm poly-poly} = 0.1\ \mu{\rm m}^2, \kappa_{\rm Dr} \equiv \kappa_{\rm DNA-poly} = 0.05\ \mu{\rm m}^2$ and $\kappa_{\rm rD} \equiv \kappa_{\rm poly-DNA} = 0.005\ \mu{\rm m}^2$. Since the DNA and polysomes are strongly segregated within the cell, relatively small values for the two cross terms are expected. On the other hand, the interface width between nucleoid (polysome-poor) and cytosol (polysome-rich) is observed to be thicker than the molecular size and we expect larger values of $\kappa_{\rm DD}$ and $\kappa_{\rm rr}$.

**VI. Auxiliary entropy.** Typical solutions of our model have the DNA volume fraction completely vanishing outside the nucleoid region, which can cause instabilities and lead to unphysical negative concentrations. To fix this issue, we follow the approach of Miangolarra *et al.* (16) and introduce an additional entropic term in the DNA free energy:

$$\beta \mathcal{F}_{\rm aux} = \frac{\alpha_1}{v_{\rm DNA}} \int {\rm d}x\ {\rm e}^{-\alpha_2 \phi_{\rm DNA}} \phi_{\rm DNA} \ln \phi_{\rm DNA}\,, \tag{25}$$

where $\alpha_1$and $\alpha_2$ are two constants. We choose $\alpha_2$ large enough so that the contribution of $\mathcal{F}_{aux}$ vanishes inside the nucleoid (with non-zero $\phi_{\rm DNA}$). On the other hand, the exponential is effectively equal to unity outside the nucleoid, as $\phi_{\rm DNA} \to 0$. Thus, $\mathcal{F}_{\rm aux}$ reduces to the standard entropic term. We choose an $\alpha_1$ small enough that this auxiliary term does not dominate in the cytosol. Accordingly, we set $\alpha_1 = 1$ and $\alpha_2 = 200$.

The auxiliary entropy adds the following term to our equation of motion:

$$\frac{\alpha_1 L_0^2 D_{\rm DNA}}{L^2 D_{\rm poly}} \partial_x \{ [\alpha_2^2 \phi_{\rm DNA}^2 \ln \phi_{\rm DNA} - 2(\ln \phi_{\rm DNA} + 1)\alpha_2 \phi_{\rm DNA} + 1] {\rm e}^{-\alpha_2 \phi_{\rm DNA}}\, \partial_x \phi_{\rm DNA} \}. \tag{26}$$

**Reactions and other dynamics.** Next, we will describe in detail the reactions involved in the model.

**I. Polysome assembly and disassembly.** Polysomes are produced when free ribosomes (the subunits) assemble on nascent mRNA that is being transcribed (co-transcriptional translation). Since we do not explicitly model mRNA, we assume that the transcription initiation rate is proportional to DNA density (volume fraction). The assembly rate of polysomes is then $\beta_{\rm p} \phi_{\rm DNA} \phi_{\rm sub}$, where $\beta_{\rm p}$ is a second-order reaction rate constant that will be estimated below.

Polysomes disassemble when the underlying mRNA backbone is degraded. We consider this to be a first-order reaction with a rate $\beta_d \phi_{\rm poly}$, where $\beta_d$ is the mRNA degradation rate. Based on Ref. (21), $\beta_{\rm d} = 0.003\ {\rm s}^{-1}$.

**II. Ribosome binding to existing polysomes and their unbinding.** The ribosomal subunits will assemble on the mRNA and unbind once they have translated the mRNA into a protein. When the subunits are assembled on the mRNA, they are removed from the free ribosomal subunit pool, which we assume occurs at rate $k_{\mathrm{on}}\phi_{\mathrm{poly}}\phi_{\mathrm{sub}}$, where $k_{\mathrm{on}}$ is the corresponding rate constant. Since the binding of the 30S subunit to mRNA is quickly followed by the binding of the 50S subunit, we consider binding to involve the whole 70S ribosome rather than sequential binding of individual subunits. Similarly, for the ribosome unbinding from mRNA, both subunits detach effectively simultaneously, and we consider the detachment of the whole ribosome. This will occur at a rate $k_{\mathrm{off}}\phi_{\mathrm{poly}}$. The average rate for a ribosome to complete protein translation is approximately $0.025\ \mathrm{s}^{-1}$, as reported in Ref. (22). In our model, we use the parameter $N_{\mathrm{ribo}}$ to describe the average number of ribosomes on the mRNA. We take $N_{\mathrm{ribo}} = 10$, as in (13,14). The rate at which any one of these ribosomes unbinds from the mRNA is thus approximated as $k_{\mathrm{off}} \approx (0.025\ \mathrm{s}^{-1})\frac{N_{\mathrm{ribo}}v_{\mathrm{sub}}}{v_{\mathrm{poly}}} \approx 0.0058\ \mathrm{s}^{-1}$.

The binding rate $k_{\mathrm{on}}$ is difficult to determine by comparisons to experiments since it is a bimolecular rate, determined by both the abundance of the polysomes and the subunits. We do know that if $k_{\mathrm{on}}$ is too large, then the pool of free ribosomes will be rapidly depleted. Also, when the polysome assembly is turned off ($\beta_{\mathrm{p}} = 0$) and the polysome volume fraction $\phi_{\mathrm{poly}}$ should approach zero and the subunit fraction $\phi_{\mathrm{sub}}$ should approach a constant value $\phi^*_{\mathrm{sub}} \approx 0.09$ corresponding to fully dissociated ribosomes. This steady-state value $\phi^*_{\mathrm{sub}}$ will be determined by the balance between the various reactions associated with the polysome at vanishing volume fraction. We can fix the binding rate $k_{\mathrm{on}}$ by balancing these reactions (ribosome binding, unbinding, and mRNA degradation): $k_{\mathrm{on}} \approx (\phi^*_{\mathrm{sub}})^{-1}\left[k_{\mathrm{off}} + N_{\mathrm{ribo}}\beta_{\mathrm{d}}\frac{\mathrm{v_{sub}}}{\mathrm{v_{poly}}}\right] \approx 0.073\ \mathrm{s}^{-1}$, where $\beta_{\mathrm{d}} = 0.003\ \mathrm{s}^{-1}$, as discussed above. As this is only a rough estimate for the rate $k_{\mathrm{on}}$, we explore a range of possible values for this parameter below.

**III. Estimation of polysome assembly rate, $\boldsymbol{\beta_p}$.** The parameter $\beta_p$, another bimolecular reaction rate constant, is similarly difficult to determine from the experiments. Again, we rely on a steady-state argument to make an estimate.

Ignoring diffusion, the ribosomal subunit volume fraction evolves as:

$$\frac{\mathrm{d}\phi_{\mathrm{sub}}}{\mathrm{d}t} = -k_{\mathrm{on}}\phi_{\mathrm{poly}}\phi_{\mathrm{sub}} + k_{\mathrm{off}}\phi_{\mathrm{poly}} + \frac{v_{\mathrm{sub}}N_{\mathrm{ribo}}}{v_{\mathrm{poly}}}\left[\beta_d\phi_{\mathrm{poly}} - \beta_{\mathrm{p}}\phi_{\mathrm{DNA}}\phi_{\mathrm{sub}}\right], \tag{27}$$

Here the factor $v_{\mathrm{sub}}N_{\mathrm{ribo}}/v_{\mathrm{poly}} \approx 0.23$ accounts for the difference in volume between the polysome and the individual ribosomes. Similarly, the polysome volume fraction obeys:

$$\frac{\mathrm{d}\phi_{\mathrm{poly}}}{\mathrm{dt}} = \beta_p\phi_{\mathrm{DNA}}\phi_{\mathrm{sub}} - \beta_d\phi_{\mathrm{poly}} + \frac{v_{\mathrm{poly}}}{v_{\mathrm{sub}}N_{\mathrm{ribo}}}\left(k_{\mathrm{on}}\phi_{\mathrm{poly}}\phi_{\mathrm{sub}} - k_{\mathrm{off}}\Phi_{\mathrm{poly}}\right). \tag{28}$$

Note that these equations ensure that the total amount of ribosomes in the cell, expressed as $V_{\mathrm{cell}}\left(\frac{\phi_{\mathrm{sub}}}{v_{\mathrm{sub}}} + N_{\mathrm{ribo}}\frac{\phi_{\mathrm{poly}}}{v_{\mathrm{poly}}}\right)$, remains constant.

Setting Eq. (28) to zero for the steady-state condition and solving for $\beta_p$ yields:

$$\beta_p = \frac{\phi^*_{\text{poly}}}{\phi^*_{\text{sub}}\phi^*_{\text{DNA}}}\left[\beta_d + \frac{v_{\text{poly}}(k_{\text{off}} - k_{\text{on}}\phi^*_{\text{sub}})}{v_{\text{sub}}N_{\text{ribo}}}\right], \tag{29}$$

where the asterisks indicate steady-state values. The ratio of the polysome to DNA volume fractions is approximated as

$$\frac{\phi^*_{\text{poly}}}{\phi^*_{\text{DNA}}} \approx \frac{\int dx\ \phi_{\text{poly}}(x)}{\int dx\ \phi_{\text{DNA}}(x)} = \frac{n_{\text{poly}}v_{\text{poly}}}{V_{\text{DNA}}} \approx 1.08, \tag{30}$$

where $n_{\text{poly}} = 600$ is the total number of polysomes. Note that $\phi_{\text{poly}}(x)$ and $\phi_{\text{DNA}}(x)$ are the volume fraction profiles extracted from the equilibrium model (without any reactions and obey the conservation conditions

$$\int \mathrm{d}x\ \phi_{\text{DNA}}(x) = \frac{2V_{\text{DNA}}}{V_{\text{cell}}} \approx 0.25, \tag{31}$$

$$\int \mathrm{d}x\ \phi_{\text{poly}}(x) = \frac{2n_{\text{poly}}v_{\text{poly}}}{V_{\text{cell}}} \approx 0.27, \tag{32}$$

The remaining parameter is the steady-state subunit volume fraction $\phi^*_{\text{sub}}$, which will depend on the number of free ribosomes. Substituting all parameters into Eq. (29), we estimate a range $\beta_p \approx 1 - 5\ \text{s}^{-1}$ for 1000 to 4000 free ribosomes in the cell. In our simulations, we choose a smaller value $\beta_p \approx 1.6\ s^{-1}$ but also explore a range of possible values (see below). Note that this estimate, like the one in the previous section for $k_{\text{on}}$, ignores the spatial distribution of the molecules and the full spatial model should be evolved in order to compare to experimental data.

**IV. Time-dependent cell growth.** Our model accommodates cell growth. We assume DNA and ribosome amounts increase proportionally to cell length during this growth. While the increase of ribosome numbers in proportion to cell size has been well documented, the amount of DNA in the cell does not follow this behavior. However, there are no overlapping replication cycles in the growth conditions we consider. Furthermore, we consider processes that start to occur at the time of the nucleoid splitting. The onset of the latter is when about 60% DNA is replicated. Around the split time, the increase in DNA amount is approximately proportional to the increase in cell size.

We treat the cell length $L$ as a time-dependent parameter $L \to L(t)$ in all of the equations of motion. In our rescaled equations, the interfacial terms (proportional to $\kappa_{ij}$) are proportional to $L^{-4}$ and the diffusive terms scale as $L^{-2}$. We update the cell length $L(t)$ according to $L(t) = L_i + \alpha_g t = L_i + \alpha_g L_0^2 \tau / (4D_{\text{poly}})$, where $\alpha_g$ is the instantaneous growth rate and $\tau$ is the dimensionless time. We can set $\lambda_g \equiv \alpha_g L_0^2/(4D_{\text{poly}})$ as the rate in terms of the $\tau$ time. A typical value, for $\alpha_g \approx 0.00020\ \mu\text{m/s}$, is $\lambda_g \approx 0.020\ \mu\text{m}$.

**V. Rifampicin treatment.** To model the effects from rifampicin (Rif) treatment, we set $\beta_p = 0$ to halt polysome assembly while maintaining a constant total number of ribosomes in the system. For simulations of Rif-treated cells, we use experimentally determined cell length vs time curve after start of treatment at

$t = 0$ : :

$$L_{\text{cell,fit}}(\text{t}) = L_0 + (L_\infty - L_0) \times \{1 - \exp[-(t - T_{\text{ab}})/\tau_{\text{rif}}]\} \quad (33)$$

The parameters $L_0$, $L_\infty$ and $\tau_{\text{rif}}$ are determined using fits to experimental data (SI Fig. S19D and SI Fig. S20D). The parameter $T_{\text{ab}}$ is the time when the antibiotic treatment starts. The best-fit values for $L_0$, $L_\infty$ and $\tau_{\text{rif}}$ values for different cell groups can be found in SI Table S4. We kept the DNA volume fraction constant during rifampicin treatment, which implies that the DNA amount grows proportional to cell length.

**Summary and model implementation.** In summary, the three equations of motion are

$$\partial_\tau \phi_{\text{DNA}} = \frac{L_0^2 D_{\text{DNA}}}{L^2 D_{\text{poly}}} \partial_x \left\{ \frac{1}{2} \gamma_{\text{DD}} \phi_{\text{DNA}}^\alpha \partial_x \phi_{\text{DNA}} + \alpha_1 [\alpha_2^2 \phi_{\text{DNA}}^2 \ln \phi_{\text{DNA}} - 2(\ln \phi_{\text{DNA}} + 1)\alpha_2 \phi_{\text{DNA}} + 1] \mathrm{e}^{-\alpha_2 \phi_{\text{DNA}}} \partial_x \phi_{\text{DNA}} + \gamma_{\text{Dr}} \phi_{\text{DNA}} \partial_x \phi_{\text{poly}} + \gamma_{\text{Ds}} \phi_{\text{DNA}} \partial_x \phi_{\text{sub}} \frac{1}{2} \right\} - \frac{4 L_0^2 D_{\text{DNA}} \phi_{\text{DNA}}}{L^4 D_{\text{poly}}} \partial_x^3 \left( \kappa_{DD} \phi_{\text{DNA}} + \kappa_{Dr} \phi_{\text{poly}} \right), \quad (34)$$

$$\partial_\tau \phi_{\text{poly}} = \frac{L_0^2}{L^2} \partial_x \left[ \left( 1 + \gamma_{rr} \phi_{\text{poly}}^\gamma \right) \partial_x \phi_{\text{poly}} + \gamma_{\text{rD}} \phi_{\text{poly}} \partial_x \phi_{\text{DNA}} + \gamma_{\text{rs}} \phi_{\text{poly}} \partial_x \phi_{\text{sub}} \right] - \frac{4 L_0^2 \phi_{\text{poly}}}{L^4} \partial_x^3 \left( \kappa_{rr} \phi_{\text{poly}} + \kappa_{rD} \phi_{\text{DNA}} \right) + \frac{L_0^2}{4 D_{\text{poly}}} \left[ \beta_p \phi_{\text{DNA}} \phi_{\text{sub}} - \beta_d \phi_{\text{poly}} + \frac{1}{\gamma_{pp}} \left( k_{\text{on}} \phi_{\text{poly}} \phi_{\text{sub}} - k_{\text{off}} \phi_{\text{poly}} \right) \right], \quad (35)$$

$$\partial_\tau \phi_{\text{sub}} = \frac{L_0^2 D_{\text{sub}}}{L^2 D_{\text{poly}}} \partial_x \left[ (1 + 8\phi_{\text{sub}}) \partial_x \phi_{\text{sub}} + \gamma_{sD} \phi_{\text{sub}} \partial_x \phi_{\text{DNA}} + \gamma_{sr} \phi_{\text{sub}} \partial_x \phi_{\text{poly}} \right] + \frac{L_0^2}{4 D_{\text{poly}}} \left[ \gamma_{pp} \beta_d \phi_{\text{poly}} - \gamma_{pp} \beta_p \phi_{\text{DNA}} \phi_{\text{sub}} - k_{\text{on}} \phi_{\text{poly}} \phi_{\text{sub}} + k_{\text{off}} \phi_{\text{poly}} \right], \quad (36)$$

where we introduced several dimensionless parameters, $\gamma_{i,j}$, to simplify the notation. These parameters are listed in Table S6. Note also that $L \equiv L(\tau) = L_i + \lambda_g \tau$ is time-dependent. All the parameters and their values are listed in Tables S5 and S6.

We employed FiPy (23), a finite volume partial differential equation (PDE) solver written in Python, to numerically solve the coupled equations Eq. (34-36). No flux (von Neumann) boundary conditions were applied at the system's two end points. The computational domain was discretized into $n = 200$ grid points, and time was discretized into steps of $\Delta\tau = 10^{-4}$. Simulations were initialized from the equilibrium state solution (without reaction terms) at the initial cell length $(L_i = 1.3\ \mu\text{m})$.

To compare the model to experimental data, which measures either the total density of 30S or 50S ribosomal subunits (most of which are incorporated into polysomes), we calculate the total ribosome density from the model as:

$$\phi_{\text{ribo,tot}}(x) = \phi_{\text{sub}}/v_{\text{sub}} + N_{\text{ribo}} \phi_{\text{poly}}/v_{\text{poly}}$$

**Comparing our model to other recent reaction-diffusion models**

Our modelling approach is like the one described in Miangolarra *et al.* (16) and Papagiannakis *et al*. (15). Like those models, we have a set of non-linear, coupled partial differential equations describing the time-evolution of concentrations inside the cell as a function of position $x$ along the long dimension of the cell. We will make a few comments on the differences between our approach and these previous ones. The major difference is that Miangolarra *et al.* allow for the full distribution of differently-sized polysomes in the system, which requires the use of an arbitrary number of concentration fields. We opt to simplify, like in Papagiannakis *et al*., and consider a single polysome concentration field representing the full distribution of differently-sized polysomes (including the size 0 representing a bare mRNA). Like the other two models, we also include a concentration field for the DNA and the freely-diffusing (unbound) ribosomes.

The other models use virial expansions to describe the excluded volume interactions between the various species. The molecules themselves are treated either as spheres (ribosomes, polysomes) or as cylinders (DNA supercoils/plectonemes) and the virial expansion is performed up to third order. On the other hand, we opt to use a variety of different interaction terms depending on the species. For example, for DNA, we use a power-law self-interaction (17) and the polysomes interact via a renormalized polymer-polymer excluded volume (18). We also allow for a variable radius of gyration for the polysome which is used in the excluded volume interactions with the DNA supercoils (treated as cylinders). Finally, for the ribosome-ribosome interactions, we use a Carnahan-Sterling expression which should work better at higher densities than the virial expansion.

Our implementation of the reactions is also slightly different because we assume that the polysome production depends on both DNA and free ribosome density. We also incorporate the cell growth dynamics directly into the equations of motion, much like in Papagiannakis *et al*. (15). Miangolarra *et al.*, on the other hand, evolve the reaction-diffusion dynamics for cells with various fixed lengths, which can lead to odd numbers of nucleoids inside the cell (see, e.g., Fig. S7 in Ref. (16)).

**The sensitivity of the model**

To assess the robustness of our model, we systematically varied three key parameters with the highest uncertainty: the ribosomal subunit binding rate ($k_{\text{on}}$), polysome production rate ($\beta_p$), and DNA self-interaction prefactor ($\gamma_{\text{DD}}$). (Note that we do not consider Cahn-Hilliard and the auxiliary entropy parameters critical one for the model.) Our default parameter values used in this work are $k_{\text{on}} = 0.073$ s$^{-1}$, $\beta_p = 1.6$ s$^{-1}$ and $\gamma_{\text{DD}} = 17.7$ as determined through parameter estimation described above. Unlike the simulations we present in the main text, we do not include time-dependent cell growth in these robustness tests. To isolate parameter effects from growth-dependent dynamics, we fixed the cell length at the reference cell length, i.e., $L = L_0 = 2$ μm. This length is near the threshold for nucleoid splitting (see Fig. 2 in the main text), so we expect to be near a transition between one and two nucleoids in parameter space. We solved the coupled PDEs Eqs. (34-36) numerically for various parameter combinations and classified steady-state solutions by nucleoid number using qualitative criteria. First, we determine whether the phase separation between DNA and crowders occurs by calculating the variance of $\phi_{\text{DNA}}(x)$ as:

$$\text{Var}[\phi_{\text{DNA}}] \equiv \frac{\sum_i |\phi_{\text{DNA}}(x_i) - \overline{\phi}_{\text{DNA}}|^2}{n},$$

where $n = 200$ is the number of grid points, $x_i$ is the $i$th grid point and $\overline{\phi}_{\text{DNA}}$ is the mean of $\phi_{\text{DNA}}(x)$. If the variance is below $2 \times 10^{-4}$, we consider the system is in "mixed" phase and not phase separated (e.g., SI Fig. S23B black triangle). Next, we apply extremum search to find minimum of $\phi_{\text{DNA}}$ (excluding the two minima on the boundary due to no flux boundary conditions). If the prominence of a minima, $A_{min}$, exceeds 5% of the neighboring maximum, it indicates nucleoid splitting. This criterion matches the one we used in experiments. In contrast, when the prominence of a minima is smaller than the threshold, the nucleoid is not considered as splitting (see SI Fig. S23B black diamond as an example).

We first test the sensitivity of the model with respect to $\beta_p$ and $\gamma_{\text{DD}}$ while keeping a fixed $k_{\text{on}} = 0.073$ s$^{-1}$. Parameter space exploration of $\beta_p$ and $\gamma_{\text{DD}}$ reveals distinct phase regions (SI Fig. S23A). The two-nucleoid phase (SI Fig. S23A, green disks) emerges for intermediate values ($1 \lesssim \beta_p \lesssim 20$ s$^{-1}$, $20 \lesssim \gamma_{\text{DD}} \lesssim 100$). If $\gamma_{\text{DD}}$ is too small, the repulsive DNA self-interaction will be too weak for polysomes within the nucleoid to form a nucleoid gap and induce nucleoid splitting. By contrast, larger $\gamma_{\text{DD}}$ leads to nucleoid splitting due to the nucleoid size expansion. However, if $\gamma_{\text{DD}}$ is too large, then the phase separation between DNA and crowders disappears as forming a DNA interface becomes too costly (SI Fig. S23A, yellow disks). On the other hand, the territory of the single-nucleoid phase expands as $\beta_p$ increases. The origin of this territory expansion is that when $\beta_p$ becomes large enough, the concentrated polysome production at the peak of $\phi_{\text{DNA}}$ will be diluted (recall the assembly rate of polysomes is $\propto \beta_p \phi_{\text{DNA}} \phi_{\text{sub}}$). As a result, the relative amount of polysomes produced at the middle of the nucleoid will be too small to split the nucleoid. Indeed, most of the polysome accumulation will occur at the poles (SI Fig. S23, diamond point).

The mixed phase territory also decreases as $\beta_p$ increases because an increasing production rate will deplete the pool of ribosomal subunits, converting them to polysomes. The polysomes drive the formation of the nucleoid phase due to their larger excluded volume. The stronger interaction between DNA and polysomes enables the phase separation to occur at lower $\gamma_{\text{DD}}$ values although the total concentration of ribosomes is conserved in the model. It is worth noting that our default parameters ($\beta_p = 1.6$ s$^{-1}$, $\gamma_{\text{DD}} = 17.7$) lie at the boundary between two nucleoids and one nucleoid phases (SI Fig. S23A, magenta star). This is because the chosen cell length, $L = 2$ µm, corresponds to the cell length where the nucleoid splitting occurs in the model with the default parameter set. Our analysis here suggests that, in the regime of this default parameter set, the model is more sensitive to the value of $\gamma_{\text{DD}}$ than $\beta_p$ since the phase transition occurs only on the vertical direction (SI Fig. S23A).

We next individually examine how the values of $k_{\text{on}}$ affect the steady-state solutions while keep $\beta_p = 1.6$ s$^{-1}$ and $\gamma_{\text{DD}} = 17.7$ fixed. The simulation reveals that increased ribosomal subunit binding rates suppress nucleoid splitting (SI Fig. S24A). This transition is similar to what has been observed in the high $\beta_p$ regime. The high subunit binding rate will dilute the fraction of polysomes being produced through the transcriptional polysome assembly at the nucleoid center. As a result, the amount of polysomes in the DNA-rich region is not enough to result in the nucleoid splitting and most of the polysomes accumulate at pole regions (SI Fig. S24B, two subplots on the right). In this case, the result from the default $k_{\text{on}} = 0.073$ s$^{-1}$ is not on the boundary between the two different phases, which implies the model is not sensitive to the value of $k_{\text{on}}$ at the regime around the default parameter set.

Interestingly, the model shows that nucleoid splitting persists even when $k_{\text{on}} = 0$ (SI Fig. S24A). This implies that merely polysome assembly at rate $\beta_{\text{p}}\phi_{\text{DNA}}\phi_{\text{sub}}$ can lead to nucleoid splitting. This occurs because our model assumes uniform polysome length ($N_{\text{ribo}} = 10$), regardless of the values of $k_{\text{on}}$ and $k_{\text{off}}$. Therefore, the effect of $k_{\text{on}}$ is less apparent in the model. However, $k_{\text{on}}$ and $k_{\text{off}}$ should govern the distribution of polysome length, which further affects the strength of interaction between DNA and polysomes. Our simplified model presented in this work can be extended to take the polysome length distribution into account by increasing the number of concentration fields and differential equations for polysomes in different lengths, as the approach used in Miangolarra *et al.* (16). In that scenario, we expect to see mixed phase when $k_{\text{on}} = 0$, as a result of all polysomes eventually lose most of the ribosomal subunits and the excluded volume interaction between DNA and polysomes decreases significantly.

**The concentric shell model**

The coupled DNA and ribosome dynamics model predicts about ten times higher ribosome fraction at the midcell (50%) than observed in the experiments (5%). We hypothesize that this discrepancy could arise from the 1D nature of the model that cannot account for a cylindrical shell of ribosomes surrounding the nucleoid. This shell has been observed in single molecule tracking experiments (24). To account for this shell, we add it *ad hoc* to the model. The shell adds extra ribosomes to those that were already present in the solution for the total ribosome volume fraction profile $\phi_{ribo,tot}(x)$ in the 1D model. We furthermore assume that the DNA density within this region is zero.

We assume the 3D cell to be a cylinder with hemispherical end caps (SI Fig. S12A). We denote the radius of the cylindrical portion by $R_{cell}^0$ and the radius in the hemispherical caps as $R_{\text{cell}}(x) = \sqrt{{R_{\text{cell}}^0}^2 - x^2}$, where $x$ is the distance into the hemispherical region along the long axis of the cell (SI Fig. S12A). This cell is then divided into two coaxial regions. The inner region (green shading in SI Fig. S12A) represents the existing solution $\phi_{ribo,tot}(x)$ from the coupled ribosome and DNA dynamics model. The outer region represents the extra shell with a constant 3D ribosome density. The interface between the two regions is defined by the radius $R_{1D}(x)$. This radius was taken to be proportional to the cell radius, that is $R_{\text{1D}}(x) = R_{\text{cell}}(x)\, R_{\text{1D}}^0 / R_{\text{cell}}^0$. Here $R_{\text{1D}}^0$ corresponds the shell's inner radius along the constant cylindrical body of the shell. This is the sole unknown parameter of the model and is determined by matching the concentric shell model to the experiment.

In the outer shell region, $\phi_{\text{3D,ribo,tot}}(x, y, z)$ was assumed to be constant, and its value matched this of the inner shell at its extreme points along the long axis of the cell $(x = 0, L_{\text{cell}})$. So $\phi_{\text{3D,ribo,tot}}(x, y, z) = \phi_{\text{ribo,tot}}(0)$ was postulated to be the 3D volume fraction in the outer added shell.

Under these assumptions a locally linear transformation between the original solution $\phi_{ribo,tot}(x)$ and the resulting 1D volume fraction profile from the concentric shell model $\tilde{\phi}_{\text{ribo,tot}}(x)$ holds:

$$\tilde{\phi}_{\text{ribo,tot}}(x) = \left(\phi_{\text{ribo,tot}}(x) + \phi_0\right)\left(\frac{R_{\text{cell}}(x)^2}{{R_{\text{cell}}^0}^2}\right)$$

Here, $\phi_0 \equiv \phi_{\text{ribo,tot}}(0)\left(\frac{{R_{\text{cell}}^0}^2}{{R_{\text{1D}}^0}^2} - 1\right)$. We found a good match to the data when $R_{\text{1D}}^0 \approx 0.5 R_{\text{cell}}^0$. This match by no means validates the concentric shell model. However, it shows that the coupled ribosome and DNA dynamics model, despite its large discrepancy in numerical values to experiment, is able to capture conceptually some of the experimental findings.

### Supplementary Figures

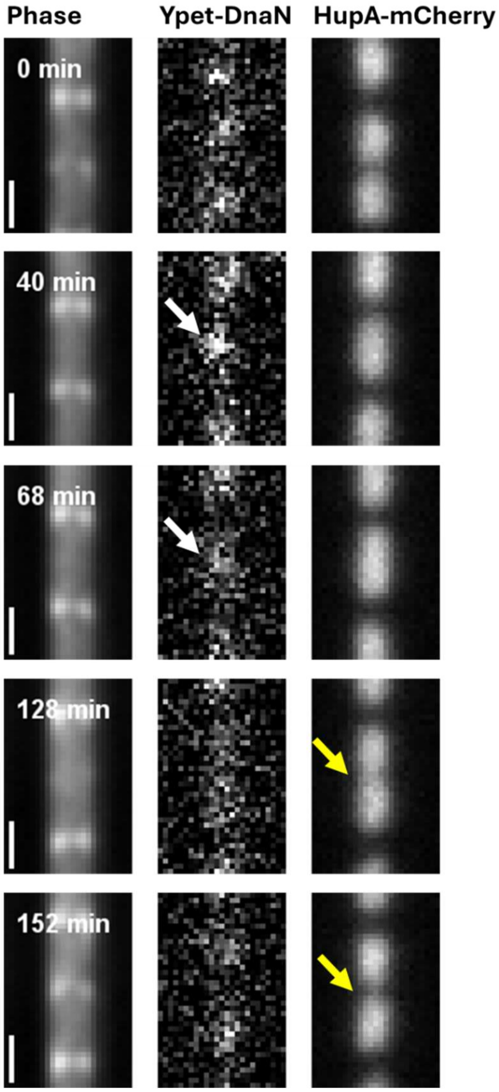


**SI Fig. S1.** Images of a representative cell with the replisome (Ypet-DnaN) and nucleoid (HupA-mCherry) labels through one cell cycle in M9 glycerol + trace elements medium (strain STK32). White arrows point Ypet-DnaN foci that form during the replication. Yellow arrows point to the DNA density minimum at the center of the nucleoid. Scale bar is 1 μm.

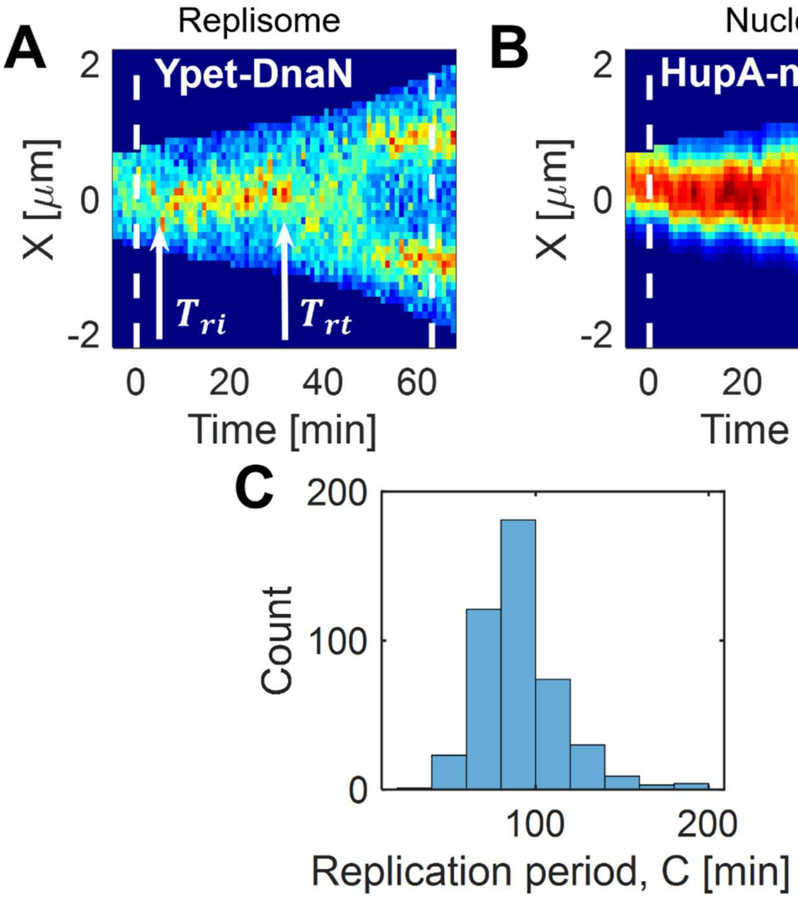


**SI Fig. S2**. Determining replication timing for strain STK32 under slow growth conditions in M9 glycerol medium. (A) A kymograph of Ypet-DnaN signal from an individual cell. The arrows point to the initiation ($T_{ri}$) and termination ($T_{rt}$) of DNA replication. The vertical dashed lines mark cell birth and division. (B) A kymograph of HupA-mCherry signal of the same cell. (C) Histogram showing the distribution of replication periods, $C = T_{rt} - T_{ri}$ for a cell population in slow growth conditions. The mean of the distribution is 91 min and standard deviation 24 min. $N = 44$

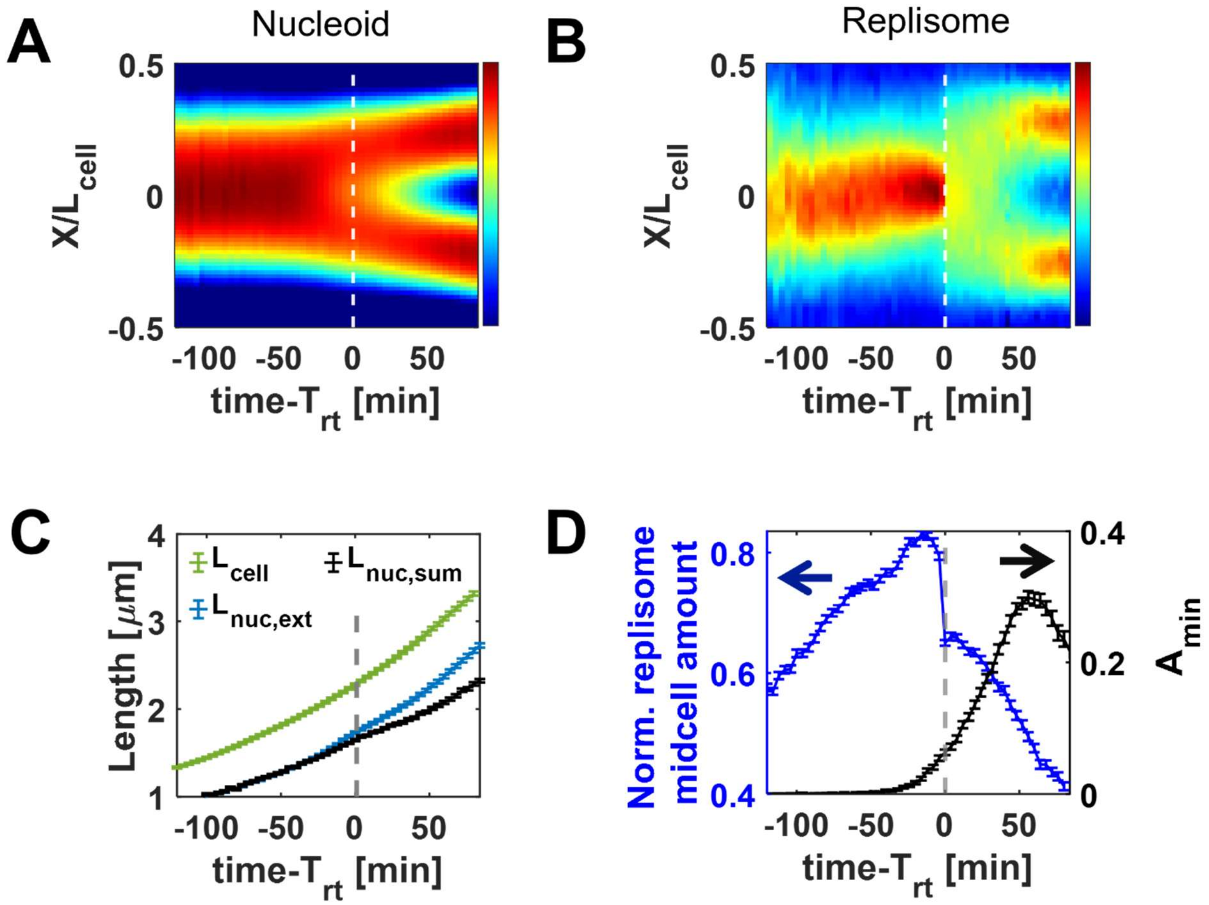


**SI Fig. S3.** Characterizing DNA replication and segregation relative to replication termination ($T_{rt}$) time in the STK32 strain during slow growth conditions in M9 glycerol medium. (A) Population averaged kymograph of DNA density distribution (HupA-mCherry) along the long axis of the cell as a function of time relative to replication termination (*N*=438). (B) The same for the density distribution of the replisome label (Ypet-DnaN). (C) Changes in cell length ($L_{\mathrm{cell}}$), nucleoid extent ($L_{\mathrm{nuc,ext}}$), and the sum of nucleoid lengths ($L_{\mathrm{nuc,sum}}$) for the same cell population aligned to replication termination. (D) Plot showing normalized replisome label Ypet-DnaN midcell amount (blue) and normalized depth of the local DNA density minimum ($A_{\mathrm{min}}$, black) aligned to replication termination. Error bars in panels C and D are s.e.m. $N = 386$.

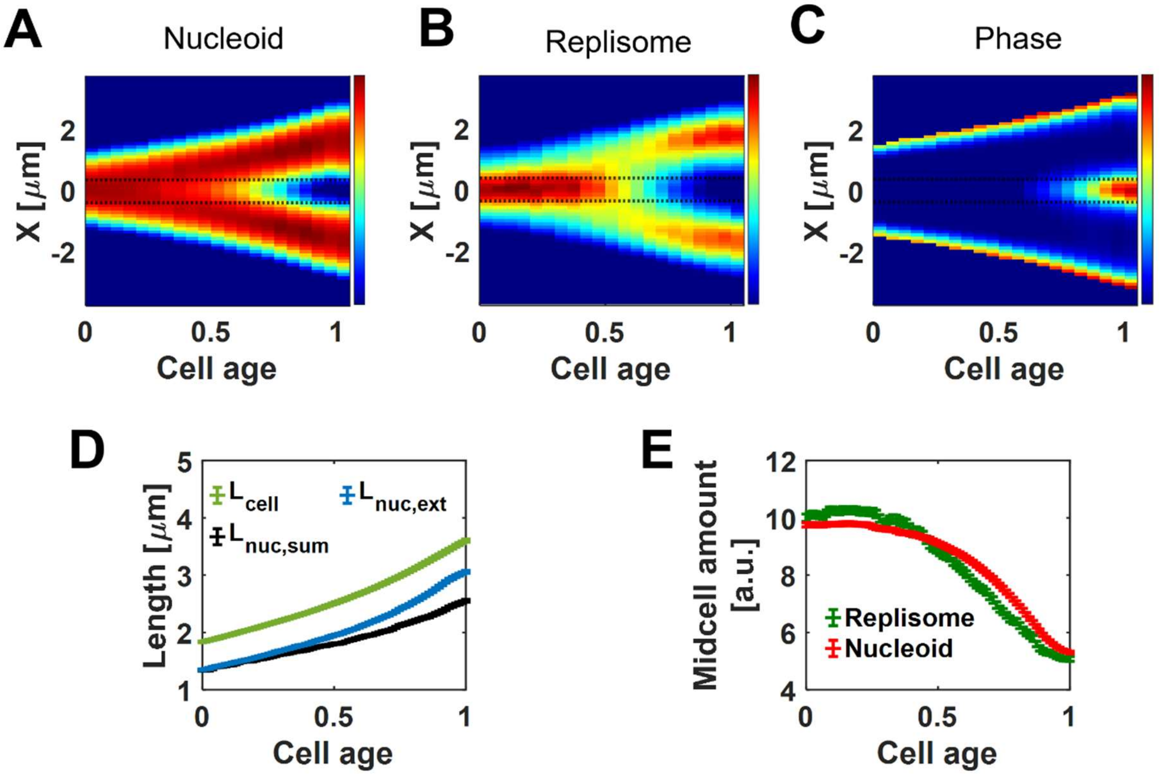


**SI Fig. S4**. Characterizing DNA replication and segregation during the cell cycle (from cell birth=0 to division=1) for STK32 strain under slow growth conditions in M9 glycerol medium. Population-averaged kymographs of (A) the density distribution of nucleoid labeled by HupA-mCherry, (B) replisome labeled by Ypet-DnaN, and (C) phase contrast signal along the long axis of the cell as a function of cell cycle time. Red corresponds to high- and blue to low-intensity values. The dashed horizontal lines indicate the midcell region. (D) Changes in cell length $(L_{\text{cell}})$, nucleoid extent $(L_{\text{nuc,ext}})$, and the sum of nucleoid lengths $(L_{\text{nuc,sum}})$ for the same cell population from cell birth to cell division. (E) Midcell amount of DNA (HupA-mCherry, red) and replisome (Ypet-DnaN, green) as a function of cell age. The midcell is defined as 0.75 μm (7 pixels) wide band. Error bars in panels (D) and (E) are s.e.m. $N = 274$

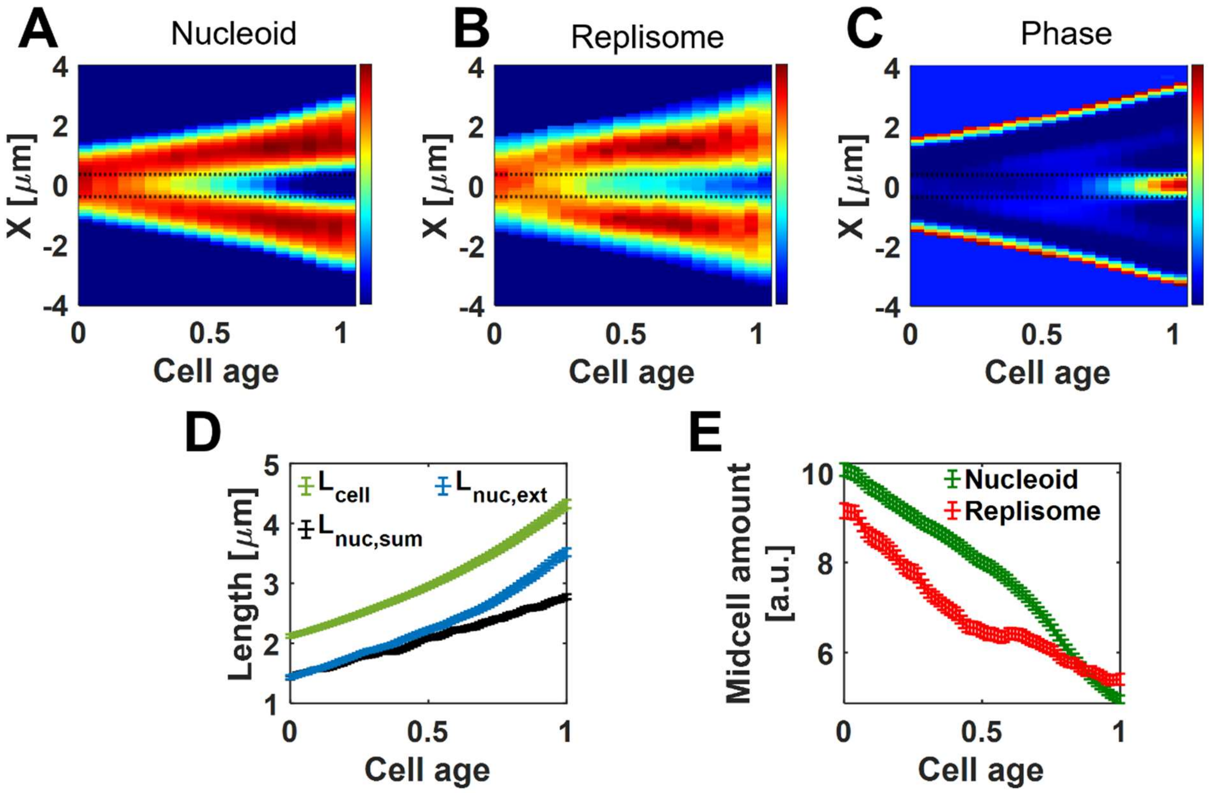


**SI Fig. S5.** Characterizing DNA replication and segregation during the cell cycle (from cell birth=0 to division=1) for STK30 strain during moderately fast growth conditions in M9 glucose + CAS medium. Population-averaged kymographs of (A) the density distribution of nucleoid labeled by HupA-mYpet, (B) replisome labeled by mCherry-DnaN, and (C) phase contrast signal along the long axis of the cell as a function of cell cycle time. Red corresponds to high- and blue to low-intensity values. The dashed horizontal lines indicate the midcell region. (D) Changes in cell length ($L_{\mathrm{cell}}$), nucleoid extent ($L_{\mathrm{nuc,ext}}$), and the sum of nucleoid lengths ($L_{\mathrm{nuc,sum}}$) for the same cell population from cell birth to cell division. (E) Midcell amount of DNA (HupA-mYpet, green) and replisome (mCherry-DnaN, red) as a function of cell age. The midcell is defined as 0.75 μm (7 pixels) wide band. Error bars in panels (D) and (E) are s.e.m. $N = 161$.

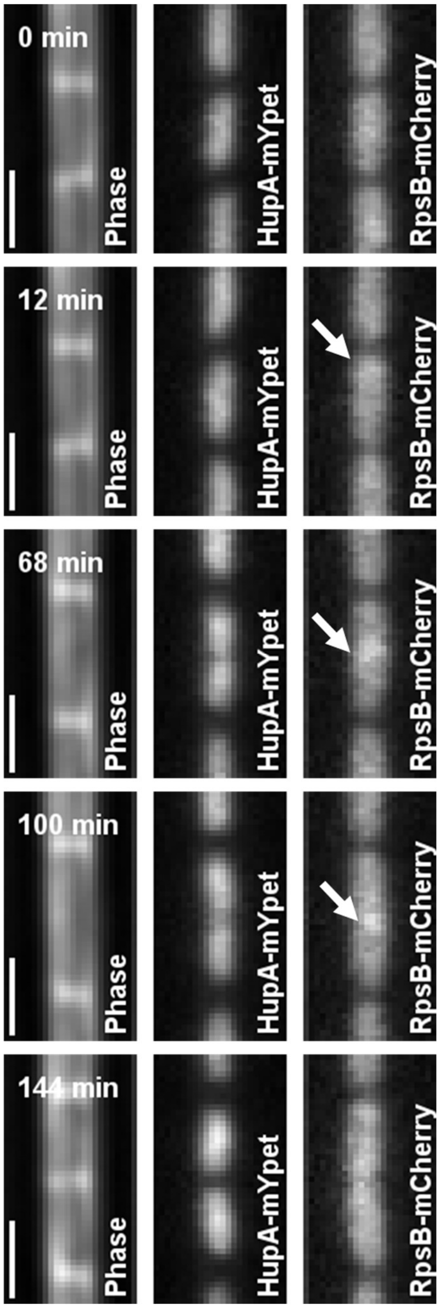


**SI Fig. S6.** Images of a representative cell with nucleoid (HupA-mYpet) and ribosome (RpsB-mCherry) labels through one cell cycle in M9 glycerol + trace elements medium (strain CA4). White arrows point to excess ribosome accumulations. Scale bar is 1 μm.

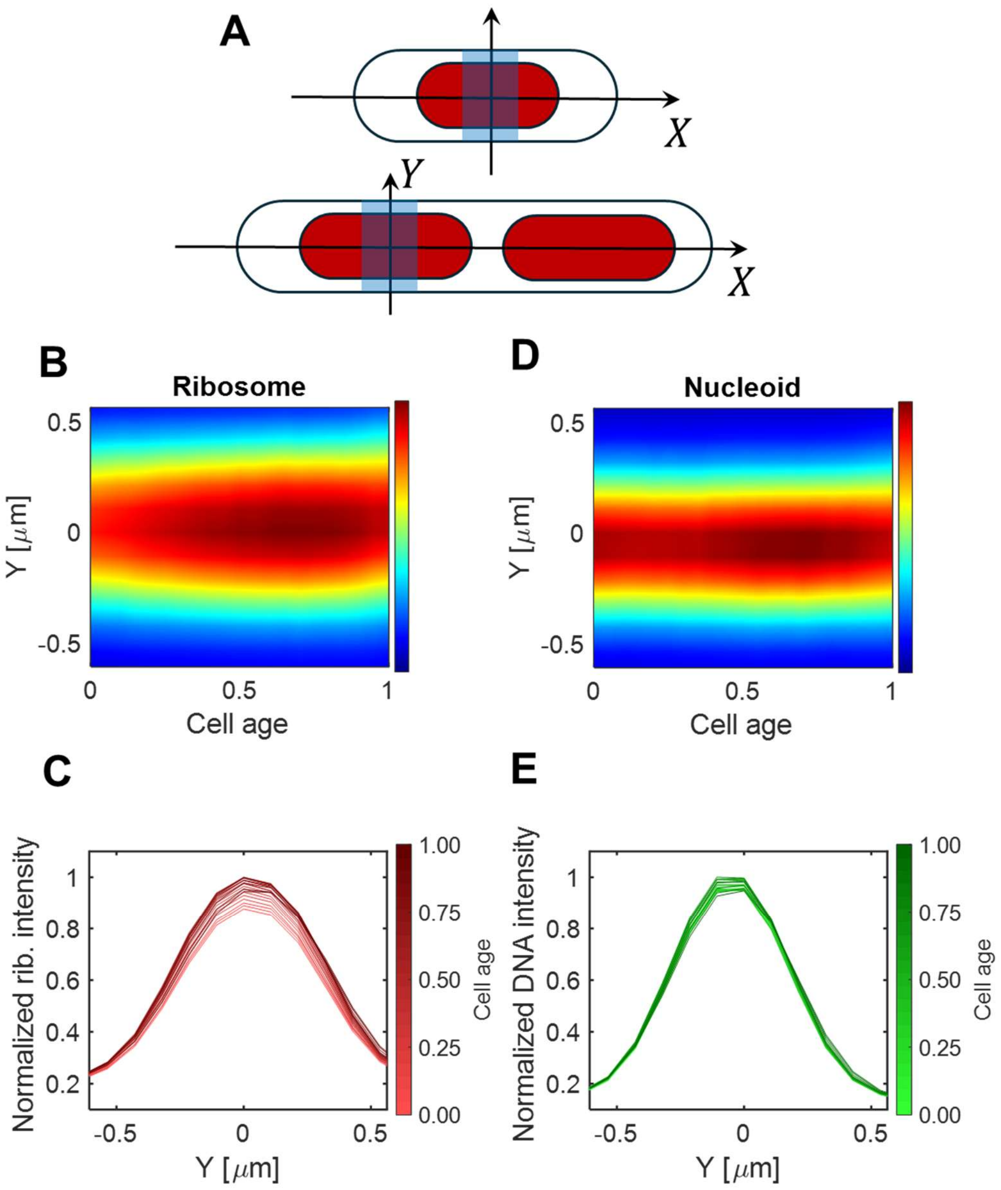


**SI Fig. S7.** The population-averaged nucleoid and ribosome density distributions along the short axes of the cell throughout the cell cycle. The measurements use CA4 strain in slow-growth conditions in M9 glycerol medium supplemented with trace elements. (A) Schematics showing calculation of the density distributions: The densities at individual cells are averaged over about 0.75 $\mu m$ wide band centered around the nucleoid centers (rather than cell centers). (B) Ribosomal density distribution along the short axes of the cell as a function of normalized cell cycle time. The distribution is based on the RpsB-mCherry signal. (C) Individual vertical slices form the kymograph in panel B at different cell cycle times. (D) DNA density distribution along the short axes of the cell as a function of normalized cell cycle time. The distribution is based on the HupA-mYpet signal. (E) Individual vertical slices form the kymograph in panel D at different cell cycle times. *N*=486.

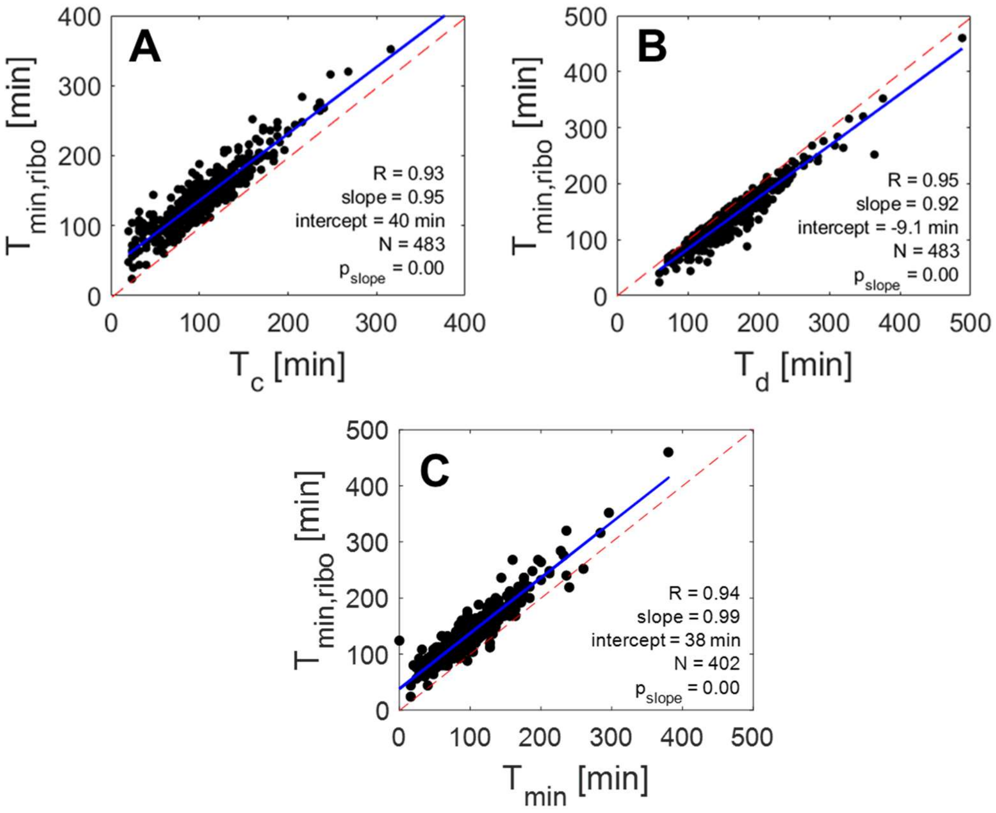


**SI Fig. S8.** Timing for the onset (appearance) of the local ribosome density minimum, $T_{min,ribo}$, as a function of other cell cycle events. $T_{min,ribo}$ is determined analogously to $T_{min}$, the timing for the onset of the DNA density minimum. For further details on how this quantity is determined, see SI Text, *Determining* $A_{min}$ *and* $T_{min}$ *from DNA distributions*. (A) $T_{min,ribo}$ as a function of the onset of constriction, $T_c$. The onset of constriction is determined from the phase contrast signal following the procedure described in (25). (B) $T_{min,ribo}$ as a function of cell doubling time (cell cycle time), $T_d$. (C) $T_{min,ribo}$ as a function of timing for the onset (appearance) of the local DNA density minimum, $T_{min}$. The blue solid lines in every panel show linear fits ($y = ax + b$) to the data. The Pearson correlation coefficient ($R$), the slope ($a$) and intercept ($b$) from each fit, number of cells plotted ($N$) and the probability that slope of the fit is zero ($p$) are indicated. The dashed red lines indicate the unit diagonal ($y = x$).

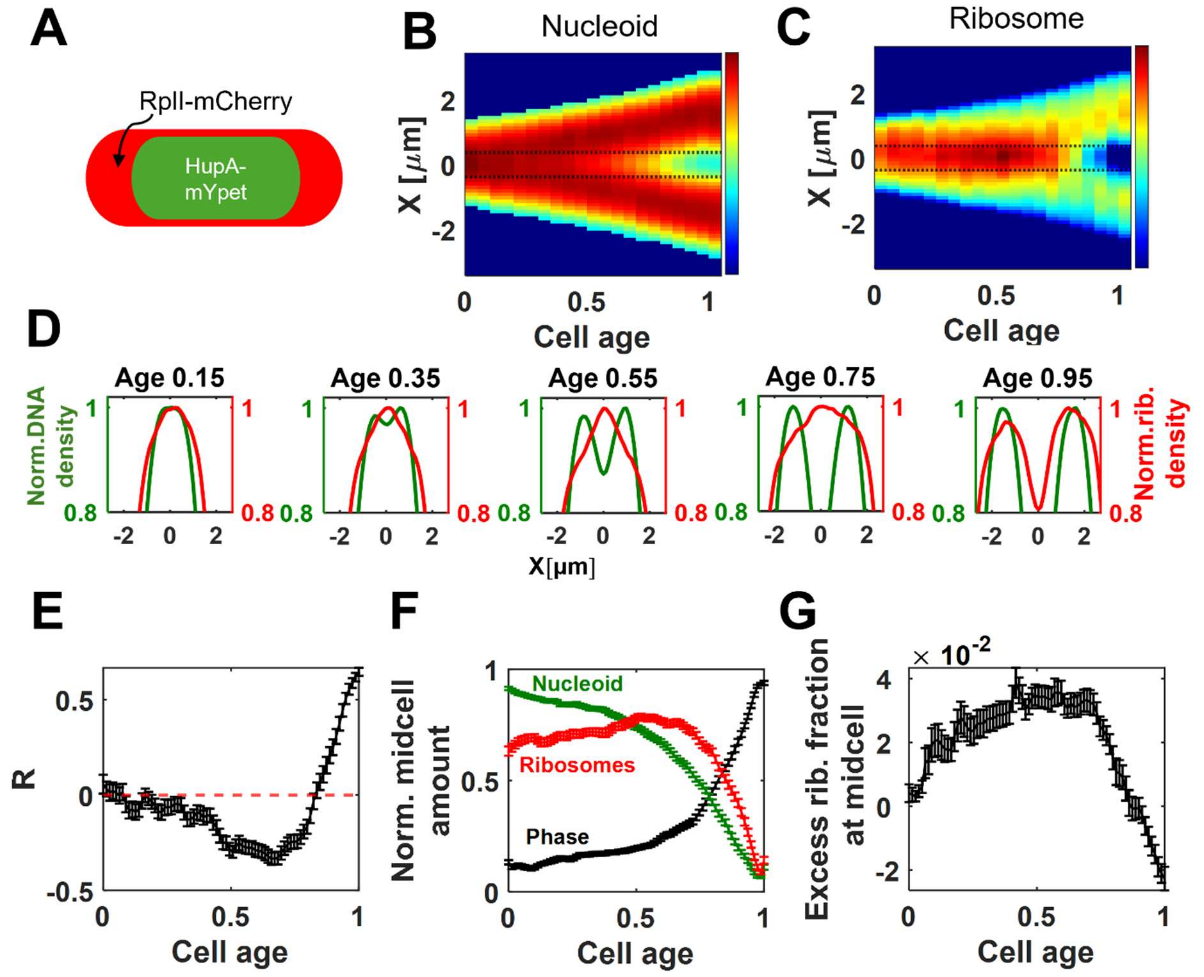


**SI Fig. S9**. Cell-cycle dependent changes in DNA and ribosome density distributions in the CA7 strain in slow growth conditions in M9 glycerol medium. CA7 carries mCherry fusion to the 50S ribosomal protein RplI (RplI-mCherry) and HupA-mYpet labeling. (A) A schematic illustrating this labeling. (B) Kymograph of the population-averaged DNA density (HupA-mYpet) distribution along the long axis of the cell as a function of cell cycle time. $N = 3140$. (C) Kymograph of the average ribosome density (RplI-mCherry) distribution for the same cell population. For both kymographs, zero marks cell birth, and one division. Red corresponds to high- and blue to low-intensity values. The dashed horizontal lines indicate the midcell region. (D) DNA (green) and ribosome (red) density distributions along the long axis of the cell at different stages of cell cycle. (E) Pearson correlation coefficient, $R$, between nucleoid and ribosomes densities as a function of cell age. The coefficient is averaged over the cell population at each cell age point. (F) The integrated intensity at the 0.75 $\mu$m wide band around the cell middle for the nucleoid (green), ribosome (red) and phase contrast signals (black). The curves from individual cells are normalized by their peak values and then averaged over the cell population. (G) The excess fraction of ribosomes at this midcell band as a function of cell cycle time. The curve is derived from a fitting of the ribosome signal (see Methods). All error bars are s.e.m. $N = 106$ for panels A-G.

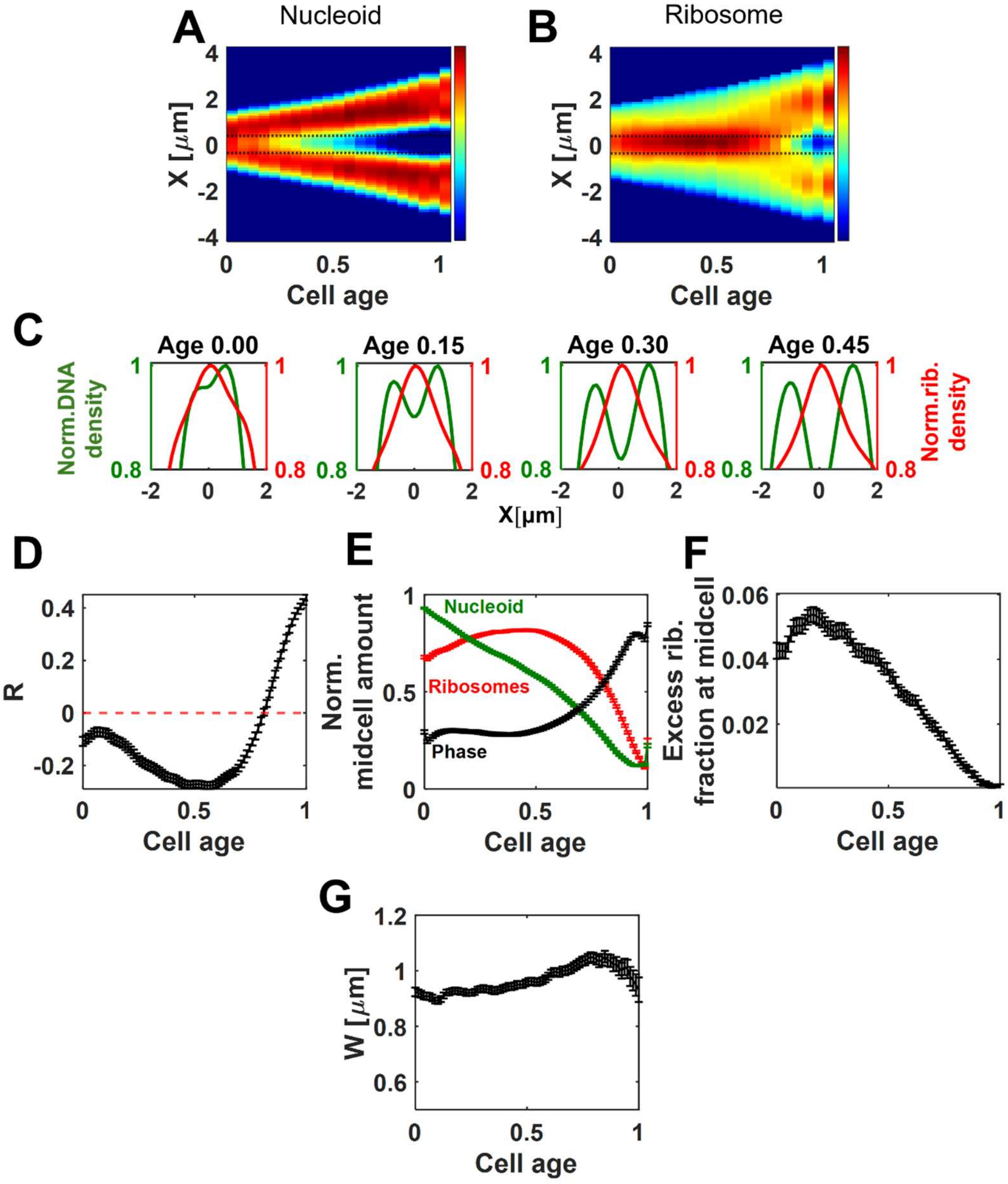


**SI Fig. S10**. Cell-cycle dependent changes of DNA and ribosome density distributions in CA4 strain in moderately fast growth conditions in M9 glucose + CAS medium. (A) The population average DNA density distribution along the long axis of the cell as a function of cell cycle time. HupA-mYpet labeling is used to infer the density distribution of DNA. (B) The corresponding average ribosome density distribution kymograph for the same cell population. RpsB-mCherry label is used to image ribosomes. (C) DNA (green) and ribosome (red) density distributions along the long axis of the cell at different stages of cell cycle. (D) Pearson correlation coefficient, $R$, between nucleoid and ribosome densities as a function of cell age. The coefficient is averaged over the cell population at each cell age point. (E) The integrated intensity at the 0.75 $\mu$m wide band around the cell middle for the nucleoid (green), ribosome (red) and phase contrast signals (black). The curves from individual cells are normalized by their peak values and then averaged over the cell population. (F) The excess fraction of ribosomes at this midcell band as a function of cell cycle time. (G) The width of the midcell ribosome accumulation, $W$, as a function of cell age. The excess fraction of ribosomes in panel F and the $W$ in panel G is determined from the fitting of the ribosome signal by generalized Gaussian curves (see Methods for details). Error bars in panels D-G are s.e.m. *N*=635

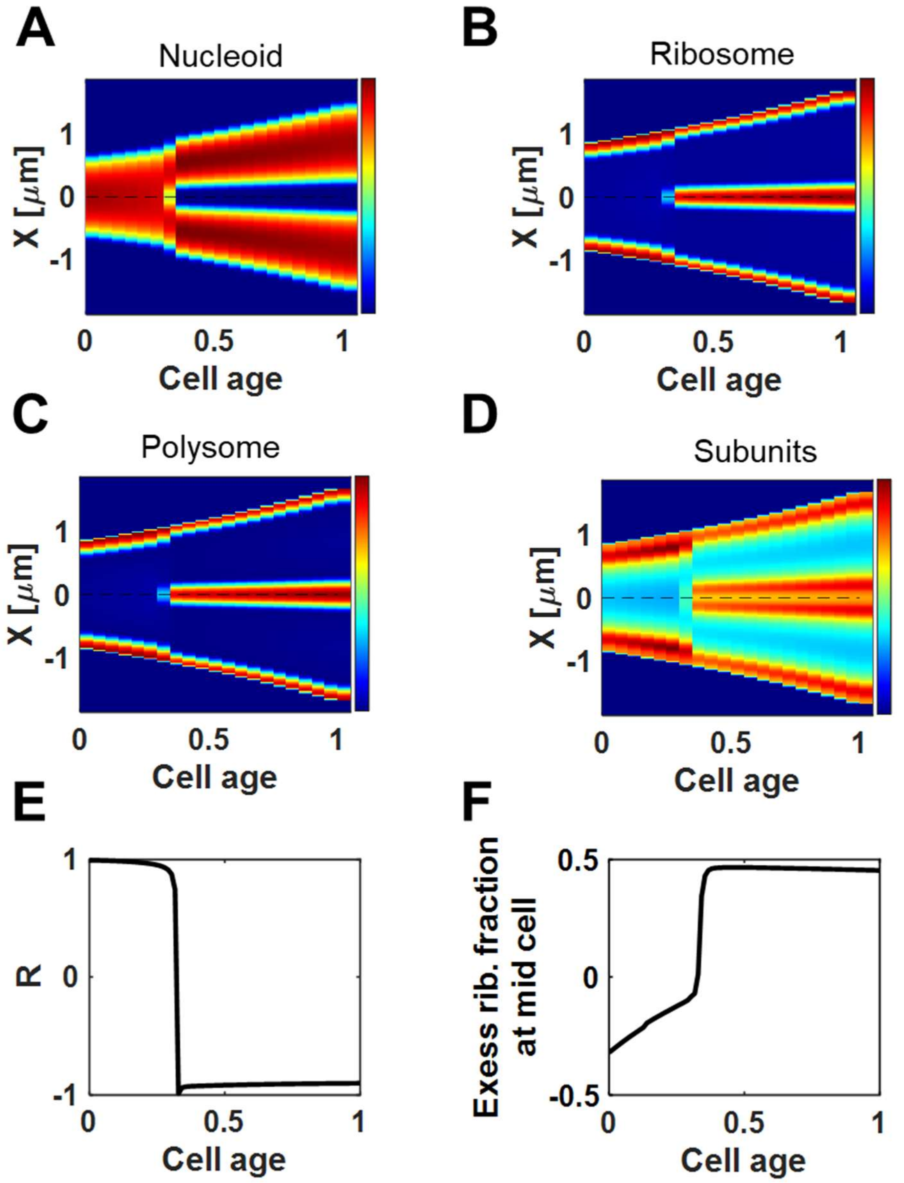


**SI Fig. S11**. Results from the coupled DNA and ribosome dynamics model. (A) The kymograph of DNA density. (B) Kymograph of the total ribosome density from the model. The total ribosome density includes ribosomes in both polysomes and free ribosomes. The total ribosome density is plotted in all the main text modeling kymographs pertinent of ribosomes. (C) Kymograph of the polysome density. The same kymograph also corresponds to the ribosome density in polysomes because each polysome is assumed to have the same number of ribosomes ($N_{ribo} = 10$). (D) Kymograph of subunit density. (E) The Pearson correlation coefficient, $R$, between DNA and total ribosome densities as a function of cell age. (F) Evolution of the excess total ribosome fraction at midcell from cell birth to cell division from the model.

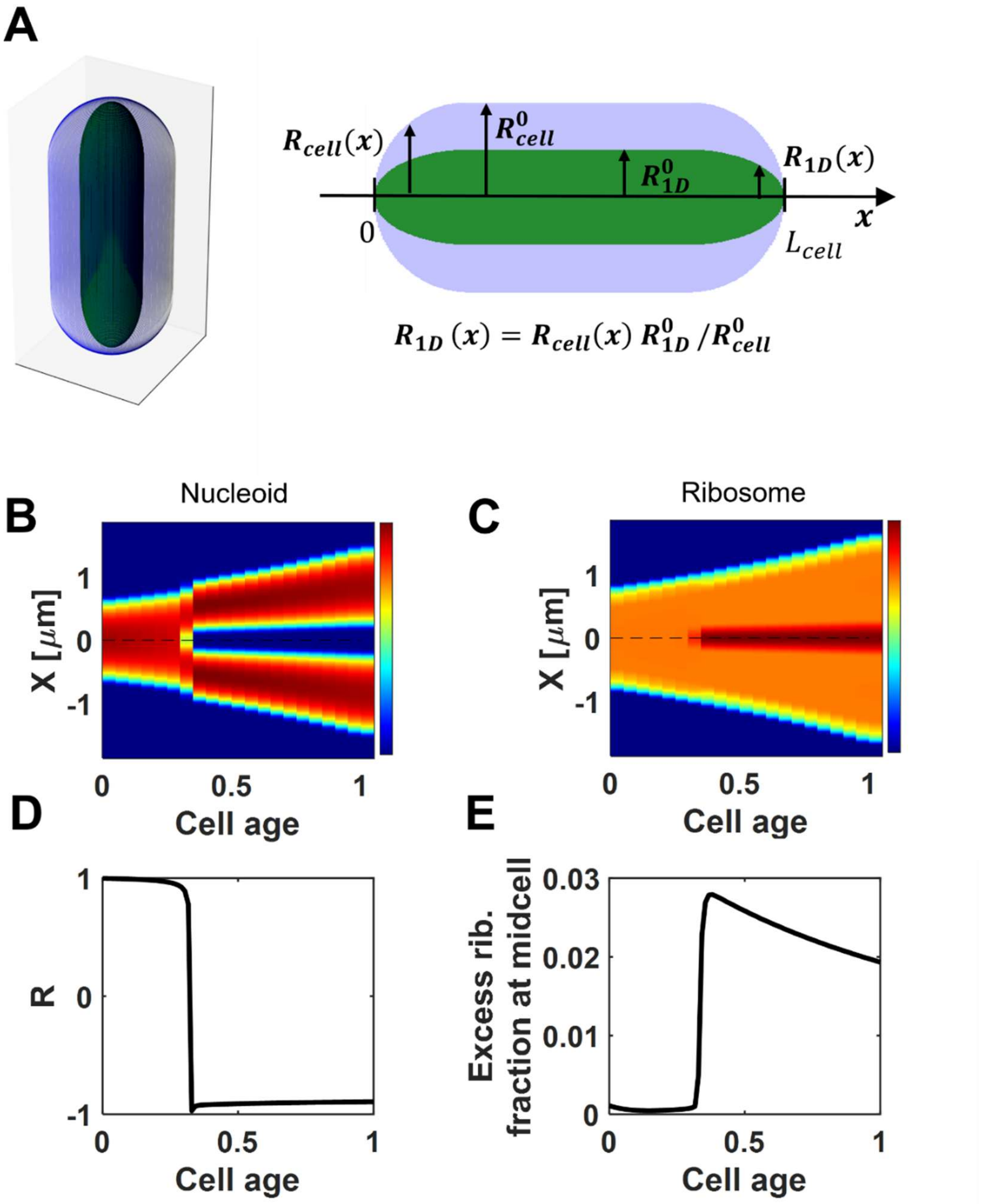


**SI Fig. S12**. The coupled DNA ribosome dynamics model with an extra *ad hoc* concentric shell of ribosomes. (A) Schematic showing the extra concentric shell (light blue) that is added to the solution from the coupled DNA and ribosome dynamics model (green). Note that the inner green region does not represent the nucleoid region but marks a region corresponding to the 1D model. Left: The 3D cylindrical cell with hemispherical end caps consists of two concentric shells. Right: Cross-section of this cell along the long axis of cylinder. The ratio of the outer radii of the two regions is constant throughout the cell length. The concentration of ribosomes is constant in the outer shell and equals that at the extreme points of the inner shell that are at $x = 0$ and $x = L_{\mathrm{cell}}$ (for more details see Methods, The Concentric Shell Model). (B) The kymographs of DNA and (C) ribosome densities from this model. (D) Pearson correlation coefficient, $R$, between DNA and ribosome densities as a function of cell age. (E) The excess ribosome fraction at midcell from cell birth to cell division from this model.

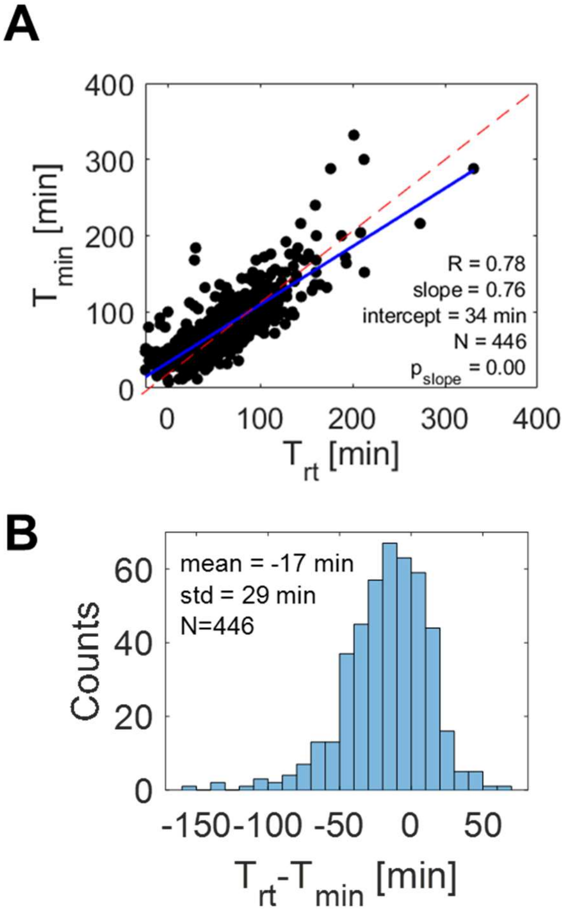


**SI Fig. S13.** The timing for the onset of the local DNA density minimum, $T_{min}$, relative to the timing of replication termination, $T_{rt}$. $T_{min}$ is determined when $A_{min}$ (see Fig. 1E) reaches the threshold value of 0.05. Note that the local minimum in DNA density appears transiently before this threshold value is reached. $T_{min}$ is thus a coarse-grained value of the appearance of the minimum. In a previous work (26), the value appeared earlier in the replication cycle because the DNA density curves were deconvolved. No deconvolution was used here. (A) $T_{min}$ vs. $T_{rt}$ in slow growth conditions in M9 glycerol supplemented with trace elements. Strain STK32 carries HupA-mCherry and Ypet-DnaN labels. The blue curve is the linear fit to the data, with the best fit parameters shown in the inset of the panel. The Pearson correlation coefficient $(\mathrm{R})$, the slope $(\mathrm{a})$ and intercept $(\mathrm{b})$ from each fit, the number of cells analyzed $(\mathrm{N})$, and the probability that the slope of the fit is zero $(p_{slope})$ are indicated. The dashed red lines indicate unit diagonal $(\mathrm{y}=\mathrm{x})$. (B) $T_{rt}-T_{min}$ histogram based on the same data.

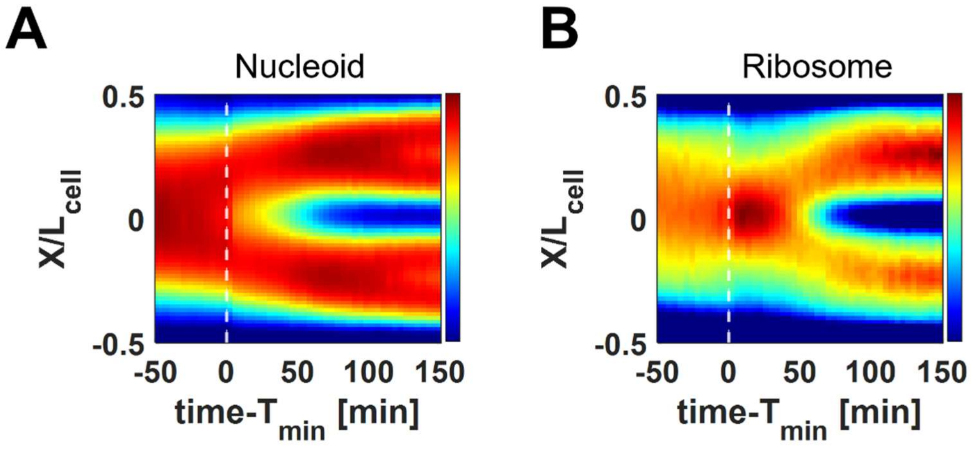


**SI Fig. S14**. Data from untreated CA4 cells in M9 glycerol medium aligned relative to the time when the local density minimum in the nucleoid distribution appears, $T_{\mathrm{min}}$. (A) Population-averaged kymograph of DNA density distribution (HupA-mYpet) along the long axis of the cell. (B) Population averaged kymograph of ribosome density distribution (RpsB-mCherry). *N*=378.

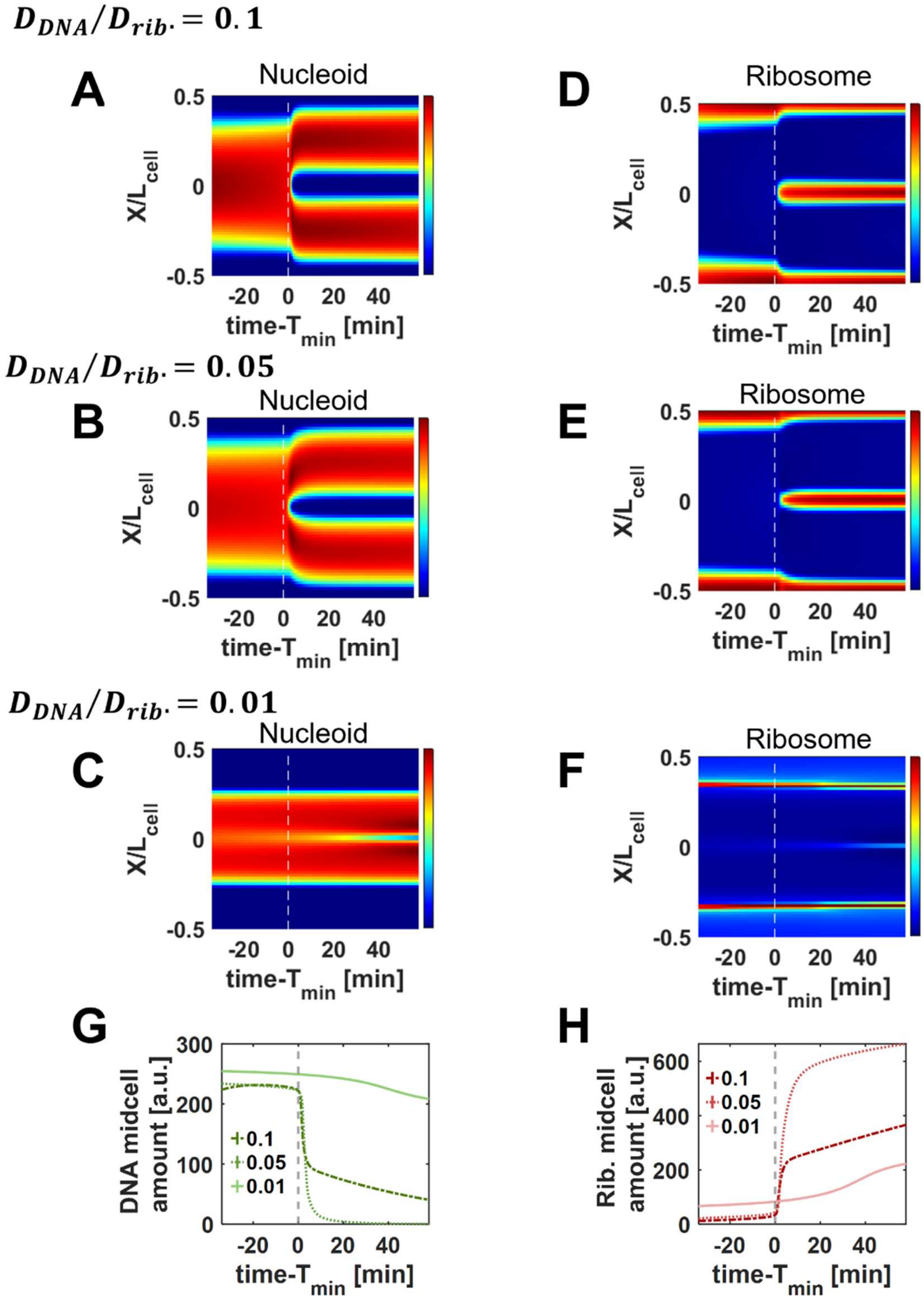


**SI Fig. S15**. The coupled DNA and ribosome dynamics model at different DNA diffusion coefficient, $D_{DNA}$ values. (A-C) Kymographs of DNA density for $D_{\mathrm{DNA}}/D_{\mathrm{rib}} = 0.1,\ 0.05,\ 0.01$, respectively. $D_{\mathrm{DNA}}/D_{\mathrm{rib}} = 0.1$ correspond to the value used in Fig. 3 of the main text. (D-F) The same kymographs for the ribosome density distributions. (G,H) Time evolution of DNA and ribosome midcell amounts for these DNA diffusion coefficients, respectively.

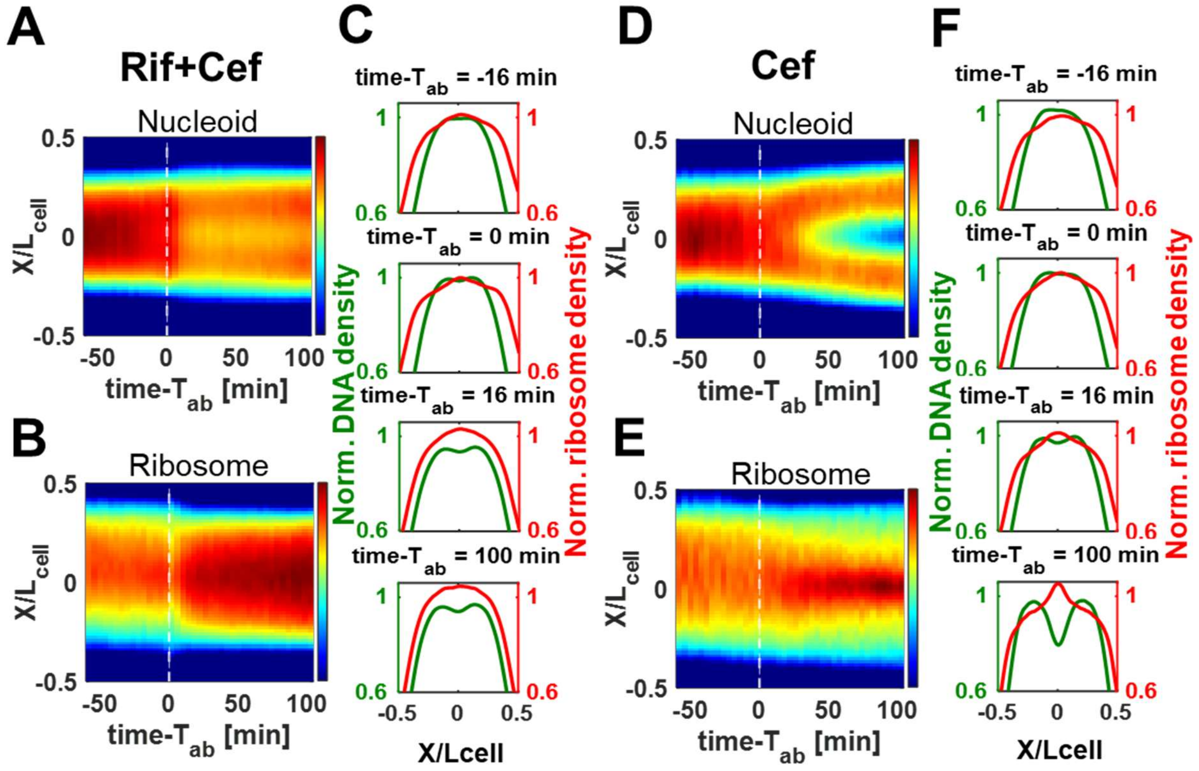


**SI Fig. 16.** The effects from the disruption of active mRNA-polysome dynamics with rifampicin (Rif) on nucleoid and ribosome distributions. (A, D) The population-averaged kymographs of DNA density. The times are relative to the start of antibiotic treatment, $T_{ab}$, with Rif and Cef (cephalexin), and with only Cef, respectively. All analyzed cells in this figure are non-constricting. (B, E) The same for the average ribosome density. (C, F) The density distributions of DNA (green) and ribosome (red) along the long axis of the cell at different time points. In all panels, the curves are normalized relative to their maximum value at $T_{ab}$. $N = 786$ for Rif+Cef and $N = 310$ for only Cef treated cells.

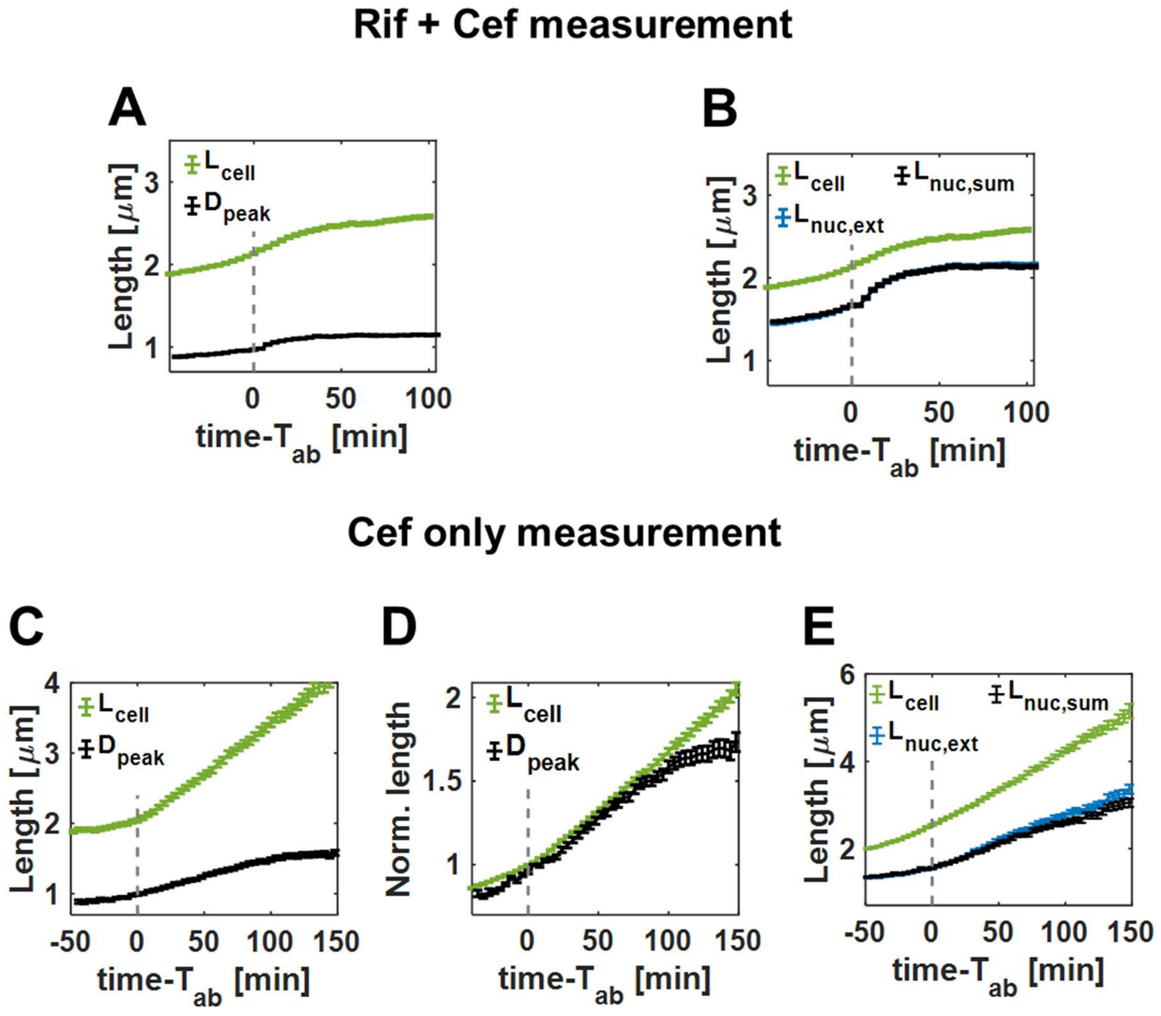


**SI Fig. S17**. The effects of disrupting active mRNA-polysome dynamics by antibiotic treatment in strain CA4 glycerol medium. (A-B) Cells treated with Rifampicin + Cephalexin. Only non-constricting cells were analyzed. (A) Unscaled cell length ($L_{\text{cell}}$) and the distance between the nucleoid peaks, $D_{\text{peak}}$, as a function of time from the beginning of antibiotic treatment, $T_{ab}$. (B) Unscaled cell length $L_{\text{cell}}$, the total nucleoid extent ($L_{\text{nuc,ext}}$), the sum of nucleoid lengths ($L_{\text{nuc,sum}}$) as a function of time. $N = 786$. (C-E) Cells treated with cephalexin only. (C) Unscaled cell length ($L_{\text{cell}}$) and the distance between the nucleoid peaks, $D_{\text{peak}}$, as a function of time from the beginning of antibiotic treatment, $T_{ab}$. (D) Scaled cell length ($L_{\text{cell}}$) and the distance between the nucleoid peaks, $D_{peak}$, as a function of time from the beginning of antibiotic treatment, $T_{ab}$. Both $L_{\text{cell}}$ and $D_{\text{peak}}$ are normalized to 1 at $T_{ab}$. (E) Change of unscaled cell length ($L_{\text{cell}}$), total nucleoid extent ($L_{\text{nuc,ext}}$), sum of nucleoid lengths ($L_{\text{nuc,sum}}$) as a function of time from the beginning of the treatment, $T_{ab}$. $N = 310$. Error bars in all plots are s.e.m.

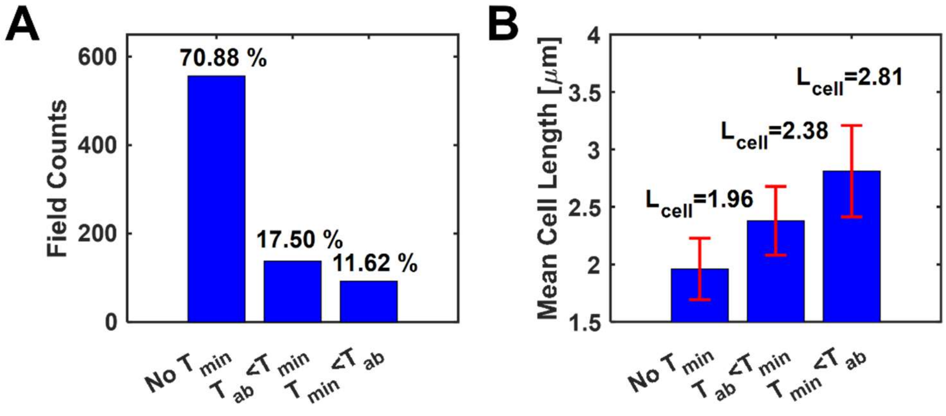


**SI Fig. S18**.Grouping of cells after rifampicin and cephalexin treatment into three categories based on the presence of a DNA density minimum: 1) no local DNA density minimum at midcell (no $T_{\min}$), 2) density minimum appears after the start of treatment ($T_{ab} < T_{\min}$), 3) density minimum present before treatment ($T_{ab} > T_{\min}$). (A) Number of cells in each group with frequencies indicated as percentages. (B) Average cell length of each group at the beginning of the antibiotic treatment. Error bars represent standard deviations.

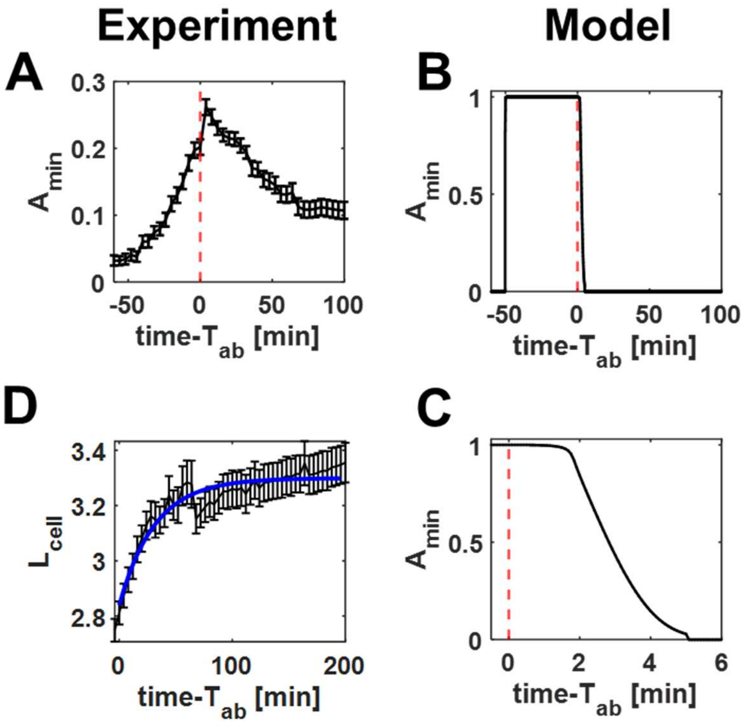


**SI Fig. S19**. Comparing experimental data from $T_{ab} > T_{min}$ group of cells in measurements with rifampicin and cephalexin treatment to the coupled DNA and ribosome dynamics model. The cells in this group have formed a local minimum in nucleoid density before the start of the treatment. (A) The normalized depth of the local DNA density minimum, $A_{min}$, as a function of time from the beginning of the antibiotic treatment. (B) The same curve from the model. (C) The same modeling result zoomed in at the vicinity of $T_{ab}$. (D) The increase in cell length as a function of time from the beginning of the antibiotic treatment. Data are from the experiment (black). The blue curve represents a least-squares fit with the equation: $L_{\text{cell,fit}}(t) = L_0 + (L_\infty - L_0) \times \left(1 - \exp\left[-\frac{t-T_{ab}}{\tau_{\text{rif}}}\right]\right)$. The best-fit values are $L_0 = 2.8$ μm, $L_\infty = 3.3$ μm, $\tau_{\text{rif}} = 31$ min. The model uses $L_{\text{cell,fit}}(t)$ in the calculations shown in panels B and C and a zero rate of polysome assembly. The error bars in panels A and D are s.e.m. $N = 92$ for these two panels.

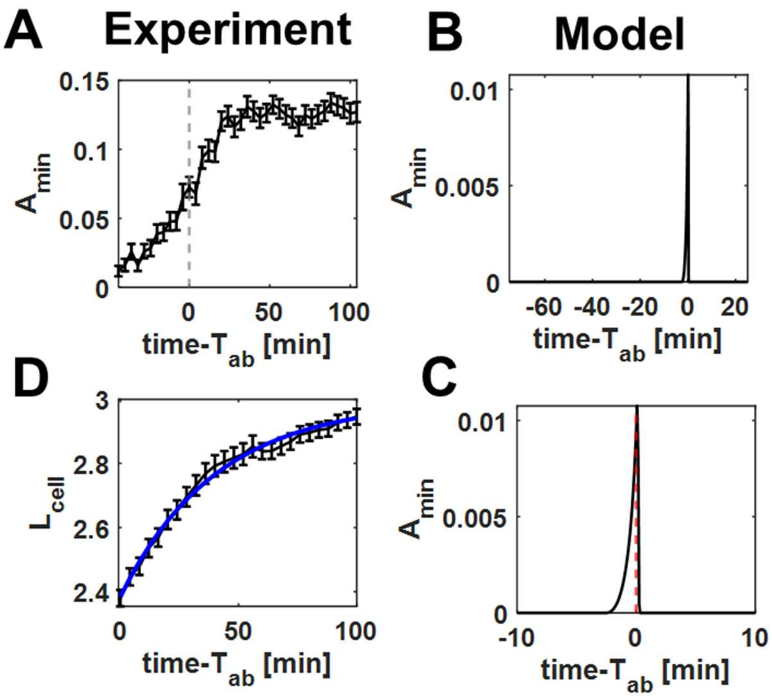


**SI Fig. S20**. Comparing experimental data from $T_{ab} < T_{\min}$ group of cells in measurements with rifampicin and cephalexin treatment to the coupled DNA and ribosome dynamics model. The local minimum in nucleoid density in this group of cells appears after the start of the treatment. (A) The normalized depth of the local DNA density minimum, $A_{\min}$, as a function of time from the beginning of the antibiotic treatment. (B) The same curve from the model. (C) The same modeling result zoomed in at the vicinity of $T_{ab}$. (D) The increase in cell length as a function of time from the beginning of the antibiotic treatment from the experiment (black). The blue curve represents a least-squares fit with the equation: $L_{\text{cell,fit}}(t) = L_0 + (L_\infty - L_0) \times \left(1 - \exp\left[-\frac{t-T_{ab}}{\tau_{\text{rif}}}\right]\right)$. The best-fit values are $L_0 = 2.4$ µm, $L_\infty = 3.0$ µm, $\tau_{\text{rif}} = 41$ min. The model uses $L_{\text{cell,fit}}(t)$ in the calculations shown in panels B and C and a zero rate of polysome assembly. The error bars in panels A and D are s.e.m. $N = 138$ for these two panels.

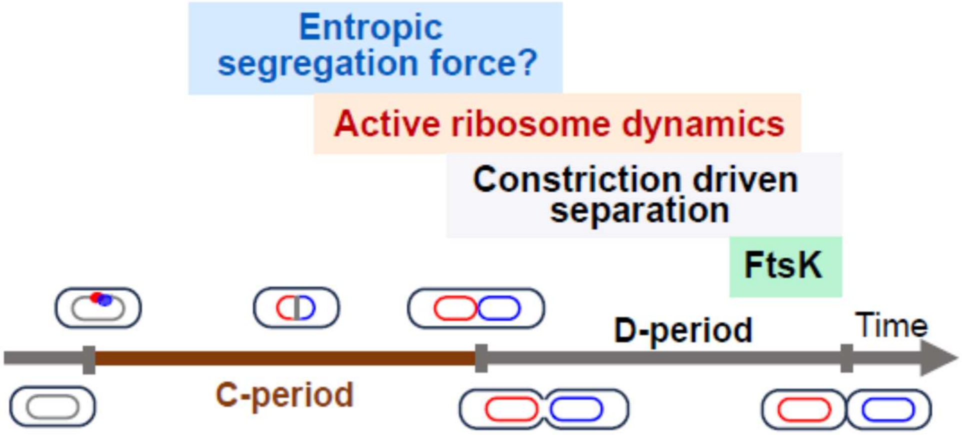


**Figure 21.** Progression of different mechanisms involved in segregation and separation of daughter chromosomes during *E. coli* cell cycle. The schematic summarizes the results from this work and from previously published studies. C-period marks the replication period, and D-period marks the duration from the termination of DNA replication to the final division of two daughter cells. The drawn schematic corresponds to slow growth conditions where the replication initiates after cell birth.

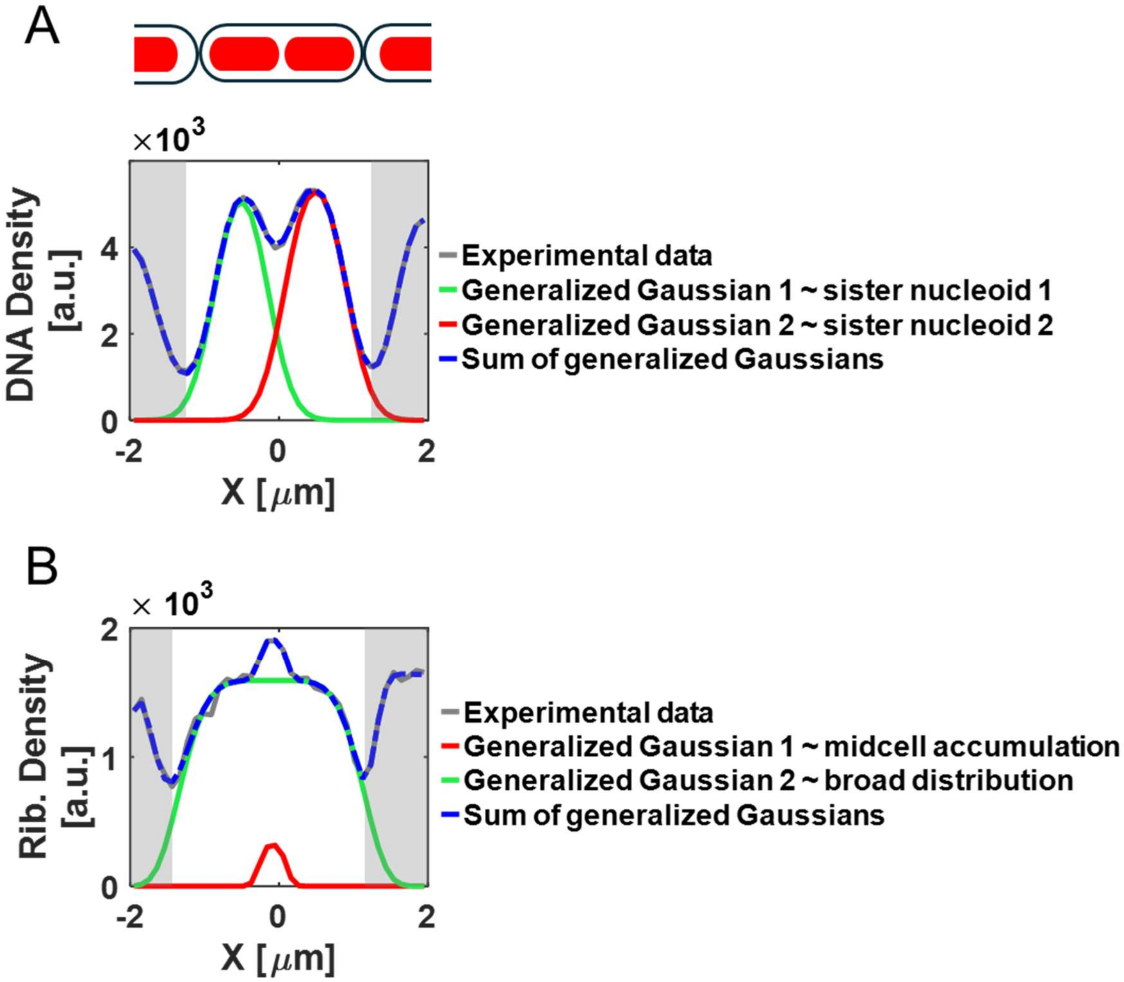


**SI Fig. S22**. Fitting nucleoid and ribosome densities with generalized Gaussian functions. (A) Top: Schematics of cells growing in the mother machine channels. Red blobs indicate nucleoids. The cell of interest and adjacent regions from two neighboring cells are included in the fitting. Bottom: HupA-mYpet signal from a representative cell of strain CA4 grown in M9 glycerol medium (grey, solid) and its fitting with four generalized Gaussians (blue, dash). Two of these generalized Gaussians account for the signal from the neighboring cells (gray shaded areas; not shown separately) and two fit signals from the cell of interest (green and red). (B) The same for the ribosome fluorescent marker RpsB-mCherry (grey, solid). Here too, the fit includes four generalized Gaussians. Two of these generalized Gaussians represent the neighboring cells (gray shaded area) and two the cell of interest (red and green). The broader fitting curve (green) accounts for near uniform distribution of ribosomes along the long axis of the cell and the narrower curve (red) the midcell accumulation.

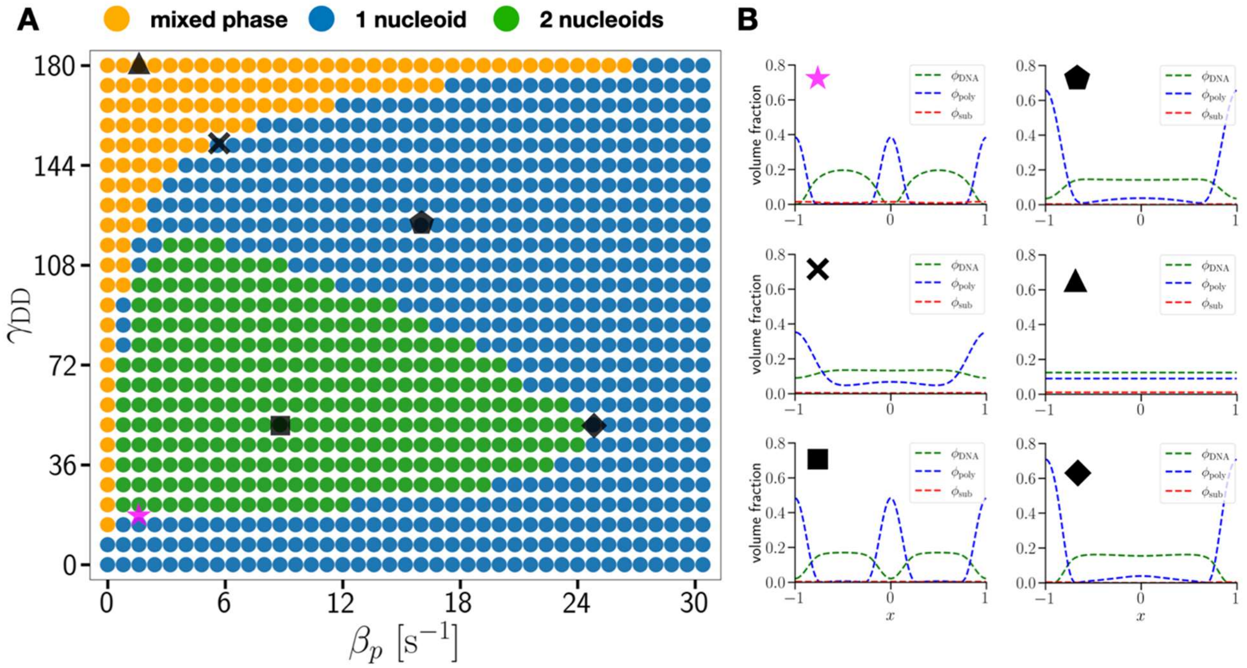


**SI Fig. S23**. Phase diagram for nucleoid formation with respect to $\beta_p$ and $\gamma_{\mathrm{DD}}$. (A) 2D Phase diagram showing the number of nucleoids in steady-state as a function of the polysome production rate $\beta_p$ and DNA self-interaction prefactor $\gamma_{\mathrm{DD}}$. The mixed phase represents a state without apparent phase separation between DNA and crowders. The default parameter set used in this study is indicated by the magenta star ($\beta_p = 1.6$ s$^{-1}$, $\gamma_{\mathrm{DD}} = 17.7$). Black symbols highlight representative parameter sets whose corresponding spatial distributions are shown in panel (B). (B) Steady-state spatial distributions of each species for the parameter sets highlighted in panel (A). Each subplot corresponds to a different parameter set on the phase diagram (indicated by matching symbols), showing volume fraction profiles for DNA, polysomes and subunits.

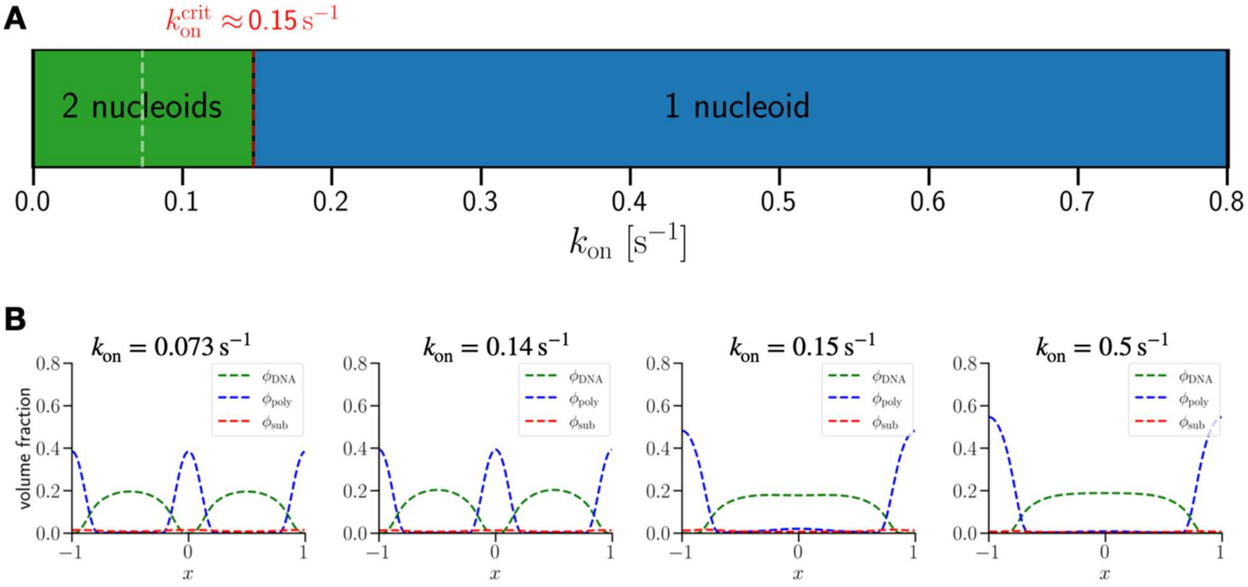


**SI Fig. S24**. Phase diagram for nucleoid formation with respect to $k_{\text{on}}$. (A) 1D Phase diagram showing the number of nucleoids in steady-state as a function of ribosomal subunit binding rate $k_{\text{on}}$. The default value of $k_{\text{on}}$ used in this study is indicated by the white-dashed line ($k_{\text{on}} = 0.073\,\text{s}^{-1}$). The red-dashed line highlights the phase transition boundary between two nucleoids and one nucleoid phases. (B) Representative steady-state spatial distributions of each species for various $k_{\text{on}}$. From left to right, each subplot corresponds to different $k_{\text{on}}$ values (0.073, 0.14, 0.15, and 0.5 s$^{-1}$, respectively.), showing volume fraction profiles for DNA, polysomes, and subunits.

**Table S1: Strains used in the study.**

| Strain | Relevant genetic marker(s) or features | Source, reference or construction |
|---|---|---|
| **BW27783** | *Δ(araD-araB)567, ΔlacZ4787(::rrnB-3), Δ(araH-araF)570(::frt), ΔaraEp-532::frt, hsdR514, φPcp8araE535, rph-1, Δ(rhaD-rhaB)568, λ-* | Yale Coli Genetic Stock Center (CGSC#: 12119) |
| **BW25113** | *Δ(araD-araB)567, ΔlacZ4787(::rrnB-3), hsdR514, rph-1, Δ(rhaD-rhaB)568, λ-* | Yale Coli Genetic Stock Center (CGSC#: 7636) |
| **STK32** | BW27783<br>*hupA::hupA- mCherry-frt*<br>*dnaN::frt-Ypet-dnaN* | *hupA-mCherry:* P1 transduction from PB384, pCP20<br>*Ypet-dnaN:* P1 transduction from RRL190, pCP20 |
| **PB384** | BW25113<br>*hupA::hupA- mCherry-frt* | Mannik et al., 2016 (27) |
| **RRL190** | AB1157<br>*dnaN::frt-aph-frt-Ypet-dnaN* | Reyes-Lamothe et al., 2010 (1) |
| **CA1** | BW25113<br>*rpsB::rpsB-mCherry-frt-aph-frt* | λ-Red recombineering |
| **JM52** | MG1655<br>*hupA::hupA-mYpet-frt-aph-ftr* | λ-Red recombineering |
| **CA4** | BW27783<br>*rpsB::rpsB-mCherry-frt*<br>*hupA::hupA-mYpet-frt* | *rpsB-mCherry:* P1 transduction from CA1, pCP20<br>*hupA-mYpet:* P1 transduction from JM52, pCP20 |
| **JM205** | BW25113<br>*rplI::rplI-mCherry-frt-aph-frt* | λ-Red recombineering |
| **CA7** | BW27783<br>*rplI::rplI-mCherry-frt*<br>*hupA::hupA-mYpet-frt* | *rplI-mCherry:* P1 transduction from JM205, pCP20<br>*hupA-mYpet:* P1 transduction from JM52, pCP20 |

**Table S2. List of oligos used in the study.**

| Name | Sequence (5'->3') | Notes |
|---|---|---|
| rpsB-F_LR | CCCAGGCGGAAGAAAGCTTCGTAGAAGCTG AGCAGGAAAGGCGACAGGAGGTGAGCAAGG GCGAGGAG | λ-Red > *rpsB-mCherry* |
| rpsB-R_LR | ACTCGAACTATTTTGGGGGAGTTATCAAGCCA TATGAATATCCTCCTTAGA | λ-Red > *rpsB-mCherry* |
| rplI-F-LR | CAGCGAAGTATTCGCGAAAGTGATCGTAAAC GTAGTAGCTGAAGTGAGCAAGGGCGAG | λ-Red > *rplI-mCherry* |
| rplI-R-LR | ACCAATGGTCGGCGTTTTTACGTCTCGTTGAA TAACGAACATATGAATATCCTCCTTAGA | λ-Red > *rplI-mCherry* |
| hupA-F | GCTAACGTACCGGCATTTGTTTCTGGCAAGG CACTGAAAGACGCAGTTAAGAGCTCGGCTGG CTCCGCTG | λ-Red > *hupA-mYpet* |
| hupA-R | AAGGGGTGAAACCACCCCTTCGTTAAAACTG TTCACTGCCACGCAATCTTACATATGAATATCC TCCTTAG | λ-Red > *hupA-mYpet* |
| rpsB-F-test | GTAACGACGACGCAATCCGTG | verification of *rpsB-mCherry* |
| rpsB-R-test | CCTTTCTGCAACTCGAAC | verification of *rpsB-mCherry* |
| rplI-test-F | CGGTCACGGTCCATTAATAC | verification of *rplI-mCherry* |
| rplI-test-R | GGTGAATGGATCTACTTGCG | verification of *rplI-mCherry* |
| hupA-check-F | CGCAGACCGGTAAAGAAATC | verification of hupA-mYpet |
| hupA-check-R | CAGCATCAATGATCGACGC | verification of *hupA-mYpet* |

**Table S3. The linear fit parameters for data in Fig. 3.** The fitting lines are defined by $y = a(\text{time} - T_{\min}) + b$

| Quantity | Panel | $a$ [$\text{min}^{-1}$] | $b$ | $R$ |
|---|---|---|---|---|
| Norm. midcell amount, exp. | I | $-1.4 \cdot 10^{-3}$ | 0.92 | $-0.97$ |
| Norm. midcell amount, model. | I | $-1.9 \cdot 10^{-3}$ | 0.43 | $-0.99$ |

**Table S4: List of parameters obtained from fitting of exponential cell length growth after Rif+Cef treatment using the equation:**

$$L_{cell,fit}(t) = L_0 + (L_\infty - L_0) \times \left(1 - \exp\left[-\frac{t - T_{ab}}{\tau_{rif}}\right]\right)$$

| Group | $L_0$ [μm] | $L_\infty$ [μm] | $\tau_{rif}$ [min] |
|---|---|---|---|
| All Cells | 2.32 | 2.87 | 45 |
| No $T_{min}$ | 2.14 | 2.62 | 43 |
| $T_{min} > T_{ab}$ | 2.38 | 2.99 | 41 |
| $T_{min} < T_{ab}$ | 2.84 | 3.3 | 31 |

**Table S5: Definitions and values of individual parameters in the coupled DNA and ribosome dynamics model**

| Parameter | Value | Parameter | Value |
|---|---|---|---|
| $g$ | 145 (13) | $\alpha$ | 0.85 (13) |
| $\gamma$ | 1.309 (13) | $\phi_{\mathrm{poly}}^{\mathrm{ov}}$ | 1.86 (13) |
| $v_{\mathrm{sub}}$ | $4.2 \times 10^{-6} \mu\mathrm{m}^3$ | $v_{\mathrm{poly}}$ | $1.8 \times 10^{-4} \mu\mathrm{m}^3$ |
| $v_{\mathrm{DNA}}$ | $6.65 \times 10^{-5} \mu\mathrm{m}^3$ (Eq.7) | $V_{\mathrm{DNA}}$ | $0.1\ \mu\mathrm{m}^3$ |
| $V_{\mathrm{DNA,0}}$ | $27\ \mu\mathrm{m}^3$ (17) | $B_{\mathrm{DNA\text{-}poly}}$ | $8.47\ \mu\mathrm{m}^3$ (Eq. 8) |
| $B_{\mathrm{poly\text{-}sub}}$ | $3.35 \times 10^{-4}\ \mu\mathrm{m}^3$ (Eq. 15) | $B_{\mathrm{DNA\text{-}sub}}$ | $0.6\ \mu\mathrm{m}^3$ (Eq. 9) |
| $D_{\mathrm{poly}}$ | $0.01\ \mu\mathrm{m}^2/\mathrm{s}$ (4) | $D_{\mathrm{DNA}}$ | $0.001\ \mu\mathrm{m}^2/\mathrm{s}$ |
| $D_{\mathrm{sub}}$ | $0.035\ \mu\mathrm{m}^2/\mathrm{s}$ | $\alpha_1$ | 1 |
| $\alpha_2$ | 200 | $\kappa_{DD}$ | $0.5\ \mu\mathrm{m}^2$ |
| $\kappa_{rr}$ | $0.1\ \mu\mathrm{m}^2$ | $\kappa_{Dr}$ | $0.05\ \mu\mathrm{m}^2$ |
| $\kappa_{rD}$ | $0.005\ \mu\mathrm{m}^2$ | $\beta_p$ | see Eq. (29) |
| $\beta_d$ | $0.003\ \mathrm{s}^{-1}$ (21) | $k_{\mathrm{on}}$ | $0.073\ \mathrm{s}^{-1}$ |
| $k_{\mathrm{off}}$ | $0.0058\ \mathrm{s}^{-1}$ (22) | $N_{\mathrm{ribo}}$ | 10 (13,14) |
| $n_{\mathrm{poly}}$ | 600 (13) | $V_{\mathrm{cell}}$ | $0.8\ \mu\mathrm{m}^3$ (28) |
| $L_0$ | $2.0\ \mu\mathrm{m}$ | $L_i$ | $1.3\ \mu\mathrm{m}$ |
| $\alpha_g$ | 0.00020 | $\lambda_g$ | $0.020\ \mu\mathrm{m}$ |

**Table S6: Definitions and values of parameters derived from dimensionless combinations in the coupled DNA and ribosome dynamics model**

| Parameter | Expression | Value |
|---|---|---|
| $\gamma_{rr}$ | $\frac{\gamma+1}{\left(0.69\phi_{\text{poly}}^{\text{ov}}\right)^{\gamma}}$ | 1.67 |
| $\gamma_{rD}$ | $\frac{B_{\text{DNA-poly}}}{V_{\text{DNA}}}$ | 84.7 |
| $\gamma_{rs}$ | $\frac{B_{\text{poly-sub}}}{v_{\text{sub}}}$ | 81.2 |
| $\gamma_{sD}$ | $\frac{B_{\text{DNA-sub}}}{V_{\text{DNA}}}$ | 6 |
| $\gamma_{DD}$ | $\frac{g\alpha(\alpha+1)v_{\text{DNA}}V_{\text{DNA,0}}^{\alpha}}{V_{\text{DNA}}^{\alpha+1}}$ | 17.7 |
| $\gamma_{Dr}$ | $\frac{v_{\text{DNA}}B_{\text{DNA-poly}}}{v_{\text{poly}}V_{\text{DNA}}}$ | 31.4 |
| $\gamma_{Ds}$ | $\frac{v_{\text{DNA}}B_{\text{DNA-sub}}}{v_{\text{sub}}V_{\text{DNA}}}$ | 95 |
| $\gamma_{pp}$ | $\frac{N_{\text{ribo}}v_{\text{sub}}}{v_{\text{poly}}}$ | 0.233 |

### Legends for Movies

#### Movie S1

Time-lapse video showing replisome and nucleoid dynamics during steady-state growth of *E. coli* cells (strain STK32) in a microfluidic mother machine channel. Phase contrast images and fluorescence signals for Ypet-DnaN (replisome marker) and HupA-mCherry (nucleoid marker) are shown. Cells were grown in M9 glycerol + trace elements medium. The scale bar corresponds to 5 μm.

#### Movie S2

Time-lapse video showing replisome and nucleoid dynamics in a single *E. coli* cell (strain STK32) under steady-state growth conditions. Phase contrast images and fluorescence signals for Ypet-DnaN (replisome marker) and HupA-mCherry (nucleoid marker) are shown. The cell was grown in M9 glycerol + trace elements medium in a microfluidic mother machine channel. Time stamps represent time from cell birth. The scale bar corresponds to 1 μm.

#### Movie S3

Time-lapse video showing ribosome and nucleoid dynamics during steady-state growth of *E. coli* cells (strain CA4) in a microfluidic channel. Phase contrast images and fluorescence signals for HupA-mYpet (nucleoid marker) and RpsB-mCherry (ribosome marker) are shown. Cells were grown in M9 glycerol + trace elements medium. The scale bar corresponds to 5 μm.

#### Movie S4

Time-lapse video showing ribosome and nucleoid dynamics in a single *E. coli* cell (strain CA4) during steady-state growth. Phase contrast images and fluorescence signals for HupA-mYpet (nucleoid marker) and RpsB-mCherry (ribosome marker) are shown. The cell was grown in M9 glycerol + trace elements medium in a mother machine microfluidic channel. Time stamps indicate time from cell birth. The scale bar corresponds to 1 μm.
